# Seizure-related gene 6 (SEZ6) encodes a Cancer stem cell-specific proangiogenic molecule: A novel glioma therapeutic target

**DOI:** 10.64898/2026.07.15.738615

**Authors:** Arani Mukherjee, Shreoshi Sengupta, Abhishek Chowdhury, Lakshay Garg, Kaval Prasasvi Reddy, Parimal Mishra, Dhrubjyoti Sharma, Subhrodeep Saha, Jayanta Chatterjee, Parichay J. Perikal, Sunil Valentine Furtado, Vijay Thiruvenkadam, Sivapriya Kirubakaran, Kumaravel Somasundaram

## Abstract

Cancer stem-like cells (CSCs) not only initiate tumors but also orchestrate angiogenesis through multiple mechanisms, thereby sustaining vascularization and promoting tumor growth. In glioblastoma (GBM; WHO grade IV glioma), the most aggressive adult brain tumor, glioma stem-like cells (GSCs; glioma CSCs) are the main drivers of tumor progression, drug resistance, and recurrence. An interrogated multi-omic secretome analysis identified Wnt-β-catenin signaling-regulated, membrane-localized Seizure Related 6 Homolog (SEZ6) as a patient-derived GSC-specific proangiogenic molecule. Silencing of SEZ6 inhibited the ability of the GSC secretome to induce angiogenic network formation by brain- and lung-derived endothelial cells, but not GSC growth as neurospheres *in vitro*. SEZ6 silencing also suppressed patient-derived GSC-initiated glioma tumors, resulting in reduced tumor vasculature in an orthotopic mouse model. We also found that SEZ6 induces TGFβ-dependent IL-8 expression in endothelial cells to promote angiogenesis. Coimmunoprecipitation and molecular dynamics simulation experiments determined that the Sushi 3 domain of SEZ6 mediates the interaction with TGFβ RII to activate the TGFβ pathway. Pharmacological inhibition of BACE1 with an in-house–developed small molecule, or blockade of the SEZ6–TGFβ RII interaction using a rationally designed peptide, markedly attenuated SEZ6-driven TGFβ signaling, and suppressed angiogenesis. Thus, our findings identify SEZ6 as a novel CSC-specific therapeutic target for GBM.

## Introduction

While genetic and epigenetic alterations drive tumor initiation and support early stages of malignant growth, the sustained progression and aggressiveness of tumors are critically dependent on angiogenesis, which provides the expanding tumor mass with a continuous supply of oxygen, nutrients, and metabolic support. Beyond sustaining growth, tumor-induced angiogenesis also contributes to invasion, therapeutic resistance, and disease recurrence. Consequently, identifying and targeting key molecular drivers of pathological angiogenesis has become an attractive and strategically important approach for developing novel anticancer therapies. However, despite a strong biological rationale, clinical translation of anti-angiogenic strategies has met with limited success. Notably, bevacizumab (Avastin), a humanized monoclonal antibody targeting VEGF-A, did not significantly improve patient outcomes in Phase III clinical trials (**Chino et al., 2014; Gilbert et al., 2014**). This failure highlights the complexity and redundancy of angiogenic signaling networks in tumors and underscores the pressing need to identify alternative pro-angiogenic targets to enable more effective and durable therapeutic interventions.

Glioblastoma (GBM), a grade IV glioma, characterizes the most aggressive primary brain tumor in adults and is associated with an extremely poor prognosis, with median overall survival remaining limited to approximately 12–18 months despite aggressive standard-of-care treatment (**Stupp et al., 2009**). Evidence has highlighted the key role of cancer stem cells (CSCs)—a rare but highly specialized subpopulation within tumors—which possess the unique capacity to initiate tumor formation, sustain long-term growth, and evade conventional therapeutic modalities (**Visvanathan and Somasundaram, 2017**). Glioma stem cells (GSCs), the CSC counterparts in GBM, are now recognized as the principal drivers of the remarkable aggressiveness, therapeutic resistance, and inevitable recurrence of GBMs (**Visvanathan et al., 2018**). Their resistance to radiotherapy and temozolomide-based chemotherapy underscores the urgent need to identify molecular vulnerabilities within this therapy-refractory GSC compartment and exploit them to develop effective adjuvant strategies that enhance the efficacy of existing GBM treatments. Importantly, GSCs preferentially localize to the perivascular niche, positioning them as critical modulators of tumor angiogenesis (**Prager et al., 2020; Calabrese et al., 2007; Sengupta et al., 2022).** Together, these observations underscore the importance of identifying GSC-specific pro-angiogenic targets as a prerequisite for developing more effective and durable therapeutic interventions for GBM.

An integrated analysis combining the secreted proteome and transcriptome of matched patient-derived GBM organoids enriched for glioma stem cells (GSCs) and their differentiated counterparts (DGCs) revealed a distinct set of molecules selectively expressed by GSCs. Subsequent functional screening identified seizure-related 6 homolog (SEZ6) as a particularly potent promoter of angiogenesis, highly expressed in GSCs, yet not influencing GSC proliferation in vitro. SEZ6 is a transmembrane protein that undergoes proteolytic cleavage by the β-site amyloid precursor protein–cleaving enzyme 1 (BACE1) (**Pigoni et al., 2016**), resulting in the release of a soluble ectodomain that acts on endothelial cells to drive angiogenic responses. We further identified the cognate endothelial cell-surface receptor mediating SEZ6 signaling and established interleukin-8 (IL8) as the principal downstream effector of SEZ6-induced angiogenesis. To therapeutically target this pathway, we developed two complementary strategies: a novel small-molecule inhibitor of BACE1 that suppresses SEZ6 shedding, and a de novo–designed peptide that disrupts the interaction between SEZ6 and its endothelial receptor. Both approaches effectively abrogated endothelial signaling and angiogenesis. Collectively, these findings establish SEZ6 as a GSC-specific pro-angiogenic molecule and provide proof of principle that targeting SEZ6 is a promising, previously unexplored therapeutic strategy for glioblastoma.

## Materials and Methods

### Cell lines

The GBM adherent cell lines LN229 and U251 were purchased from Sigma-Aldrich (Saint Louis, Missouri, USA). DBT-Luc cells were a kind gift from Dr. Dinesh Thotala at Washington University in St. Louis, St. Louis, Missouri, United States. ST1 endothelial cells were a kind gift from Dr. Ron Unger, Johannes Gutenberg University, Germany. B.End3 cell line purchased from The American Type Culture Collection (ATCC, #CRL-2299). The HBMECs were purchased from Cell Biologics, USA (#H-6023). Glioma cell lines (LN229) and HEK-293T were grown in Dulbecco’s Modified Eagle’s Medium (DMEM) (Sigma-Aldrich, #D5648) supplemented with 10% Fetal Bovine Serum (FBS), Penicillin, and Streptomycin.

### Plasmids used

shRNA constructs for all the genes mentioned in the study were obtained from the TRC (The RNAi Consortium) shRNA library. The SEZ6 overexpression plasmid was purchased from Origene (USA; #RC212279). SEZ6 Promoter Luc (SPL) was bought from Genecopoeia, USA (#HPRM33968)

### GBM sphere culture

Primary human tumor-derived GSCs MGG4, MGG6, and MGG8 were kindly gifted by Dr. Wakimoto (Massachusetts General Hospital, Boston, USA). Spheres are collected, spun down, washed with PBS, and chemically dissociated using NeuroCult™ Chemical Dissociation Kit (Mouse) from Stem Cell Technologies, per the manufacturer’s instructions. Single cells are counted and 30000-500000 cells are plated per 6 well in Neurobasal Medium (Invitrogen) containing the necessary supplements-N2 (final conc. 0.25X, Invitorgen), B27 (final conc. 1X), Heparin (Sigma, final conc. 2 μg/ml), Glutamine (Invitrogen, final conc. 3mM), FGF2 (Peprotech, final conc. 20ng/ml), EGF (R and D, final conc. 20ng/ml) and 1X antibiotics (as used for DMEM media for monolayer culture).

### GSC differentiation

Fully grown spheres were collected on the 7th day of culture, washed with PBS, and re-plated without much agitation into standard attachment cell culture dishes in complete DMEM medium (Sigma) supplemented with 10% FBS and antibiotics. The cells were allowed to differentiate for 15 days and then trypsinized using 1X Trypsin-EDTA solution for further studies.

### ST1 endothelial cell culture

ST1 cells, which are immortalized pulmonary microvascular endothelial cells, received as a generous gift from Dr. Ron Unger, University of Mainz, were grown in Medium 199 (Sigma) supplemented with Heparin, Glutamine, Endothelial cell growth supplements (Sigma), and 20% FBS.

### Neurosphere assay

Viral infection was performed in GSCs using a lentivirus encoding non-targeting shRNA (shNT) or shRNA targeting the gene of interest. The small pellets of cells were collected 48 hrs after viral infection, dissociated to form single cells that were counted and re-plated in 6-well plates (3×10^4^ cells/well) in complete Neurobasal medium. At the same time, cells were harvested and checked for specific gene manipulation (like knockdown verification). The medium was replenished every 2-3 days, and sphere formation was monitored until the 6th or 7th day after re-plating. The number of spheres, sphere diameter, and size were analyzed using Image J software. Spheres having area less than 50 μm^2^ were excluded from the analysis.

### Limiting Dilution assay

For each condition, 1, 10, 50, 100, and 200 GSCs (single cells) were plated in 10 wells each, respectively, of a 96-well plate, and sphere formation was assessed over the next 5-7days. The number of wells not forming spheres were counted and plotted against the number of cells per well. Extreme Limiting Dilution Assay using the ELDA software available online (https://www.elda.at/cdscontent/?contentid=10007.854970&portal=eldaportal).

### Secretome collection and processing

Spheres were grown in complete Neural Stem Cell medium containing Glutamine, Heparin, N2, B27, EGF, and FGF for 6 days, then thoroughly washed with PBS and re-plated without disrupting the spheres in incomplete Neural Stem Cell medium with no added supplements or growth factors. 36 hours after plating in incomplete medium, the conditioned medium was collected, centrifuged at 1500 rpm for 10 min at 4°C, filtered through 0.4 µm syringe filters, and stored at -80°C. The differentiated cells were grown in complete DMEM with 10% FBS, washed well with PBS, and the secretome was collected after 36 hours in incomplete medium and stored under similar conditions.

### Lentivirus preparation and transduction of cells

HEK-293T cells were seeded in 60 mm or 90 mm Poly-L-Lysine-coated cell culture dishes. The cells were transfected with shRNA plasmid and helper plasmids psPAX2 and pMD2.G using Opti-MEM (Invitrogen #22600-050) medium and lipofectamine (https://www.thermofisher.com/order/catalog/product/12566014 Invitrogen) when the cells were 60-70% confluent. Six hours after transfection, the Opti-MEM medium was replaced with fresh DMEM medium supplemented with 10% FBS. Sixty hours post-transfection, the supernatant from the transfected cells was collected in 15 ml Falcon tubes, centrifuged at 5000 rpm for 10 min, filtered through a 0.45 μm filter, and stored at -80°C for future use. For lentivirus used in endothelial cells, DMEM was removed after 24 hrs. of transfection and replaced with complete M199, and the lentivirus was collected after 60 hrs., as mentioned previously. The same method was used to collect lentiviruses for GSCs (DMEM was replaced with NBM).

### Screening for the effect of knockdown of the TCF/LEF1 target genes

To test the effect of the individual TCF/LEF1 target gene silencing on neurosphere formation, 5 × 10^4^ MGG8-GSCs were plated in each well of ultra-low attachment plates from Corning. The cells were plated in 1ml of lentivirus for each condition, along with polybrene. After 48 hrs of infection, the cells were harvested, counted, and 3 × 10^4^ cells were plated in each well of the 6-well plate. The neurosphere formation was monitored till the 6^th^ day after re-plating. Images of the spheres were taken for quantification on the 6^th^ day.

To assess the impact of gene silencing on the ability of CMs from MGG8 to modulate network formation by endothelial cells, CMs were collected from shRNA-mediated gene-silenced neurospheres that were plated for neurosphere growth. After the neurosphere formation assay was performed, the spheres were replated in incomplete NBM, kept for 36 hrs, and the conditioned media were collected 36 hrs. later. This medium was concentrated, and 50 μg of protein was used from each condition to assess the effect on endothelial cell network formation.

### Angiogenesis proteome profiler array

CMs from MGG8/shNT and MGG8/shSEZ6 were collected as described below. 100 μg of proteins were added from each condition to ST1 cells plated in incomplete Medium 199. The ST1 cells were kept in the CM for 24 hrs. and were harvested. The cell pellets were lysed using RIPA lysis buffer (composition mentioned below), 150 μg of proteins were taken from each condition and were mixed with the array buffer provided with the kit (Proteome Profiler Array, Human Angiogenesis Array Kit, catalog number AY007, R and D Systems, USA), and then incubated with the detection antibody cocktail mix for 1 hr. In the meantime, the membranes, which came pre-spotted with primary antibodies for 55 angiogenesis-related molecules, were blocked using the buffer provided with the kit. After pre-incubation, the lysate-detection antibody cocktail mix was added to the membranes and incubated overnight at 4 °C. The following day, the membranes were washed, incubated with Streptavidin-HRP, and developed with a chemiluminescent reagent on a ChemiDoc. The spot intensities were measured with an Alpha Digi-Doc, and the mean pixel densities were normalized to the reference spots.

### ELISA

For ELISA, the ST1 endothelial cells were treated with 50 μg of proteins from MGG8/shNT and MGG8/shSEZ6 conditioned medium, respectively, for 24 hours. To detect intracellular protein levels, treated ST1 cells were pelleted, lysed in RIPA lysis buffer, and 100 μg of protein from each condition was analyzed by ELISA. To study the secretome, cells were transferred to incomplete Medium 199 after CM treatment and collected after 36 hours. The collected CM was concentrated using Amicon Ultra centrifugal units (from 10 mL to 500 μL) and used (100 μg equivalent) for the ELISA for each condition. For IL8 detection, the DuoSet ELISA development kit from R&D Systems was used.

### Boyden chamber assay for cell migration

Transwell assay was performed in 24-well Boyden chambers with 8-μm-pore-size polycarbonate membranes (BD Biosciences, San Diego, USA). ST1 cells (5 × 10^4^) were re-suspended in 500 μl serum-free Medium 199 and placed in the upper chamber, and the lower chamber was filled with serum-free Medium 199 with 750 μl conditioned medium or 400 nM final concentration of recombinant protein dissolved in incomplete medium (serving as a chemo-attractant). After 24 hrs of incubation, cells remaining on the membrane’s upper surface were removed with a wet cotton bud. The cells that had migrated to the lower surface of the membrane were fixed in ice-cold methanol, stained with crystal violet, and imaged with a light microscope.

### Boyden chamber assay for cell invasion

Transwell assay was performed in 24-well Boyden chambers with polycarbonate membranes (8 μm pore size) coated with Matrigel (BD Biosciences, San Diego, USA). ST1 cells (5 × 104) were re-suspended in 500 μl serum-free Medium 199 and placed in the upper chamber, and the lower chamber was filled with serum-free Medium 199 with 750 μl conditioned medium or 400 nM final concentration of recombinant protein dissolved in an incomplete medium (serving as a chemo-attractant). After 24 hrs. of incubation, cells remaining on the membrane’s upper surface were removed with a wet cotton bud. The cells that had invaded the lower surface of the membrane were fixed in ice-cold methanol, stained with crystal violet, and imaged under a light microscope.

### Cell proliferation assay (MTT assay)

ST1 cells (1.5 × 10^3^) were plated in 2% FBS-containing M199 Medium in each well of a 96-well plate. MTT assay was performed as per the established protocol. MTT was added to each well, and Formazan crystals formed after 3 hrs. of incubation were dissolved in DMSO, and the absorbance was measured at 420 nm. The first reading served as the untreated condition (0^th^ time-point). After this reading, the cells were treated with 400 nM rhSEZ6 or an equivalent concentration of BSA, and readings were taken every 24 hrs., till 96 hrs. The cell viability was then plotted as a line graph.

### In-vitro angiogenesis assay (tube formation assay)

15,000 endothelial cells (ST1 cells) in Medium 199 (Sigma) without growth factors were seeded per well in a 96-well plate coated with Geltrex (Invitrogen) and incubated for 1 hour. Serum-free conditioned media from different conditions were added on top of the cells, and an overnight incubation was done. After 10-12 hours of incubation, endothelial cells form tube-like structures. Each complete circular structure was treated as a single network, and the total number of networks for each condition was plotted. For BSA (Sigma-Aldrich, A9647-100G), 10 mg/ml stock solutions were prepared by dissolving the lyophilized protein in sterile water, after which appropriate dilutions were prepared to achieve the working concentration. 100 ng each of rhSE6 and rhIL8 were used for the assays.

### Generation of deletion mutants of SEZ6

Deletion constructs were generated using Q5® Site-Directed Mutagenesis Kit (NEB, #E0554S), according to the manufacturer’s instructions. Domain-specific deletion primers were generated using the NEBaseChanger® primer designing tool. The SEZ6 overexpression construct was used as a template for amplifying the domain-specific deletion constructs. The clones thus obtained were confirmed using sequencing and used for further experimentation.

### RNA isolation, cDNA synthesis, and RT-qPCR

Total RNA was isolated from the shRNA transduced cells using the TRI reagent (Sigma) as per the manufacturer’s instructions. RNA integrity was assessed by running it on a 2% MOPS formaldehyde gel. They were quantified by NanoDrop. 2 μg of RNA was used for cDNA synthesis using the cDNA synthesis kit (Applied Biosystems).

### Quantitative real-time PCR (qPCR)

RNA was isolated using TRI reagent (Sigma-Aldrich, #T9424) as per the manufacturer’s protocol. 2 μg of RNA was used to prepare cDNA using the high-capacity cDNA conversion kit (Life Technologies, #4368813). The cDNA thus generated was diluted with nuclease-free water to a final concentration of 10 ng/μl. Subsequently, qRT-PCR was done using the ABI PRISM 7900 HT Sequence Detection System (Life technologies, USA) with the cDNA as the template and DyNAmo Flash SYBR Green qPCR kit (Applied biosystems, #F-416L), using gene-specific primer sets under default conditions 95◦C for 15 min, 40 cycles of 95°C for 20 seconds, 60°C for 25 seconds and 72°C for 30 seconds followed by the dissociation cycle. Each sample was run either in duplicate or in triplicate. ATP5G was used as an internal control, and quantification of genes was performed by the ΔΔct method.

### Western Blot

Cells were harvested and centrifuged at 3000 rpm for 5 minutes to obtain the protein pellet. The pellet was lysed using RIPA (Radio Immuno Precipitation Assay) buffer. For cell lysis, the solution was vortexed and incubated on ice for 30 minutes. After incubation, the solution was centrifuged at 14000 rpm for 30 minutes at 4°C. The protein was isolated by collecting the supernatant in a fresh tube, without disturbing the pellet. Protein estimation was done using Bradford Assay Reagent. The required amount of protein was mixed with the 6X protein loading dye containing BPB (bromophenol blue), SDS (sodium dodecyl sulphate), and β-mercaptoethanol, and was subjected to denaturation of proteins for 15 minutes at 95°C. Proteins were separated by carrying out 12-14% SDS-PAGE (Sodium Dodecyl Sulfate-Polyacrylamide Gel Electrophoresis). The separated proteins were transferred onto polyvinylidene difluoride (PVDF) membrane by a semi-dry transfer method. The membrane was then blocked by incubating it in 5% non-fat dried skimmed milk prepared in TBST (20 mM Tris-HCl, pH 7.4, 137 mM NaCl, 0.1% Tween 20) for 45 minutes to prevent non-specific antibody binding. The blot was incubated with primary antibodies prepared in 5% BSA, overnight (12-14 hours). The membrane was washed three times with TBST, for 10 minutes each, at high rocker speed. Then the membrane was incubated with anti-rabbit and anti-mouse horse-radish peroxidase (HRP)-conjugated secondary antibodies (1:5000 dilution) prepared in 5% skimmed milk in TBST for 2-3 hours at room temperature at very slow rocker speed. The membrane was washed three times with TBST, for 10 minutes each, at high rocker speed. The blots were developed using Clarity ECL Western Blotting Substrate (BioRad, catalog #170-5061) in ChemiDoc™ Imaging System (#12003153) from BioRad. The following antibodies were used: Anti-SEZ6 (1:1000 dilution), Anti-β-Catenin (1:1000 dilution), Anti-SOX2 (1:1000 dilution), Anti-DDK (1:3000 dilution), Anti-BACE1 (1:1000 dilution), Anti-TGFβ RII (1:4000 dilution), Anti-Na-K-ATPase (1:40,000 dilution), Anti-β-ACTIN (1:20,000 dilution), Anti-GAPDH (1:20,000 dilution).

### Luciferase assay

For the luciferase assay, MGG8-GSCs or HEK-293T cells were plated in 6-well plates. After 18 hours of seeding, cells were co-transfected with Luciferase reporter constructs and vector control or overexpression constructs. After 24 or 72 hours of transfection, cells were harvested and lysed in 1X reporter lysis buffer (Promega, #E3971). An equal amount of protein was used for the luciferase assay utilizing luciferase assay reagent (Promega, #E1483) using a luminometer (Berthold detection system, Sirius). The Student’s t-test was used to calculate the statistical significance.

### Immunoprecipitation

Immunoprecipitation experiments were performed following the standard protocols. Cells were scraped from the culture dish in Pierce IP lysis buffer (25 mM Tris-HCl pH 7.4, 150 mM NaCl, 1 mM EDTA, 1% NP-40 and 5% glycerol, Sodium orthovanadate (1mM), Sodium Fluoride (1mM), Phenyl methyl sulfonate (1 mM), Protease inhibitor cocktail (Sigma, #S8830)) and kept for lysis at 4°C for 30 minutes with agitation. Debris was cleared by centrifugation of the lysate at 14000 rpm for 30 minutes at 4°C. For each IP preparation. 500 µg of protein lysate was incubated with 2.5 μg of antibody or control IgG antibody for 18-20 hours at 4°C. 30 μL of protein G-conjugated magnetic beads (Invitrogen, catalog #10004D) were then added to each set and incubated for 4 hours at 4°C to capture the immune complexes. Beads were washed thrice with ice-cold Pierce IP lysis buffer by gentle rotation at 4°C for 5 minutes. The immunoprecipitated proteins were denatured at 95°C for 15 min in 6X Laemmli buffer and then resolved by 10% SDS-PAGE followed by transfer to PVDF membrane (MERCK, Cat #IPVH00010). The membrane was blocked by TBST buffer containing 5% non-fat dried milk for 1 hour at RT. The membrane was probed with the primary antibody overnight at 4 °C after 3 washes with TBST buffer. The membrane was probed with HRP-conjugated secondary antibodies (Invitrogen, Goat anti-rabbit IgG - catalog#31460 and Goat anti-Mouse IgG - catalog#31430) at RT for 1-2 hours. The blots were developed using Clarity ECL Western Blotting Substrate (BioRad, catalog #170-5061) in ChemiDoc™ Imaging System (#12003153) from Bio-Rad.

### Chromatin Immunoprecipitation (ChIP)

Chromatin was isolated from 2x T-75 ultra-low-attachment flasks of MGG8 GSCs per condition, according to the manufacturer’s protocol (CST; Cat. no. 9003). Briefly, cells were cross-linked by adding 1% formaldehyde and incubating at RT for 10 minutes. Glycine 125 mM was used to quench the reaction by incubating the reaction for 5 minutes at RT. Nuclei were subsequently prepared using specific buffers, digested with MNase, and sonicated to generate chromatin fragments ranging from 150 to 900 bp. Sheared chromatin was incubated with 2 μg of the required antibody/antibodies for 16 hours, followed by incubation with Protein G magnetic beads (Invitrogen) for 4 hours at 4°C. An equal amount of IgG antibody was used as a negative control. Chromatin DNA was eluted by phenol-chloroform extraction, and the eluted DNA was used for quantitative PCR with a promoter-specific primer to amplify the target promoter region. The qPCR conditions were as follows: 95°C for 5 min; 95°C for 30 sec, 60°C for 30 sec, and 72°C for 30 sec for 40 cycles. Fold enrichment over IgG was calculated.

### Immunocytochemistry (ICC)

Cells were fixed in 4% paraformaldehyde for 30 minutes at 37°C. They were permeabilized with permeabilizing solution (1X PBS + 0.25% Triton X-100) for 20 minutes at RT, followed by blocking with blocking solution (1X PBS + 1% BSA + 0.3% Triton X-100 + 5% goat serum) for 1-2 hours at a gentle rocking speed. Cells were then incubated with the primary antibody against DDK and Na-K-ATPase, prepared in blocking solution, overnight at 4°C. After washing thrice with PBST (5 min each at RT with gentle rocking speed), cells were incubated with secondary antibody Alexa 488 anti-rabbit IgG (Invitrogen, catalog #A-11034) for 2-3 hours at RT at gentle rocking speed in the dark. DAPI (100 ng/ml) was added and incubated for 5 min at RT. Cells were then mounted on glass slides using a Fluoromount-G mounting medium (Invitrogen, catalog #00-4958-02) and covered with coverslips. Images were acquired in a Zeiss 880 microscope at 63X and quantified using Zen software.

### Orthotopic xenograft mouse model

The Institutional Animal Ethics Committee approved the following animal procedures (CAF/Ethics/076/2024). MGG8-GSCs were transduced with shNT and shSEZ6 lentivirus. Infected cells were subjected to puromycin selection for 72 hours, after which cells were harvested as single cells, and an equal number were used for intracranial injections. 0.1 × 10^6^ cells were stereotaxically injected into the hippocampus of 5-6 weeks-old athymic nude mice (coordinates: AP = +2.0, ML = +1.5, DV = 2.5) using a stereotaxic apparatus (Kopf and RWD). The survival of the shNT (n = 10) and shSEZ6 (n = 10) groups was monitored and plotted using GraphPad Prism 8. The Mantel-Cox log-rank test was utilized to measure the difference in survival between the two groups. (AP = Anterior-Posterior, ML = Medial-Lateral, DV = Dorsal-Ventral.)

### Cryo-sectioning of fixed mouse brain

Transcardiac perfusion was performed on selected mice with 1X PBS, followed by 4% paraformaldehyde. The mice brains were harvested and stored in 4% PFA for 24 hrs and subsequently in 30% sucrose solution for 48 hours. Poly-freeze solution (Sigma, #35059990) was used to embed the mouse brains, and the embedded brains were then sectioned at 30 μm thickness with an RWD Cryostat. The sections were stored at -80°C in Tissue Cutting Solution (0.1 M phosphate buffer, Ethylene glycol, and Glycerol).

### Immunohistochemistry (IHC)

For immunofluorescence, the brain sections were removed from TCS, put in 12-well plates, and washed thoroughly with 1X PBST. Following that, tissues were incubated in a permeabilization solution (1X PBS + 0.25% Triton-X 100) for one hour at RT. Next, they are blocked using blocking solution (1X PBS + 1% BSA + 0.3% Triton-X 100 + 10% goat serum) for 2-3 hours, followed by two PBST washes. Primary antibodies (CD133, SEZ6, CD31) prepared in blocking solution were added to the tissues in a 96-well plate and incubated overnight at 4°C. After 12-16 hours of incubation, the tissues were washed with PBST twice, and secondary antibodies (Invitrogen, Alexa Fluor 488 anti-mouse IgG, catalog #A11029; Alexa Fluor 594 anti-rabbit IgG, catalog #A11037), prepared in blocking solution, were added and left at RT for 2 hours. Subsequently, tissues were stained with DAPI (100ng/ml) for 30 minutes, followed by two PBST washes. Finally, the tissues were individually mounted on glass slides using the ProLong™ Glass Antifade Mountant (Invitrogen, #P36980) and covered by coverslips. Images were acquired in a Zeiss 880 microscope at 20X and quantified using Zen software.

### Hematoxylin and Eosin staining

Cryosections of mouse brain tissues were transferred onto poly-L-lysine-coated glass slides and incubated at 42°C for 48 hours. The slides were initially dipped in 70% ethanol for 1 minute, then in hematoxylin for 4 minutes. Subsequently, they were washed in distilled water for 1 minute and stained with eosin for 1 minute. After washing with distilled water, tissues were dipped in ethanol for differentiation of the staining, followed by another wash with water. The tissue sections were then mounted using a xylene-based DPX mounting medium and covered with coverslips. Images were taken using the Lawrence and Mayo digital microscope (0.7X).

### Scoring methods for confocal images

The areas of the blood vessels and the fluorescence intensities for all the fluorophores used in the tissue sections were measured using the Zeiss Black software (https://www.zeiss.com/microscopy/int/products/microscope-software.html), converted to percentages, and plotted as bar diagrams.

### Signal P analysis to identify potentially secreted proteins

A list of differentially expressed genes from the RNA-Seq data comparing GSC and DGC was used as input to SignalP 5.0, which predicts the presence and location of signal peptide cleavage sites in amino acid sequences from different organisms, including Gram-positive and Gram-negative prokaryotes and eukaryotes. The method incorporates predictions of cleavage sites and signal-peptide/non-signal-peptide status, based on a combination of several artificial neural networks. (http://www.cbs.dtu.dk/services/SignalP/).

### Gene Set Enrichment Analysis (GSEA)

The differentially expressed genes between the GSC and DGC (as identified in GSE54792) were pre-ranked by fold change and used as input for GSEA. All gene sets available in the Molecular Signature Database (MSigDB) were used to perform GSEA. We acknowledge our use of the GSEA software and MSigDB (**Subramanian et al., 2005**) (http://www.broad.mit.edu/gsea/).

### Survival analysis

Kaplan-Meier survival analysis was done using GraphPad Prism 5.0 (GraphPad Software, San Diego, California, USA).

### Quantification and Statistical Analysis

Bar diagrams were generated using Microsoft Excel. The box plots were generated using GraphPad Prism 5.0 (GraphPad Software, San Diego, California, USA). p-value is calculated by an unpaired t-test with Welch’s correction, or a student t-test was done using Microsoft Excel. Student t-test has been used to calculate p-value for all the bar graphs, and the scatter plots, where * is p<0.05, ** is p<0.01, *** is p<0.001, and ns is non-significant. The Mantel-Cox log-rank test was utilized to measure the difference in survival between the two groups

### Expression, isolation, and purification of the ectodomain of TGFβ RII

The expression, isolation, and purification of the TGFβ RII ectodomain (46-155aa) were performed as previously described (**Hinck et al., 2000).** In brief, the pET32a-TGFβRII ectodomain plasmid is transformed into the E. coli strain BL21(DE3)-CodonPlus selected on LB-agar plates containing chloramphenicol and ampicillin and scaled up in LB medium until reaching mid-log phase (0.6 OD_600_), where protein expression was induced with 0.8 mM IPTG for 3 hours at 37 °C in the form of insoluble inclusion bodies. The inclusion bodies were disrupted in a buffer containing 1 M NaCl, then treated with 1% Triton X-100 to remove contaminants and solubilized overnight in a buffer containing 8 M urea and 20 mM Tris (pH 7.0). This non-soluble fraction was clarified by centrifugation, and the supernatant was subjected to initial purification via DEAE-Sepharose anion-exchange chromatography, eluting with a linear 0–300 mM NaCl gradient. Target fractions identified by reducing SDS-PAGE were pooled, reduced with 25 mM DTT, and dialyzed against 100 mM acetic acid. Controlled protein refolding was initiated by slowly diluting the denatured sample into a folding buffer containing 200 mM Tris, 2 mM reduced-glutathione, and 0.5 mM oxidized-glutathione (pH 8.0), followed by incubation for 18–24 hours at 4 °C. Finally, the refolded solution was concentrated, dialyzed into 20 mM MES (pH 6.0), and polished using a Source 15Q HR 10/10 column with a linear 0–300 mM NaCl gradient. The purified native receptor fractions were dialyzed to remove salt, concentrated to 5–10 mg/mL, and stored at −20 °C.

### Isothermal Colorimetric assay (ITC)

Experiments were conducted on a MicroCal PEAQ ITC (Malvern Panalytical) machine. All experiments were conducted at 10 °C while stirring at 750 rpm in water. The cell (280 μl) was loaded with the purified TGFβ RII ectodomain, respectively, and the microsyringe was loaded with 40 μl of peptides. Either 50 µM or 40 µM TGFβ RII ectodomain is used for 30 µM peptides. Titration for i15 peptide was conducted using an initial injection of 0.4μl followed by 13 identical injections of 3 μl with a duration of 6 sec (per injection) and a spacing of 150 sec between injections. Titrations for i13 and i4 peptides were conducted using an initial injection of 0.4μl followed by 18 identical injections of 2 μl with a duration of 4 sec (per injection) and a spacing of 150 sec between injections. Thermodynamic parameters were calculated (Δ*G* = Δ*H* - *T*Δ*S* = -R*T*lnKB, where Δ*G*, Δ*H, and ΔS are the changes in free energy, enthalpy, and entropy of binding,* respectively) using the MicroCal PEAQ ITC Analysis software.

### Circular dichroism

Measurements of far-ultraviolet circular dichroism were performed using a JASCO-815 equipped with a temperature-controlled multi-cell holder. Wavelength scans were measured from 260 to 190 nm at 20 to 95 °C at a rate of 5 °C/min, and again at 25 °C after fast refolding (about 5 min). The experiments were performed using 50 μM peptides in water in a 1-mm path-length cuvette.

### Protocol for the synthesis of peptides having an amidated carboxy-terminal

Peptides were synthesized on Rink Amide AM resin (0.8 mmol g-1) on a 200 mg scale (0.16 mmol) using a standard Fmoc-based strategy (**Chatterjee et al., 2012**). The resin was swollen in DMF and deprotected with 20% piperidine in DMF (5 min x 1, 15 min x 1) followed by thorough washing with DMF (3 times). The C-terminal amino acid (3.5 eq.) was loaded onto the resin by using standard coupling reagents (2.5 equiv HOBt, 2.5 equiv DIC) in DMF for 2 hours at room temperature. The entire peptide was assembled using the same protocol.

### Cleavage from the resin and global deprotection

Peptides were cleaved off from the resin using TFA: TIPS: H2O (95: 2.5: 2.5) for 2 hours. The deprotected peptide was precipitated in chilled ethyl ether, yielding the crude product, and was further purified by RP-HPLC (solvent A: 0.1% TFA in water; solvent B: 0.1% TFA in acetonitrile; gradient 30-75% B over 30 minutes).

### Synthesis of Compound S2C

#### General Experimental Information

All reactions were carried out under a Nitrogen atmosphere. Boronate ester was purchased from BLD Pharm and used as received. Potassium phosphate (tribasic, anhydrous) and Cesium carbonate (anhydrous) were purchased from HYMA Synthesis and were stored in a desiccator. Pd_2_(dba)_3_ and ligands such as Brettphos and Xphos, and Morpholine amine were purchased from BLD Pharm, and [PdCl_2_(PPh_3_)_2_] was purchased from Sigma Aldrich. 1,4-Dioxane (AR Dry) used was purchased from Finar by Actylis. The solvent/solvent-water mixture was used only after thorough degassing. It was purged with nitrogen for 30 mins. A nitrogen environment was achieved in reaction vessels using balloons and 18-gauge needles. All other chemicals, reagents were purchased as reagent grade from standard commercial sources (BLD Pharm, TCI, Sigma Aldrich) and were used as received.

The compounds synthesized were characterized by 1H NMR, and HRMS. 1H Nuclear Magnetic Resonance spectra were recorded on an Ascend NMR-500MHz (Bruker) spectrometer. For the ^1^H and ^13^C NMR spectra, all chemical shifts are reported in parts per million (δ) units and are relative to the residual CDCl_3_ signal at 7.26 ppm and residual DMSO-*d_6_* at 2.5 ppm for ^1^H, respectively. HRMS analysis was performed using a Synapt G2S LC-MS (Waters).

### Synthesis of Compound S2C

Compound S2C was synthesized according to the synthetic route outlined in **Scheme 1**.

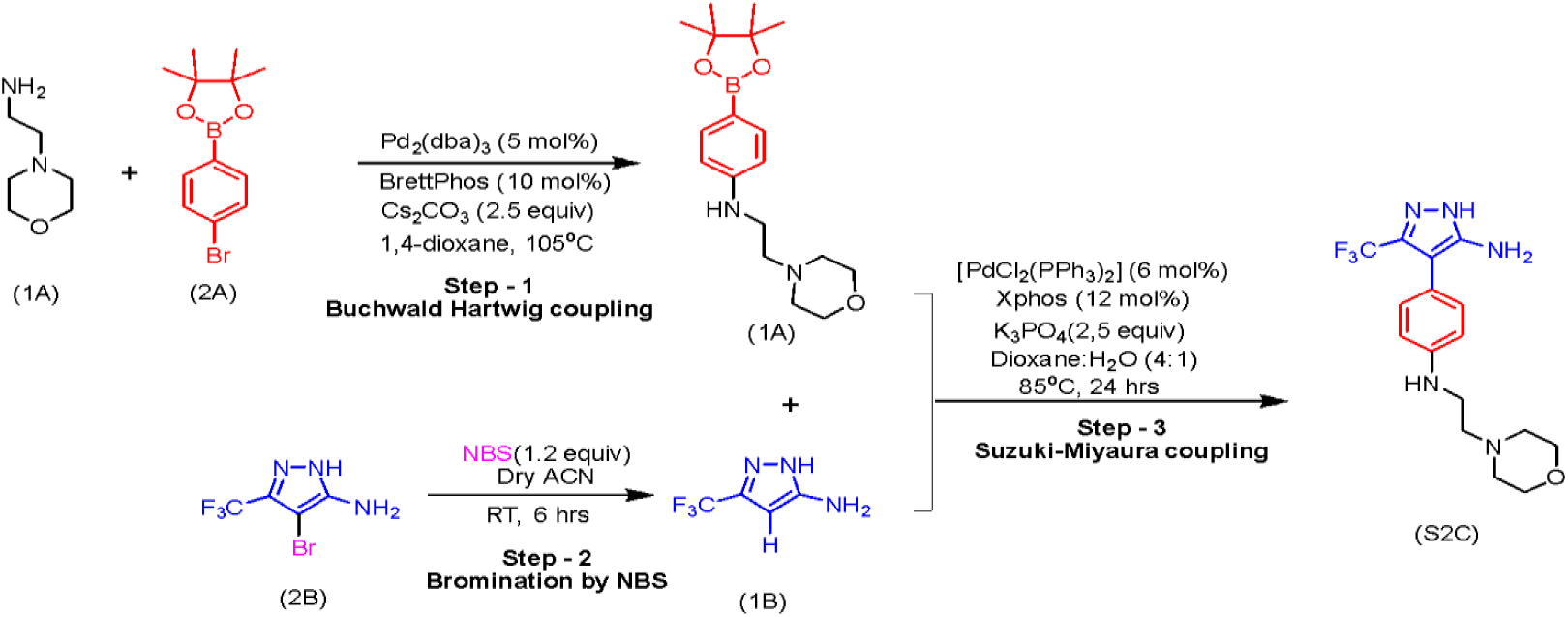

#### Synthesis of Compound 3A

The Boronate ester(2A) (1.06mmol) was reacted with Morpholino-ethyl amine (1A) (1.25 equiv, 1.32 mmol) in the presence of Pd_2_(dba)_3_ (5 mol%, 0.05mmol), BrettPhos (10 mol%, 0.10mmol) using 1,4-Dioxane (0.3M) as solvent under 105°C for 16 hours in a Buchwald-Hartwig style CN coupling **(Surry et al., 2008)**. After completion of the reaction, as monitored by TLC, the reaction mixture was cooled to room temperature, filtered through Celite, and then subjected to liquid-liquid extraction. The product was extracted with ethyl acetate (3 × 30mL), and the combined organic layers were washed with brine, dried over anhydrous sodium sulfate, filtered, and concentrated under reduced pressure. The crude product was purified by silica gel column chromatography using an ethyl acetate-hexane solvent system to afford compound 3A as a yellow-coloured sticky solid.

#### Compound 3A

Yield: 65-73%; R_f_ = 0.35 (ethyl acetate: hexane = 95:5).

**^1^H NMR** (500 MHz, DMSO-*d_6_*) *δ* (ppm): 7.38 (d, *J* =16.2 Hz, 2H), 6.54 (d, *J* =8.9 Hz, 2H), 5.84 (s, 2H), 3.58 (t, *J* =8.9 Hz, 4H), 3.15 (d, *J* =12.4 Hz, 2H), 2.56 (d, *J* =8.5 Hz, 4H), 2.42 (d, *J* =10.4 Hz, 2H), 1.24 (s, 12H);

#### HRMS (ESI)

m/z calculated for C_18_H_30_BN_2_O_3_^+^ [M+H]^+^: 333.2344; found: 333.2343.

The structure of compound 3A was confirmed by NMR spectroscopy and high-resolution mass spectrometry.

### Synthesis of Compound 2B

The Pyrazole linker (1.98 mmol) was brominated with N-bromosuccinimide (NBS) (1.2 equiv, 2.38 mmol) in acetonitrile at room temperature for 6 hours **(Zhao et al., 2007)**. After completion of the reaction, as monitored by TLC, the reaction mixture was then subjected to liquid-liquid extraction. The product was extracted with ethyl acetate (3 × 50mL), and the combined organic layers were washed with brine, dried over anhydrous sodium sulfate, filtered, and concentrated under reduced pressure. The crude product was purified by silica gel column chromatography using an ethyl acetate-hexane solvent system to afford compound 2B as an orange solid.

#### Compound 2B

Yield: 85-90%; R_f_ = 0.4 (ethyl acetate: hexane = 60:40).

**^1^H NMR** (500 MHz, DMSO-*d_6_*) *δ* (ppm): 12.56 (s, 1H), 5.62 (s, 2H);

#### HRMS (ESI)

m/z calculated for C_4_H_4_BrF_3_N_3_^+^ [M+H]^+^: 229.3535; found: 229.3526.

The structure of compound 2B was confirmed by high-resolution mass spectrometry and NMR spectroscopy.

### Synthesis of Compound S2C

The Intermediates **3A** (0.30 mmol) and **2B** (1.1 equiv, 0.33 mmol) were coupled using Suzuki-Miyaura style CC cross coupling in the presence of [PdCl_2_(PPh_3_)_2_] (6 mol%, 0.018 mmol) as Pd catalyst and Xphos (12 mol%, 0.036 mmol) as ligand. K_3_PO_4_ (2.5 equiv, 0.75 mmol) was used as base with 1,4-Dioxane (0.2M) and water mixture as a solvent under 85°C for 24 hours **(Miyaura et al., 1995).** After completion of the reaction, as monitored by TLC, the reaction mixture was cooled to room temperature, filtered through Celite, and then subjected to liquid-liquid extraction. The product was extracted with ethyl acetate (3 × 30mL), and the combined organic layers were washed with brine, dried over anhydrous sodium sulfate, filtered, and concentrated under reduced pressure. The crude product was purified by column chromatography on silica gel using a DCM-methanol solvent system to afford the final compound S2C as a Dark Brown, sticky solid.

#### Compound S2C

Yield: 30-37%; R_f_ = 0.27 (DCM: Methanol = 95:5).

**^1^H NMR** (500 MHz, DMSO-*d_6_) δ* (ppm): 12.20 (s, 1H), 6.99 (d, *J* =16.6 Hz, 2H), 6.60 (d, *J* =16.9 Hz, 2H), 4.93 (s, 2H), 3.60 (s, 4H), 3.15 (s, 2H), 2.53 (d, *J* =3.5 Hz, 2H), 2.46 (s, 4H); (500 MHz, CDCl_3_) *δ* (ppm): 7.40 (d, *J* =16 Hz, 2H), 6.65 (d, *J* =17 Hz, 2H), 3.66 (d, *J* =9 Hz, 2H), 3.12 (m, 2H), 2.59 (d, *J* =11 Hz, 2H), 2.43 (s, 4H);

#### HRMS (ESI)

m/z calculated for C_16_H_21_F_3_N_5_O ^+^ [M+H]^+^: 356.1693; found: 356.1712.

The structure of compound S2C was confirmed by high-resolution mass spectrometry and NMR spectroscopy (Spectra in Supplementary files).

### *In silico* studies of S2C with BACE1 Protein and Ligand Preparation

The crystal structure of human β-secretase (BACE1) in complex with a bound inhibitor (PDB ID: 3ZMG; resolution 1.74 Å) was retrieved from the RCSB Protein Data Bank **(Hilpert et al., 2013)**. The protein structure was processed using the Protein Preparation Wizard in Schrödinger Maestro (version 2026-1) **(Sastry et al., 2013)**. Preprocessing steps included assigning bond orders, adding hydrogen atoms, generating disulfide bonds, and removing crystallographic waters more than 5.0 Å from heteroatoms. Protonation states of ionizable residues were assigned at pH 7.0 ± 2.0 using PROPKA **(Olsson et al., 2011)**. The structure was subsequently energy-minimized using the OPLS4 force field **(Lu et al., 2021)**, with a heavy-atom RMSD convergence threshold of 0.30 Å. The 2D structure of compound S2C was sketched in 2D Sketcher and prepared using LigPrep, which generated possible ionization states at pH 7.0 ± 2.0 **(Epik; Shelley et al., 2007)** and low-energy 3D conformers with the OPLS4 force field **(Lu et al., 2021)**.

### Molecular Docking

A receptor grid was generated around the centroid of the co-crystallized ligand in 3ZMG using the Receptor Grid Generation panel in Glide, with an enclosing box of 10 × 10 × 10 Å and outer box scaled to ligand length. Molecular docking of S2C into the BACE1 active site was performed using Glide SP docking **(Friesner et al., 2006)** in the Maestro module of Schrödinger Suite, version 2026-1. Default Glide scoring function parameters were used, and the best-scoring pose was selected based on Glide Score and visual inspection of the binding mode for chemically reasonable interactions with the catalytic dyad and flap region.

### Molecular Dynamics Simulation

The best-scoring BACE1–S2C docked complex was used as the starting structure for all-atom molecular dynamics (MD) simulations performed using Desmond **(Bowers et al., 2006)** Schrödinger Suite, version 2026-1. The complex was solvated in an orthorhombic simulation box with a buffer distance of 10 Å using the TIP3P water model **(Jorgensen et al., 1983).** The system was neutralized by adding [Na⁺/Cl⁻] counter-ions to achieve a physiological ionic concentration of 0.15 M NaCl. The OPLS4 force field **(Lu et al., 2021)** was used to describe all system parameters. The solvated system was relaxed using the default Desmond relaxation protocol prior to production simulation. Production MD was carried out for 1000 ns (1 μs) under the NPT ensemble at 300 K and 1.01325 bar, with the Nose-Hoover thermostat and Martyna-Tobias-Klein barostat, respectively, using a time step of 2.0 fs. Long-range electrostatic interactions were treated using the Particle Mesh Ewald (PME) method **(Essmann et al., 1995)** with a cutoff radius of 9.0 Å.

### Trajectory Analysis

Trajectory stability and conformational dynamics were assessed by computing root-mean-square deviation (RMSD) of the protein backbone and ligand heavy atoms (relative to the first frame, post-alignment), root-mean-square fluctuation (RMSF) per residue, protein–ligand contact persistence (H-bonds, hydrophobic contacts, ionic/water-bridge interactions), and radius of gyration (Rg), using the Simulation Interaction Diagram (SID) tool in Maestro.

### ADME/Pharmacokinetic Property Prediction

Absorption, distribution, metabolism, and excretion (ADME) properties of compound S2C were predicted using QikProp **(Jorgensen and Duffy, 2002)**, which estimates pharmacokinetically relevant molecular descriptors and evaluates compliance with Lipinski’s Rule of Five **(Lipinski et al., 2001)**. The energy-minimized, LigPrep-processed structure of S2C was used as input. QikProp was run in normal mode, generating physicochemical properties, including partition coefficients (QPlogPo/w, QPlogPoct, QPlogPw), blood-brain barrier permeability (QPlogBB), serum albumin binding (QPlogKhsa), hydrogen-bond donor and acceptor counts, and molecular weight. All predicted values were compared against recommended optimal ranges for orally active drug-like molecules.

### AlphaFold2 modelling

Amino acid sequences of Sushi-3, Sushi-4, and Sushi-5 domains from human SEZ6 were obtained from UniProt (Accession: Q53EL9). Three-dimensional structures of individual domains and the contiguous Sushi-3-4-5 stretch were predicted using locally installed ColabFold (v1.5.2) **(Mirdita et al., 2022)** with GPU acceleration. Multiple sequence alignments were generated using MMseqs2 against UniRef30 and environmental databases, with an MSA depth of 512:1024. The PDB100 database was utilized for template-guided modeling. Predictions were run with 12 recycles, with early stopping enabled when the pLDDT deviation fell to 0 across consecutive recycles. To improve conformational sampling, stochastic dropout was activated. Each prediction used 20 random seeds, generating five models per seed, yielding a total of 100 models per protein. All models underwent AMBER force-field relaxation with 2000 minimization steps to optimize stereochemistry. Structural quality of AlphaFold2 predictions was evaluated using pLDDT profiles and PAE matrices, along with independent stereochemical validation via a Ramachandran plot-based assessment of backbone dihedral angles using PROCHECK **(Laskowski et al., 1993)**. All figures were generated using custom R scripts.

### Protein-protein docking

The crystal structure of TGFβ RII ectodomain (ECD) was obtained from the Protein Data Bank (PDB ID: 1KTZ). Structure pre-processing, including removal of extraneous chains and residue renumbering, was performed using pdb-tools **(Rodrigues et al., 2018)**. Blind protein-protein docking of the TGFβ RII ectodomain against Sushi-3-4-5 was performed using five independent docking platforms: ClusPro2.0 **(Kozakov et al., 2018)**, GRAMM **(Singh et al., 2024)**, HDOCK **(Yan et al., 2020)**, InterEvDock3 **(Quignot et al., 2021)**, and LZerD **(Christoffer et al., 2021)**. Docking parameters were standardized to generate 100 models for ClusPro2.0, GRAMM, and HDOCK, and 50 models for InterEvDock3 and LZerD, with complexes subsequently ranked according to algorithm-specific scoring metrics. Interfacial residues within 8 Å were identified using custom Python scripts, and top-ranked complexes were structurally superimposed and rendered using UCSF ChimeraX (v1.15) **(Meng et al., 2023)**.

### Sequence and structure alignment

The amino acid sequences of the Sushi-3, Sushi-4, and Sushi-5 domains of SEZ6 were retrieved from UniProt (Accession: Q53EL9) and aligned using Clustal Omega **(Sievers et al., 2011)** using BLOSUM62 scoring matrix, gap opening penalty of 10.0, gap extension penalty of 0.05, and mBed-like clustering for guide tree generation. Output alignments were generated in Clustal format and visualized using ESPript (v3.2) **(Gouet et al., 1999)**. AlphaFold2-predicted structures of Sushi-3, Sushi-4, and Sushi-5 domains were structurally aligned using UCSF ChimeraX (v1.15). Pairwise sequence-guided structural alignments were performed using the Needleman-Wunsch algorithm with BLOSUM-62 amino acid scoring matrix, gap opening penalty of 12.0, gap extension penalty of 4.0, and iterative refinement with 2 Å distance cutoff for C_α_ atoms. Root mean square deviation (RMSD) values were calculated for each pairwise superposition, and superimposed structures were visualized and rendered using UCSF ChimeraX.

### All-atom molecular dynamics simulations and analysis

Simulation systems were built using the CHARMM-GUI input generator **(Lee et al., 2016)**. Solvation was performed with TIP3P water molecules in a cubic box with roughly 10 Å of water padding. Counter-ions (Na⁺ and Cl⁻) were added to achieve physiological ionic strength (0.15 M NaCl) and neutralize the net system charge. Simulations were carried out using GROMACS (v2023.3) **(Van Der Spoel et al., 2005)** on a GPU-accelerated high-performance computing server, with the CHARMM36m force-field **(Huang et al., 2017)**. Energy minimization was carried out using the steepest-descent algorithm for up to 50,000 steps or until the maximum force converged below 100 kJmol^−1^nm^−1^. Bond lengths involving hydrogen atoms were constrained using the LINCS algorithm, and the Verlet cutoff scheme was used for non-bonded interactions. Following minimization, a two-step equilibration protocol was applied. NVT ensemble equilibration was performed for 1 ns at 303.15 K using the V-rescale thermostat. NPT ensemble equilibration was performed for 1 ns at 303.15 K and 1 bar, maintained with the Parrinello-Rahman barostat. After equilibration, all positional restraints were removed, and a 1000 ns production simulation was performed in the NPT ensemble at 303.15 K and 1 bar using a 2 fs integration time step. Van der Waals interactions were modeled using the Lennard-Jones potential, and Coulombic interactions were calculated using the Particle Mesh Ewald (PME) method. Periodic boundary conditions were maintained in all three dimensions. Trajectory coordinates and velocities were saved every 100 ps, producing 10,000 frames for subsequent analysis. Binding free energy (ΔG) calculations were performed using gmx_MMPBSA (v1.6.4) **(Valdes-Tresanco et al., 2021)**. Intermolecular pairwise distances were computed with MDAnalysis (v2.8.0) **(Michaud-Agrawal et al., 2011),** and non-covalent interactions were characterized through PyContact (v1.0.2) **(Scheurer et al., 2018)**. Residue-wise secondary-structure assessment of peptides was conducted using MDTraj (v1.10.0) (**McGibbon et al., 2015**) with the DSSP algorithm. Coiled-coil heptad repeat patterns in peptides were identified using Socket2 **(Kumar et al., 2021)**. All figures were generated through custom R scripts.

### *De novo* peptide binder design

*De novo* peptide binder design was performed using BindCraft **(Pacesa et al., 2025)** on a GPU-accelerated high-performance computing server. Cysteine residues were excluded to prevent uncontrolled disulfide formation. Multimer design mode was enabled to capture realistic binding interfaces through simultaneous co-folding of peptide and receptor. A four-stage iterative refinement protocol was employed for design optimization, comprising 75 soft iterations, 45 temporary iterations, 5 hard iterations, and 15 greedy iterations at a 5% perturbation rate, to progressively narrow sequence diversity. Loss function weights were optimized to prioritize interface complementarity: inter-chain contact weight (0.5), intra-chain contact weight (0.5), interface pTM weight (0.05), global pLDDT weight (0.1), inter-chain pAE weight (0.2), and intra-chain pAE weight (0.4). Interfacial contacts were defined using an inter-residue distance cutoff of 20.0 Å and an intra-chain distance cutoff of 14.0 Å, with a minimum of two contacts per residue to ensure substantial interface formation. Helicity weighting (0.95) with fixed helicity was used to promote stable helical scaffold structures. AlphaFold2 structure predictions were optimized to employ 1 recycling iteration during design for computational efficiency and 3 recycling iterations during validation for enhanced confidence. ProteinMPNN sequence sampling was performed with soluble weights and low sampling temperature (0.1) to generate 10 sequence variants per design trajectory while preserving scaffold geometry. Interface residues were fixed during ProteinMPNN refinement to maintain the designed binding geometry. The top 2 sequences per trajectory passing initial quality filters were retained. High-confidence designs were selected using comprehensive multi-metric criteria involving average pLDDT ≥ 0.8, average i_pLDDT ≥ 0.8, average pTM ≥ 0.55, average i_pTM ≥ 0.5, average pAE ≤ 0.35 Å, average i_pAE ≤ 0.35 Å, average surface hydrophobicity ≤ 0.35, average shape complementarity ≥ 0.6, average PackStat ≥ 0.65, ≥7 average interface residues, ≥3 average interface H-bonds, ≤ 4 average unsatisfied H-bonds, average binder RMSD ≤ 3.5 Å, average binder energy score ≤ 0, average dSASA ≥ 1, average dG ≤ 0, average hotspot RMSD ≤ 6 Å, and ≤ 3 lysines/methionines at interface. Maximum 500 design trajectories were generated, with unrelaxed trajectories and complexes automatically removed.

## Results

### An integrated secretome analysis identifies Wnt-β-catenin signaling-regulated SEZ6 as a CSC-specific proangiogenic molecule in glioma

The cancer stem cell (CSC) secretome plays a major role in processes such as extracellular matrix (ECM) remodeling, Tumor microenvironment (TME) recruitment and activation, metastatic spread, and resistance to radio- and chemotherapy (**Lopez de Andres et al., 2020**). A close association between CSCs and tumor vasculature has been reported in many cancers (**Lv et al., 2023; Lizarraga-Verdugo et al., 2020; Thirant et al., 2012; Calabrese et al., 2007; Wang et al., 2010; Ricci-Vitiani et al., 2010; Soda et al., 2010**). To dissect the proangiogenic functions of GSCs, we investigated the secreted proteome (**Sengupta et al., 2022**) and transcriptome (**Suva et al., 2014**) of three patient-derived GSCs (MGG4, MGG6, and MGG8) compared with their matched differentiated glioma cells (DGCs). GO analysis of differentially abundant proteins in the GSC conditioned medium (CM) identified significant enrichment of many terms related to “angiogenesis” (**Figure 1A**). Additionally, GO analysis of secreted differentially expressed genes (DEGs) in GSCs revealed enrichment for “angiogenesis”-related terms (**Supplementary Figure 1A**), establishing a connection between the CSC secretome and angiogenesis. To confirm this observation, we tested the ability of the GSC CM to induce angiogenic network formation by immortalized human pulmonary microvascular endothelial cells (ST1). GSC CM collected from three GSCs induced efficient angiogenesis in ST1 cells (**Figure 1B**).

**Figure 1:**
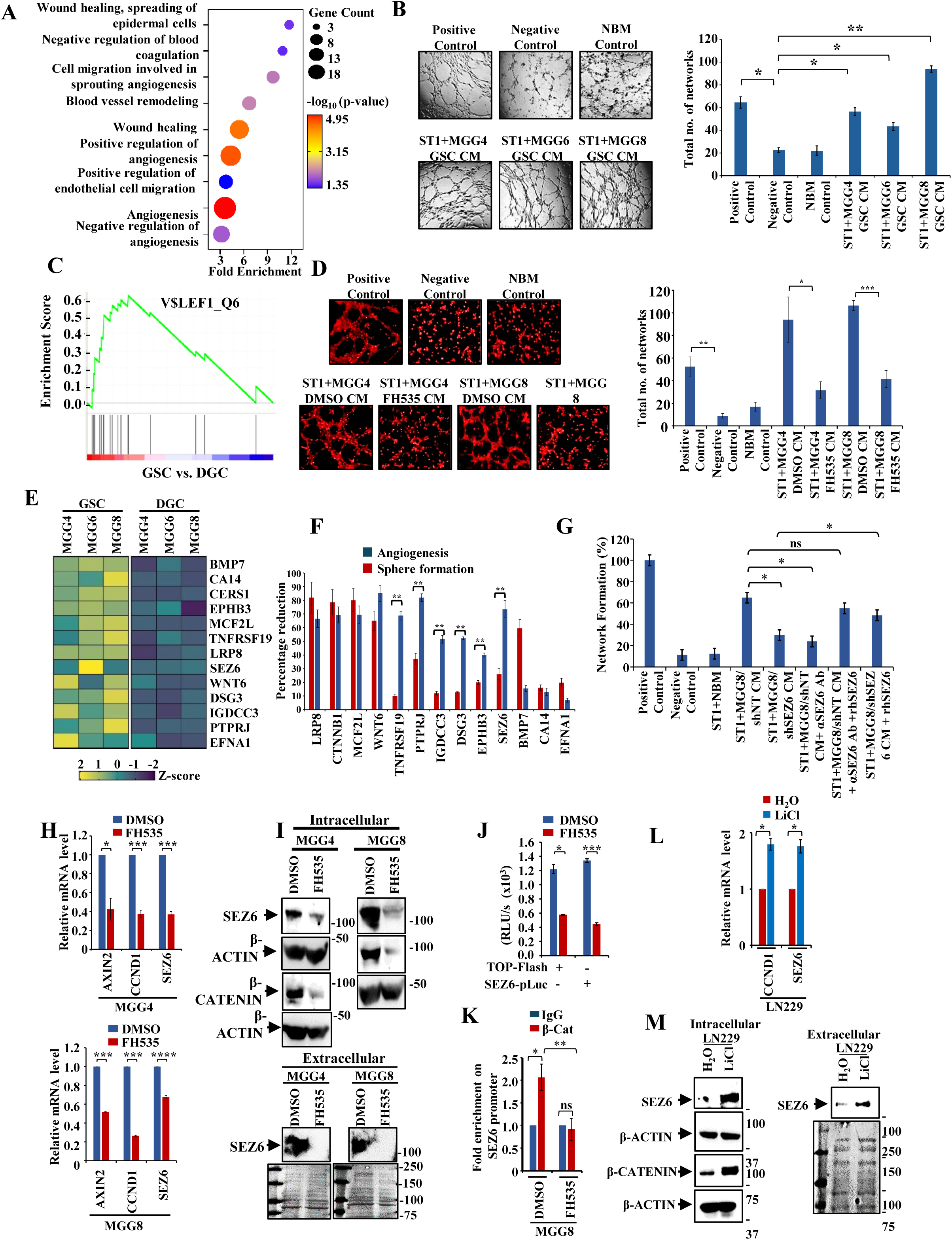
Identification of SEZ6 as a GSC-specific, Wnt/β-Catenin regulated, pro-angiogenic protein. (A) Bubble plot showing significant enrichment of several angiogenesis-related terms in the secretome of Glioma Stem-like Cells (GSCs) over Differentiated Glioma Cells (DGCs) when the list of differentially abundant proteins in the secretome of GSCs versus DGCs was subjected to Gene Ontology (GO) analysis. Gene Count indicates the number of genes from the input list involved in the corresponding GO term. p-Value <0.05 is considered significant. (B) Representative images of the in vitro network formation by ST1 cells upon treatment with CM collected from the three patient-derived GSCs (MGG4, MGG6, and MGG8). In the upper panel, the positive control is ST1 cells plated in serum-supplemented complete endothelial cell medium. The negative control is for ST1 cells plated in incomplete medium. The NBM control is for ST1 cells plated in incomplete neurobasal media (NBM). In the bottom panel, ST1 cells are plated with the CM collected from the indicated GSC lines. Magnification 4X, Scale 200 µm. The bar graph shows the total number of complete networks in each condition. p-Value <0.05 was considered significant with *, **, and *** representing p-values <0.05, 0.01, and 0.001, respectively. ns, non-significant. (C) GSEA plot depicts the enrichment of the gene set V$LEF_Q6 in GSCs over DGCs, where DEGs from the RNA sequencing data were subjected to GSEA analysis. (D) Representative images of the in vitro network formation by ST1 cells upon treatment with CM collected from DMSO-treated GSCs and FH535-treated GSCs, across two GSC lines, MGG4 and MGG8. In the upper panel, the positive control is ST1 cells plated in serum-supplemented complete endothelial cell medium. The negative control is for ST1 cells plated in incomplete medium. The NBM control is for ST1 cells plated in incomplete neurobasal media (NBM). In the bottom panel, ST1 cells are plated with CM collected from the indicated GSC lines and their corresponding treatments. Magnification 4X, Scale 200 µm. The bar graph shows the total number of complete networks in each condition. p-Value <0.05 was considered significant with *, **, and *** representing p-values <0.05, 0.01, and 0.001, respectively. ns, non-significant. (E) Heatmap showing the Z-scores of the thirteen genes of the TCF/LEF gene set (“V$LEF_Q6”), obtained from the RNA sequencing data GSC vs DGC. A higher Z-score (yellow) indicates upregulation of LEF target genes in GSCs compared to DGCs (blue). (F) Composite graph showing the impact, in terms of percentage reduction, on neurosphere formation (red bars) upon silencing each of the thirteen genes of the LEF target gene set in GSCs, followed by the collection of CM from the silenced GSCs for determining the percentage reduction in the in-vitro network formation assay of ST1 cells. p-Value <0.05 was considered significant with *, **, and *** representing p-values <0.05, 0.01, and 0.001, respectively. ns, non-significant. (G) Bar graph showing the percentage network formation by ST1 cells upon treatment with CM collected from MGG8-shNT and MGG8-shSEZ6, followed by rescue with rhSEZ6 in shSEZ6 CM and inhibition by SEZ6-specific antibody in shNT CM. The positive control is for ST1 cells plated with serum-supplemented complete endothelial cell media, the negative control is for ST1 cells plated in incomplete medium, and the NBM control is for ST1 cells plated in incomplete neurobasal media (NBM). p-Value <0.05 was considered significant with *, **, and *** representing p-values <0.05, 0.01, and 0.001, respectively. ns, non-significant. (H) RT-qPCR showing downregulation of the SEZ6 transcript in MGG4 GSC (upper panel) and MGG8 GSC (lower panel) upon treatment with FH535 (10 µM for 24 hours) compared to DMSO. Downregulation of bona fide targets of the Wnt/β-catenin pathway, AXIN2 and CCND1, is used as a positive control. p-Value <0.05 was considered significant, with *, **, and *** representing p-values <0.05, 0.01, and 0.001, respectively. ns, non-significant. (I) Western blot showing reduced expression of intracellular (upper panel) and extracellular (lower panel) SEZ6 protein upon treatment of MGG4 GSC and MGG8 GSC with FH535 (10 µM for 24 hours) compared to DMSO. Reduced unphosphorylated β-Catenin is shown as a marker of inhibition of the Wnt β-Catenin pathway (upper panel). The Ponceau-stained membrane shows equal loading of the conditioned medium (lower panel). (J) SEZ6-promoter-luciferase (SEZ6-pLuc) activity showing reduced luciferase activity of the SEZ6 promoter upon treatment of MGG8 GSC with FH535 (10 µM for 24 hours) compared to DMSO. The TOP-Flash construct is used as a positive control because it is a known reporter luciferase plasmid for the canonical Wnt/β-Catenin pathway. p-Value <0.05 was considered significant, with *, **, and *** representing p-values <0.05, 0.01, and 0.001, respectively. ns, non-significant. (K) RT-qPCR showed significantly higher fold enrichment of β-Catenin on the SEZ6 promoter, which was reduced upon treatment of MGG8 GSCs with FH535 (10 µM for 24 hours) compared to DMSO. p-Value <0.05 was considered significant, with *, **, and *** representing p-values <0.05, 0.01, and 0.001, respectively. ns, non-significant. (L) RT-qPCR showing significant upregulation of the SEZ6 transcript in LN229 cells upon treatment with LiCl (10 mM for 24 hours), as compared to water. The upregulation of bona fide target of the Wnt β-catenin pathway, CCND1, is used as a positive control. p-Value <0.05 was considered significant, with *, **, and *** representing p-values <0.05, 0.01, and 0.001, respectively. ns, non-significant. (M) Western blot showing higher expression of intracellular (left panel) and extracellular (right panel) SEZ6 protein upon treatment of LN229 cells with LiCl (10 mM for 24 hours) compared to water. Increased unphosphorylated β-Catenin is shown as a marker of activation of the Wnt β-Catenin pathway (left panel). The Ponceau-stained membrane shows equal loading of the conditioned medium (right panel).

Next, we sought to delineate the signaling pathways governing the angiogenic potential of GSCs. Investigation of GSC-specific secreted DEGs by Gene Set Enrichment Analysis (GSEA) showed a sole significant enrichment of “V$LEF1_Q6” gene set (**Figure 1C**), which consists of genes that contain the binding motif of the LEF1 transcription factor (https://www.gsea-msigdb.org/gsea/msigdb/cards/LEF1_Q6). The heterodimeric transcription factors β-catenin/LEF1 (lymphoid enhancer-binding factor; also called T-cell factor, TCF) are important mediators of the Wnt/β-catenin signaling pathway (**Liu et al., 2022**). As the Wnt-β-catenin signaling pathway has been shown to promote angiogenesis (**Daneman et al., 2008; Olson et al., 2017; Easwaran et al., 2003; Hu et al., 2009**), the secreted targets of β-catenin may participate in GSC secretome-induced angiogenesis. Conditioned medium collected from GSCs treated with FH535 (a small-molecule inhibitor of Wnt/β-catenin signaling) showed a significant loss of ability to induce angiogenesis (**Figure 1D**), confirming the critical role of Wnt/β-catenin signaling. To identify the key angiogenesis-inducing molecule, we focused on the 13 enriched genes in the “V$LEF1_Q6” gene set, which showed higher expression in GSCs than in DGCs (**Figure 1E**). GSC-specific higher expression of these genes was independently validated (**Supplementary Figure 1B**). We then silenced the thirteen enriched genes (**Supplementary Figure 1C**) individually and tested their requirement for GSC growth (as measured by neurosphere growth and a limiting dilution assay) and for the ability of GSC CM to induce angiogenesis. This analysis revealed that silencing TNFRSF19, PTPRJ, IGDCC3, DS3, EPHB3, and SEZ6 affected the ability of GSC CM to induce angiogenesis significantly, with minimal effect on GSC neurosphere growth (**Figure 1F; Supplementary Figure 1D, E, and F**). Since the role of TNFRSF19, PTPRJ, IGDCC3, DS3, and EPHB3 in tumorigenesis is well established (**Guo et al., 2020**; **Smart et al., 2012; Yao et al., 2023; Wang et al., 2025; Jang et al., 2020),** we focused on Seizure Related 6 Homolog (SEZ6), which has limited literature. SEZ6 is a membrane protein shed into the stroma upon cleavage by ***b***eta-site ***a***myloid precursor protein ***c***leaving ***e***nzyme ***1*** (BACE1) (**Pigoni et al., 2016**). While SEZ6 has been implicated in dendritic arborization and in motor and cognitive functions (**Gunnersen et al., 2007; Nash et al., 2020**), there are no reports on its role in tumorigenesis and angiogenesis.

Next, we evaluated the angiogenic potential of SEZ6. The capacity of conditioned media (CM) from MGG8/shNT cells to induce angiogenesis was significantly attenuated upon preincubation with an anti-SEZ6 antibody; this effect was effectively reversed by the addition of recombinant SEZ6 (rhSEZ6) (**Figure 1G; Supplementary Figure 1G**). Consistently, CM derived from SEZ6-silenced GSCs (MGG8/shSEZ6) exhibited a reduced ability to promote angiogenesis, which was restored following supplementation with rhSEZ6 (**Figure 1G; Supplementary Figure 1G**). Furthermore, exogenous rhSEZ6 enhanced angiogenic network formation in both human brain microvascular endothelial cells (HBMECs) and mouse brain-derived endothelial cells (B.End3) (**Supplementary Figures 2A and 2B**). In addition, rhSEZ6 markedly stimulated migration, invasion, and proliferation of ST1 endothelial cells (**Supplementary Figures 2C, D, and E**). Collectively, these findings demonstrate that SEZ6, present in the GSC secretome, is a potent inducer of angiogenesis.

**Figure 2:**
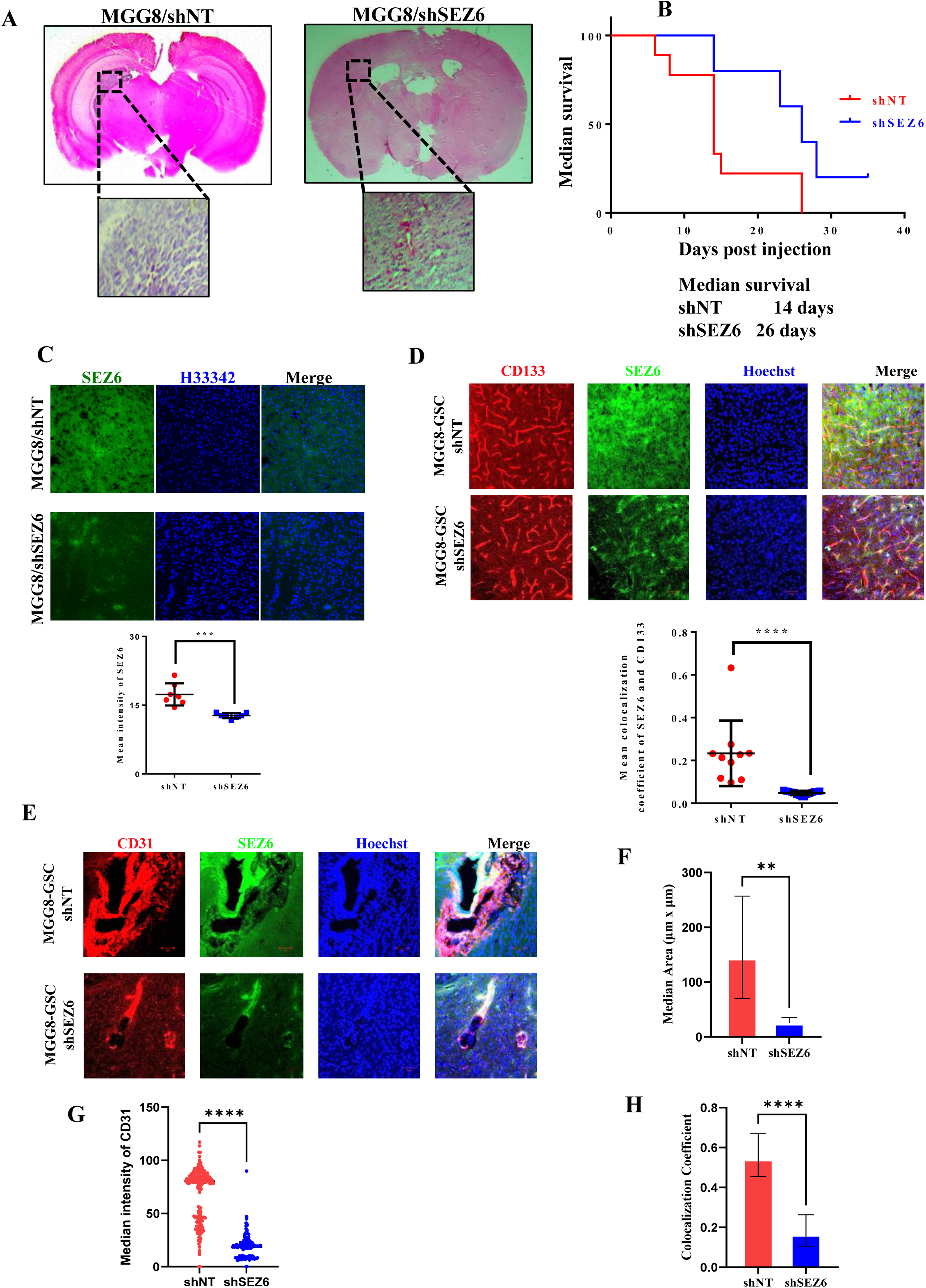
Impact of SEZ6 on glioma growth in vivo. (A) Representative H&E micrographs of the whole brain sections showing a visibly larger tumor in animals injected with MGG8/shNT GSCs compared to animals injected with MGG8/shSEZ6 GSCs, on day 30 of tumor growth. (B) Kaplan Meier survival plot showing a significant improvement in the median survival in the MGG8/shSEZ6 group (26 days) compared to the MGG8/shNT group (14 days). The p-value of 0.464 was calculated by the Log-rank (Mantel-Cox) test. (C) Representative IHC images of brain sections showing significant downregulation of SEZ6 in the MGG8/shSEZ6 group, as shown by a decrease in SEZ6 mean intensity compared to the MGG8/shNT group. p-Value was calculated by Student T-test, and <0.05 was considered significant with *, **, and *** representing p-values <0.05, 0.01, and 0.001, respectively. ns, non-significant. (D) Representative IHC images of brain sections showing a significant decrease in mean colocalization coefficient between CD133 (red) and SEZ6 (green) in the MGG8/shSEZ6 group compared to the MGG8/shNT. p-Value was calculated by Student T-test, and <0.05 was considered significant with *, **, and *** representing p-values <0.05, 0.01, and 0.001, respectively. ns, non-significant. (E) Representative IHC images of brain sections showing visibly smaller blood vessels in the MGG8/shSEZ6 group, as shown by CD31 staining (red) compared to the MGG8/shNT group. The significant reduction in the mean blood vessel area in the MGG8/shSEZ6 group compared to the MGG8/shNT group is shown in panel (F). p-value was calculated by Student T-test, and <0.05 was considered significant with *, **, and *** representing p-values <0.05, 0.01, and 0.001, respectively. ns, non-significant. (G) Plot showing a significant reduction in the mean intensity of CD31 in the MGG8/shSEZ6 group compared to the MGG8/shNT group. p-value was calculated by Student T-test, and <0.05 was considered significant with *, **, and *** representing p-values <0.05, 0.01, and 0.001, respectively. ns, non-significant. (H) Plot showing a significant decrease in mean colocalization coefficient between CD31 and SEZ6 in the MGG8/shSEZ6 group compared to the MGG8/shNT. p-value was calculated by Student T-test, and <0.05 was considered significant with *, **, and *** representing p-values <0.05, 0.01, and 0.001, respectively. ns, non-significant.

We next examined the regulation of SEZ6 by the Wnt/β-catenin signaling pathway. Pharmacological inhibition of this pathway using FH535 led to a marked reduction in both SEZ6 transcript (**Figure 1H**) and protein levels (**Figure 1I**) in GSCs. Consistently, FH535 treatment significantly diminished luciferase activity driven by both the β-catenin–responsive TOP Flash reporter and the SEZ6 promoter construct in MGG8 GSCs (**Figure 1J**). Chromatin immunoprecipitation (ChIP) analysis further confirmed the direct occupancy of β-catenin at the SEZ6 promoter in MGG8 GSCs, an interaction that was abolished upon FH535 treatment (**Figure 1K**). Notably, SEZ6 expression at both the transcript and protein levels was elevated in GSCs compared to DGCs (**Supplementary Figures 1H, I, and J**). Conversely, activation of the Wnt/β-catenin pathway in DGCs using LiCl resulted in a significant upregulation of SEZ6 expression (**Figure 1L and M**). Collectively, these findings establish that SEZ6 is a downstream target of Wnt/β-catenin signaling and that GSC-secreted SEZ6 functions as a key mediator of angiogenesis.

### Sez6 is required for in vivo glioma growth but not for in vitro GSC growth

The above findings indicate that SEZ6 is dispensable for GSC proliferation but plays a significant role in promoting angiogenesis in vitro. To further delineate the contribution of SEZ6 to glioma progression *in vivo*, we assessed tumor growth in NIH nu/nu mice orthotopically implanted with MGG8 GSC cells. Notably, MGG8/shSEZ6 GSCs formed significantly smaller tumors compared to MGG8/shNT controls (**Figure 2A**), accompanied by a marked increase in overall survival (**Figure 2B**). Consistent with effective knockdown, tumors derived from MGG8/shSEZ6 cells exhibited reduced SEZ6 expression (**Figure 2C**). Immunofluorescence analysis revealed substantial colocalization of SEZ6 with the stem cell marker CD133 in MGG8/shNT tumors, which was markedly diminished in MGG8/shSEZ6 tumors (**Figure 2D**). Furthermore, MGG8/shSEZ6 tumors exhibited reduced vascularization, characterized by smaller vessel caliber, decreased CD31 staining, and diminished colocalization of SEZ6 with CD31 compared with control tumors (**Figure 2E, F, G, and H**). Based on these data, we demonstrate that GSC-expressed SEZ6 promotes tumor angiogenesis and is critical for glioma growth *in vivo*.

### SEZ6-induced angiogenesis requires the TGFβ pathway to induce IL8 in endothelial cells

To elucidate the SEZ6-induced signaling pathway in endothelial cells that promotes angiogenesis, we subjected intracellular protein extracts from ST1 cells treated with CM derived from MGG8/shNT and MGG8/shSEZ6 GSCs to the Proteome Profiler Human Angiogenesis Array (R&D Systems), which quantifies the relative levels of 55 angiogenesis-related proteins. This analysis identified seven proteins that are expressed at high levels in endothelial cells treated with MGG8/shNT GSC-derived CM but downregulated significantly upon treatment with the MGG8/shSEZ6 GSC-derived CM (**Figure 3A; Supplementary Figure 3A**). We chose IL-8, the most downregulated gene, as the potential mediator of SEZ6-induced angiogenesis. IL-8 is a well-studied pro-angiogenic cytokine that forms part of an angiogenic signature commonly found in endothelial cells (**Fahey and Doyle, 2019**). First, we confirmed that IL-8 is induced by SEZ6, as evidenced by reductions in IL-8 transcript and protein levels (intracellular and extracellular) in ST1 cells treated with CM from MGG8/shSEZ6 compared to MGG8/shNT (**Figure 3B, C, and D**). The addition of recombinant human IL-8 (rhIL-8) induced angiogenic network formation in ST1 cells, like rhSEZ6 (**Figure 3E; Supplementary Figure 3B**). However, the addition of rhSEZ6 failed to induce angiogenic networks in IL-8-silenced ST1 cells (**Figure 3E; Supplementary Figure 3B**). Further, the addition of rhIL-8 induced IL-8-silenced ST1 cells to form angiogenic networks (**Figure 3E; Supplementary Figure 3B**). These experiments confirm that IL-8 mediates SEZ6-induced angiogenesis in ST1 cells.

**Figure 3.**
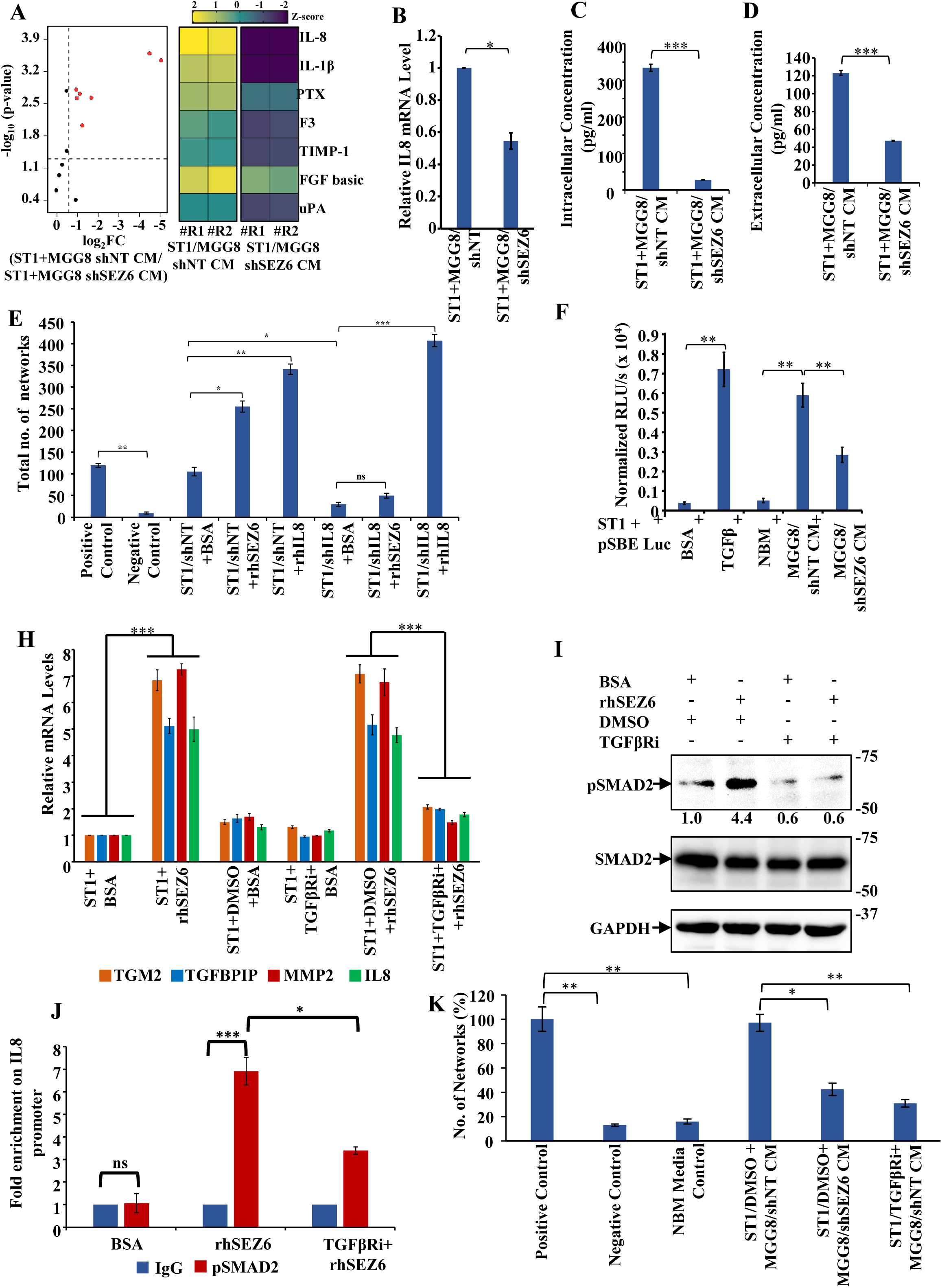
SEZ6 induces IL8 in a TGF β pathway-dependent manner in endothelial cells to promote angiogenesis. (A) Scatter plot showing the downregulated proteins in ST1 cells upon treatment of ST1 cells with CM derived from MGG8-shNT versus MGG8-shSEZ6. Mean pixel densities were measured for both conditions (n=22), and proteins with a mean pixel density exceeding 20,000 in the MGG8-shNT condition were selected for further analysis (n=13). Mean pixel density values were log2-transformed, and log2 fold change was calculated. Proteins that showed a log2-fold downregulation of 0.58 or greater and a log10 p-value of -1.3 or less are indicated as red spots (n=7). Black spots represent the proteins that did not show significant downregulation. The heatmap (right panel) shows the Z-scores of the significantly downregulated proteins (n=7) in duplicate samples for each condition (ST1/MGG8 shNT and ST1/MGG8 shSEZ6). (B) RT-qPCR showing significant downregulation of IL8 transcript in ST1 cells upon treatment with CM derived from MGG8-shSEZ6 as compared to when treated with CM derived from MGG8-shNT. p-value <0.05 was considered significant, with *, **, and *** representing p-values <0.05, 0.01, and 0.001, respectively, and ns, non-significant. (C) Bar graph showing a significant decrease in the intracellular level of IL8 protein in ST1 cells upon treatment with CM derived from MGG8-shSEZ6 as compared to when treated with CM derived from MGG8-shNT, as measured by ELISA. p-value <0.05 was considered significant; *, **, and *** denote p-values <0.05, 0.01, and 0.001, respectively; ns, non-significant. (D) Bar graph showing a significant decrease in the extracellular level of IL8 protein in ST1 cells upon treatment with CM derived from MGG8-shSEZ6 as compared to when treated with CM derived from MGG8-shNT, as measured by ELISA. p-value <0.05 was considered significant; *, **, and *** denote p-values <0.05, 0.01, and 0.001, respectively; ns, non-significant. (E) Bar graph showing a significant reduction in total networks formed by ST1/shIL8 cells as compared to ST1/shNT cells, followed by treatment of ST1/shNT cells with rhSEZ6 and rhIL8, which show a marked increase in network formation, and treatment of ST1/shIL8 cells with rhSEZ6, which had no impact, while treatment with rhIL8 increased the network formation. p-Value <0.05 was considered significant, with *, **, and *** representing p-values <0.05, 0.01, and 0.001, respectively, and ns, non-significant. (F) Bar graph showing a significant induction in luciferase activity in SBE-Luc, a TGFβ pathway-dependent reporter luciferase construct, in ST1 cells treated with CM derived from MGG8-shNT, which is compromised when cells are treated with CM derived from MGG8-shSEZ6. The activation of SBE-Luc upon treatment with TGFβ is a positive control for the reporter luciferase construct, as it is a canonical ligand. Treatment with NBM serves as a negative control for CMs derived from GSCs. p-value <0.05 was considered significant; *, **, and *** denote p-values <0.05, 0.01, and 0.001, respectively; ns, non-significant. (G) RT-qPCR showing transcript level of IL8, and the bona fide targets of TGFâ pathway, TGM2, TGFBIP, and MMP2, upon pretreatment of ST1 cells with DMSO or TGFβ-RI inhibitor (TGFβRi), SB341542 (10 µM) for 12 hours, followed by addition of BSA or rhSEZ6. p-value <0.05 was considered significant; *, **, and *** denote p-values <0.05, 0.01, and 0.001, respectively; ns, non-significant. (H) Western blot showing pSMAD2 and total SMAD2 protein level in ST1 cells upon pretreatment of ST1 cells with DMSO or TGFβ-RI inhibitor (TGFβRi), SB341542 (10 µM) for 12 hours, followed by addition of BSA or rhSEZ6. The numbers indicate densitometric quantification of pSMAD2 bands using Image Lab (BIORAD) and normalized to total SMAD2. (I) RT-qPCR showing significantly higher fold enrichment of pSMAD2 at the IL8 promoter upon treatment of ST1 cells with rhSEZ6, which was compromised upon pretreatment of ST1 cells with TGFβ-RI inhibitor (TGFβRi), SB341542 (10 µM), followed by addition of rhSEZ6. p-Value <0.05 was considered significant, with *, **, and *** representing p-values <0.05, 0.01, and 0.001, respectively. ns, non-significant. (J) Bar graph showing the per cent network formation by ST1 cells upon pretreatment with TGFâ-RI inhibitor (TGFâRi), SB341542 (10 µM), followed by addition of CM derived from MGG8/shNT. The positive control is ST1 cells plated in serum-supplemented complete endothelial cell medium. The negative control is for ST1 cells plated in incomplete medium. The NBM control is for ST1 cells plated in incomplete Neurobasal Media (NBM). p-value <0.05 was considered significant, with *, **, and *** representing p-values <0.05, 0.01, and 0.001, respectively, and ns, non-significant.

Next, we investigated the signaling pathway activated by SEZ6 that induces IL-8 production in endothelial cells. SEZ6 encodes a 994 amino acid protein which contains two CUB (complement subcomponent ***C***1r, C1s/ sea urchin embryonic growth factor ***U***egf/ bone morphogenetic protein1 ***B***mp1) domains and five Sushi (CCP/SCR) domains (**Shimizu-Nishikawa et al., 1995; Pigoni et al., 2016**). Reports demonstrated that different proteins containing CUB domains activate EGFR signaling (**Law et al., 2016; Dong et al., 2011**), Src (**Liu et al., 2011**), Hedgehog signaling (**Jakobs et al., 2014; Wierbowski et al., 2020**), and the TGFβ signaling pathway (**Predes et al., 2019**). Our investigation identified that SEZ6 activates the TGFβ pathway, as seen by the fact that the CM from MGG8/shNT GSCs, but not from MGG8/shSEZ6 GSCs, induced luciferase activity from pSBE-Luc (contains SMAD binding elements) (**Figure 3F**). As expected, TGF-β1 addition activated luciferase activity from pSBE-Luc in ST1 cells (**Figure 3F**). Further, high levels of TGM2, TGFBIP, MMP2 (bona fide TGFβ targets), and IL8 transcripts seen in ST1 cells treated with CM derived from MGG8/shNT GSCs are decreased upon treatment with the CM of MGG8/shSEZ6 GSCs (**Figure 3G**). The addition of rhSEZ6 to ST1 cells efficiently induced TGM2, TGFBIP, MMP2, and IL8 transcripts, but this effect was not observed after treatment with a TGFβR inhibitor (**Figure 3G**). Additionally, rhSEZ6 treatment induced pSMAD2 levels in ST1 cells, but not after treatment with a TGFβ inhibitor (**Figure 3H**), and showed a several-fold enrichment of pSMAD2 on the IL-8 promoter (**Figure 3I**). More importantly, the ability of the CM from MGG8/shNT GSCs to induce ST1 cells to form angiogenic networks is severely affected by the TGFβR inhibitor treatment (**Figure 3J; Supplementary Figure 3C**). Based on these results, we conclude that SEZ6 induces IL-8 by activating the TGF-β signaling pathway in ST1 cells, thereby promoting angiogenesis.

### SEZ6 interacts with TGFβ RII through its Sushi 3 domain to induce angiogenesis

Based on our findings demonstrating that SEZ6 activates TGFβ signaling in endothelial cells, we next explored the mechanistic basis of this effect by probing a potential physical and functional interaction between SEZ6 and the TGFβ type II receptor (TGFβ RII). The extracellular domain of the TGFβ RII receptor binds specifically to the TGFβ class of ligands (TGFβ1-3) (**Goebel et al., 2019**). SEZ6 and TGFβ RII expressing constructs (DDK-SEZ6 and HA-TGFβ RII, respectively) were transfected into HEK-293T cells, and the localization of both proteins was analyzed by confocal microscopy. While TGFβ RII staining is seen primarily on the cell membrane, the SEZ6 staining is seen on the membrane and cytoplasm, with TGFβ RII and SEZ6 having a co-localization index of 95% (**Figure 4A**), suggesting the possibility of an interaction between SEZ6 and TGFβ RII. To confirm this, we conducted co-immunoprecipitation experiments using protein extracts derived from HEK-293T cells transfected with DDK-SEZ6 and HA-TGFβ RII constructs. SEZ6 immunoprecipitation (α-DDK) efficiently pulled down TGFβ RII (**Figure 4B**). Conversely, TGFβ RII immunoprecipitation (α-HA) efficiently pulled down SEZ6 (**Figure 4C**). To identify the regions of SEZ6 that mediate the interaction with TGFβ RII, we created several deletions of SEZ6 (**Figure 4D).** SEZ6 with a deletion of either or both CUB domains (SEZ6 ΔC1, SEZ6 ΔC2, and SEZ6 ΔC1/2) bound to TGFβ RII efficiently (**Figure 4E**). Similarly, SEZ6 with a deletion of the DR (disordered region; SEZ6 ΔDR) domain bound to TGFβ RII efficiently (**Figure 4E**). However, SEZ6 with a deletion of Sushi domains 3, 4, and 5 (SEZ6 ΔS3/4/5 and SEZ6 ΔS3/4/5, ΔC1/2) failed to bind to TGFβ RII (**Figure 4E**), suggesting Sushi domains 3/4/5 mediate the interaction between SEZ6 and TGFβ RII proteins. To confirm this finding, we expressed the SEZ6 protein lacking all domains except the Suchi domains 3, 4, and 5 (SEZ6 ΔDR, ΔS1/2, ΔC1/2) and tested its ability to interact with TGFβ RII. The SEZ6 protein containing the Sushi 3/4/5 domains alone interacted efficiently with TGFβ RII (**Figure 4F and G**), thus confirming that the Sushi 3/4/5 domains of SEZ6 mediate the interaction with TGFβ RII.

**Figure 4.**
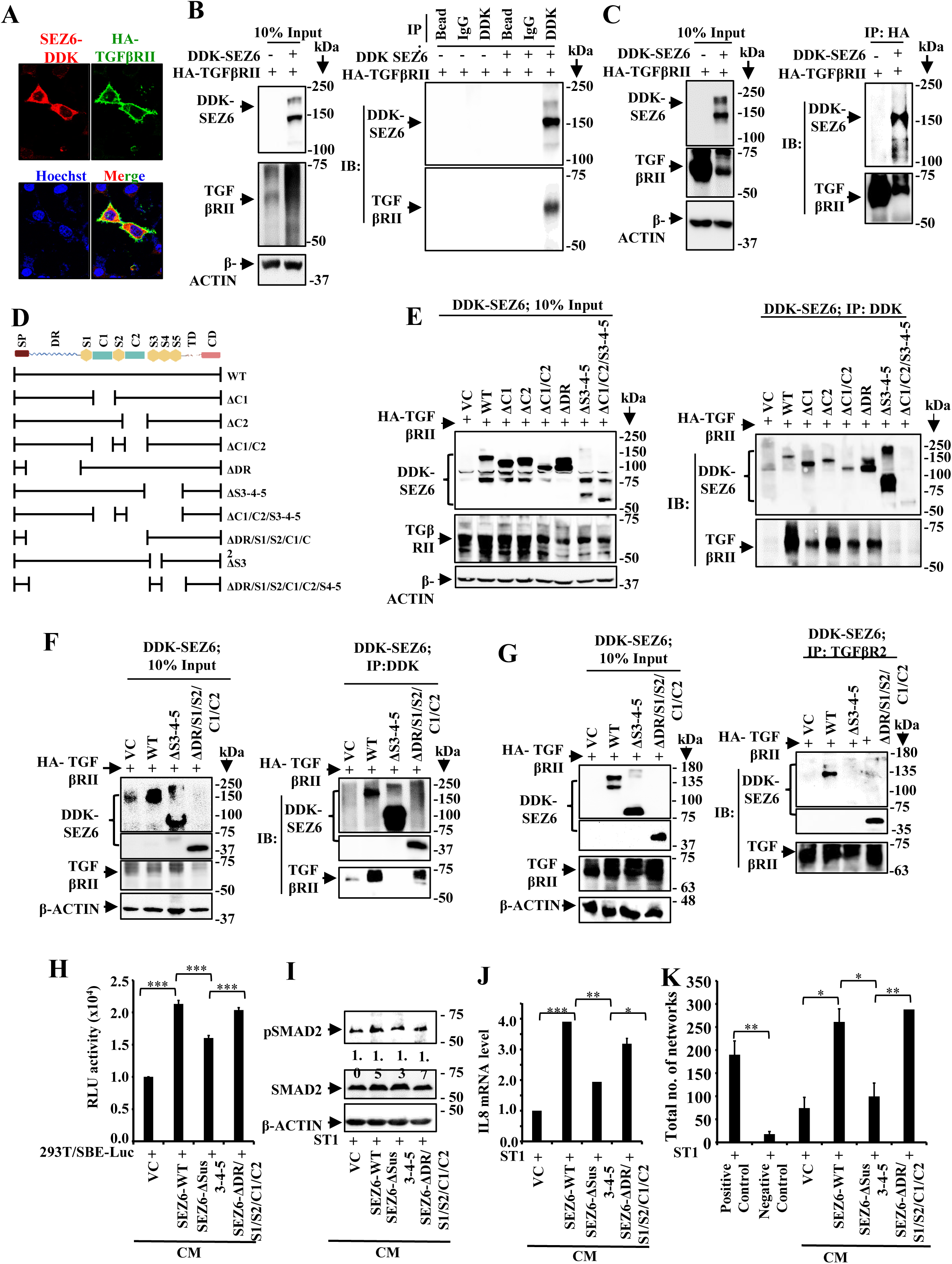
Sushi 3-4-5 domains in SEZ6 are essential for TGFβ pathway activation in endothelial cells to promote angiogenesis. (A) Representative images of confocal microscopy showing membrane colocalization of DDK-tagged SEZ6 (red) and HA-tagged TGFβ RII (green) ectopically expressed in HEK-293T cells. The merged image displays yellow pixels on the cell membrane, indicating the presence of both proteins. Hoechst dye is used to stain the nucleus. Magnification 63X. (B) Western blot showing direct interaction of SEZ6 and TGFβ RII in HEK-293T cells ectopically co-expressing SEZ6-DDK and HA-TGFβ RII, upon immunoprecipitation with DDK, followed by immunoblotting with TGFβ RII, compared to ectopic co-expression of vector control (VC) and TGFβ RII. The left panel shows the Western blot for 10% input, and the right panel shows the Western blot for DDK-immunoprecipitation. (C) Western blot showing direct interaction of SEZ6 and TGFβ RII in HEK-293T cells ectopically co-expressing SEZ6-DDK and HA-TGFβ RII, upon immunoprecipitation with HA, followed by immunoblotting with TGFβ RII, compared to ectopic co-expression of vector control (VC) and TGFβ RII. The left panel shows the Western blot of the 10% input, and the right panel shows the Western blot from the HA-immunoprecipitation. (D) Schematic representation of the domain structure of the SEZ6 protein, comprised of CUB domains and Sushi domains. The following line diagrams represent the deletion of the specific domain as indicated in the figure. (WT- Wild type, SP- Signal peptide; DR- Disordered region; S1- Sushi 1; S2-Sushi 2; C1- CUB1; S2-Sushi 2; C2- CUB2; S3- Sushi 3; S4- Sushi 4, S5-Sushi 5, TD- Transmembrane domain, CD- Cytoplasmic domain). (E) Western blot showing the interaction profile of WT-SEZ6 and the corresponding deletion constructs of SEZ6 with TGFβ RII in HEK-293T cells ectopically co-expressing the indicated DDK-tagged SEZ6 constructs and HA-TGFβ RII, upon immunoprecipitation with DDK, followed by immunoblotting with TGFβ RII. The left panel shows the Western blot for 10% input, and the right panel shows the Western blot for DDK-immunoprecipitation. (F) Western blot showing loss of interaction with TGFβ RII upon Sushi 3-4-5 deletion, followed by restoration of interaction upon expression of only Sushi 3-4-5 in HEK-293T cells ectopically co-expressing the corresponding DDK-tagged SEZ6 constructs and HA-TGFβ RII, upon immunoprecipitation with DDK, followed by immunoblotting with TGFβ RII. The left panel shows the Western blot for 10% input, and the right panel shows the Western blot for DDK- immunoprecipitation. (G) Western blot showing loss of interaction of TGFβ RII with SEZ6 upon Sushi 3-4-5 deletion, followed by restoration of interaction upon expression of only Sushi 3-4-5 in HEK-293T cells ectopically co-expressing the corresponding DDK-tagged SEZ6 constructs and HA-TGFβ RII, upon immunoprecipitation with HA, followed by immunoblotting with DDK. The left panel shows the Western blot for 10% input, and the right panel shows the Western blot for DDK- immunoprecipitation. (H) Bar graph showing SBE-luciferase (SBE-Luc) activity upon treatment of SBE-Luc- transfected HEK-293T cells with the CM containing the shed forms WT-SEZ6 and the shed forms of the indicated deletion mutants of SEZ6. VC CM is used as a negative control for the CMs containing the corresponding shed forms of SEZ6 proteins. p-value <0.05 was considered significant with *, **, and *** representing p-values <0.05, 0.01, and 0.001, respectively. ns, non-significant. (I) Western blot showing pSMAD2 and total SMAD2 protein level in ST1 cells upon treatment with the CM containing the shed forms WT-SEZ6 and the shed forms of the indicated deletion mutants of SEZ6 for 6 hours in incomplete medium. VC CM is used as a negative control for the CMs containing the corresponding shed forms of SEZ6 proteins. The numbers indicate densitometric quantification of pSMAD2 bands using Image Lab (BIORAD) and normalized to total SMAD2. (J) RT-qPCR showing levels of IL8 transcript in ST1 cells upon treatment with the CM containing the shed forms WT-SEZ6 and the shed forms of the indicated deletion mutants of SEZ6 for 14 hours in incomplete medium. VC CM is used as a negative control for the CMs containing the corresponding shed forms of SEZ6 proteins. p-value <0.05 was considered significant with *, **, and *** representing p-values <0.05, 0.01, and 0.001, respectively. ns, non-significant. (K) Bar graph showing the percentage network formation by ST1 cells upon treatment with CM containing the shed forms WT-SEZ6 and the shed forms of the indicated deletion mutants of SEZ6. VC CM is used as a negative control for the CMs containing the corresponding shed forms of SEZ6 proteins. The positive control is ST1 cells plated in serum-supplemented complete endothelial cell media, and the negative control is ST1 cells plated in incomplete medium. p-value <0.05 was considered significant with *, **, and *** representing p-values <0.05, 0.01, and 0.001, respectively. ns, non-significant.

Since Sushi3/4/5 domains of SEZ6 alone could interact with TGFβ RII, we further investigated whether this portion of SEZ6 could activate the TGFβ signaling pathway. The CM collected from HEK-293T cells transfected with full-length SEZ6 (SEZ-WT) or a deletion of Sushi 3, 4, and 5 domains (SEZ6-ΔS3/4/5) or containing only Suchi 3, 4, and 5 domains (SEZ6-ΔDR/S1/S2/C1/C2) and their ability to activate the TGFβ pathway in HEK-293T cells and induce angiogenesis of ST1 endothelial cells was tested. While the SEZ-WT and SEZ6-ΔDR/S1/S2/C1/C2, but not SEZ6-ΔS3/4/5 activated luciferase activity from pSBE-Luc, increased the pSMAD2 levels and induced angiogenic network formation (**Figure H, I, and K; Supplementary Figure 4A**), signifying that the Sushi 3, 4, and 5 domains of SEZ6 interact with TGFβ RII to activate the TGFβ pathway and angiogenesis.

Next, we sought to delineate which of the Sushi 3/4/5 domains mediates the interaction with TGFβ RII. We predicted the structure of the fragment carrying Sushi 3/4/5 domains (aa 708 to 897) by AlphaFold 2, and the top-ranked structure (**Supplementary Figure 5A, B, C, and D**) was allowed to dock blindly with the extracellular domain of TGFβ RII (TGFβ RII ECD) using five different docking tools (**Supplementary Figures 5E, F, G, H, and I)**. This exercise predicted a more favorable binding of the Sushi 3 domain, not 4 or 5 domains, to TGFβ RII ECD (**Figure 5A**). Sushi domains are typically 60 amino acids long, with conserved four cysteine residues that form the characteristic two disulfide bonds (**Ichinose et al., 1990**). To identify differences between the Sushi 3 and Sushi 4/5 domains, we analysed primary amino acid sequences for similarity and identity, and predicted the structures of the Sushi 3, 4, and 5 domains. The sequence similarity search identified four conserved cysteines across all three Sushi domains, with similarity indices of 40-47% and identity indices of 27-34% between Sushi domains 3, 4, and 5 (**Figure 5B**). Next, we predicted the structures of the Sushi-3, -4, and -5 domains individually using AlphaFold2 and confirmed their quality using pLDDT, PAE analysis, and Ramachandran plots (**Figure 5C and D; Supplementary Figures 6A, B, and C**). Upon superimposition, the structures of Sushi 3, -4, and -5 were found to be significantly similar (**Figure 5E**). Since the primary and secondary structures failed to reveal any unique differences between the Sushi 3 and Sushi 4/5 domains, we studied the interactions of the Sushi 3, 4, or 5 domains individually with the TGFβ RII ECD using molecular dynamics simulations (MDS; 1 microsecond). This analysis revealed that Sushi 3 binds stably to the TGFβ RII ECD, with the lowest ΔG (Gibbs Free Energy) of -33.01 kcal/mol, followed by Sushi-5 (-30.43 kcal/mol) and Sushi-4 (-0.36 kcal/mol) (**Figure 5F**). While Sushi-3 and Sushi-5 had similar average ΔG values in the molecular simulation study, the contact map analysis from the MDS revealed that more residues from Sushi-3 were within 20 Å distance from TGFβ RII-ECD than those from Sushi-5 (**Figure 5G**). Additionally, bond formation analysis revealed that Sushi 3, but not Sushi 4 or 5, formed several strong H bonds, Salt bridges, and Hydrophobic interactions throughout the simulation period (**Supplementary Figures 7, 8, and 9**). These results collectively demonstrate that the Sushi 3 domain mediates the interaction between SEZ6 and TGFβ RII.

**Figure 5:**
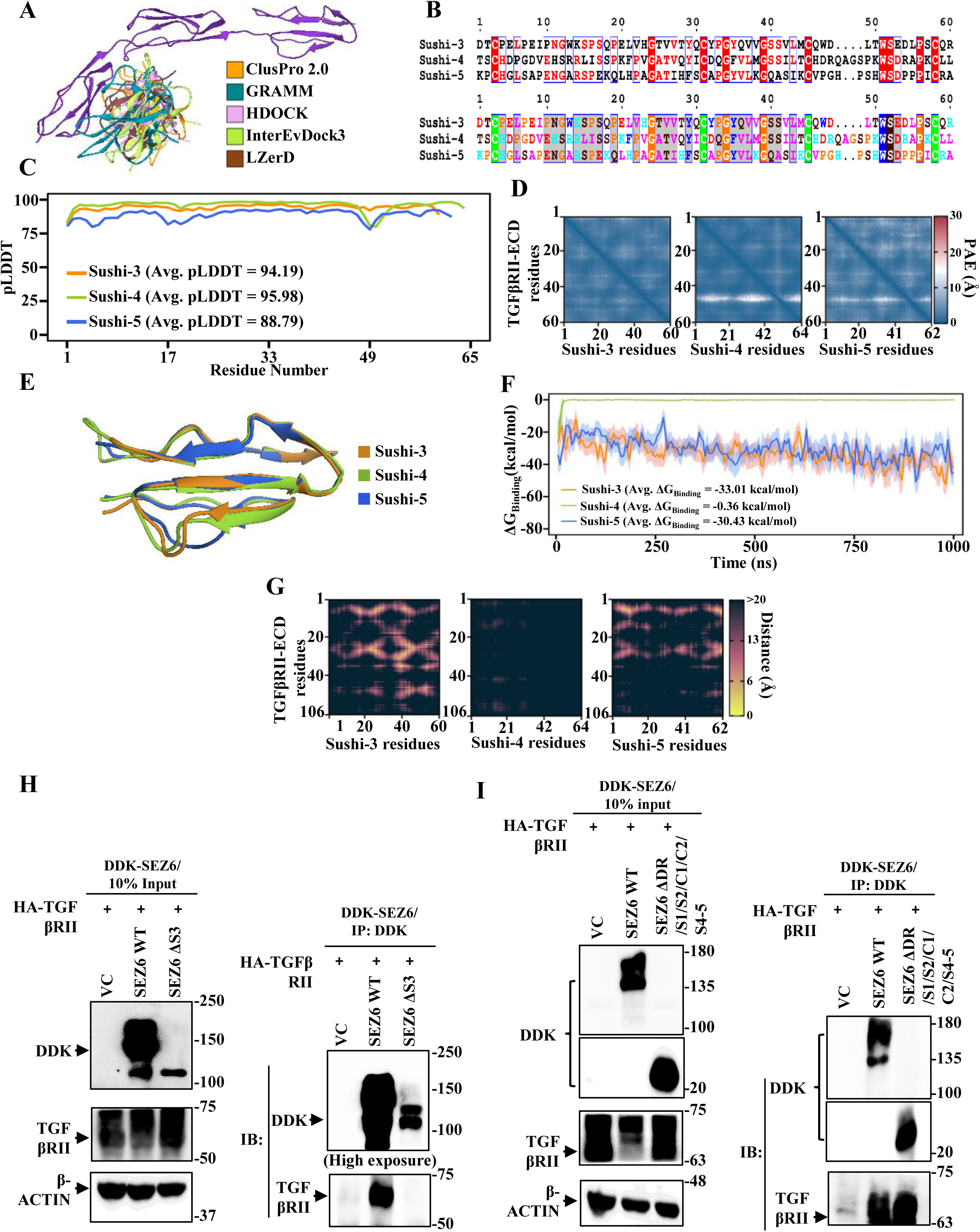
Computational and experimental determination of Sushi-3 as the most important and sufficient domain for interaction with TGFβ RII. **A)** Superposition of top-ranked docked complexes reveals consensus in binding interfaces across five independent methods, identifying Sushi-3 as the primary TGFβ RII-interacting domain. **B)** Multiple sequence alignment (MSA) of Sushi-3, Sushi-4, and Sushi-5 performed using Clustal Omega, with identity (top) and similarity (bottom) matrices highlighting conserved hallmark residues: cysteines (green strip), tryptophan (blue strip), and glycines/prolines (orange strip). **C)** pLDDT (Predicted Local Distance Difference Test) scores for AlphaFold2-predicted structures of Sushi-3, Sushi-4, and Sushi-5 show high residue-level structural reliability across all three domains. **D)** PAE (Predicted Aligned Error) matrices reveal low pairwise positional error (blue spots), indicating high confidence in relative residue positioning in AlphaFold2-predicted structures of Sushi-3, Sushi-4, and Sushi-5. **E)** Structural alignment shows high backbone similarity among the three Sushi domains, validating the conserved β-sandwich architecture. **F)** MM-PBSA (Molecular Mechanics Poisson-Boltzmann Surface Area) analysis quantifying TGFβ RII binding affinities reveals Sushi-3 as the strongest TGFβ RII binder, Sushi-5 with favorable binding, and Sushi-4 exhibiting negligible affinity. **G)** Average pairwise residue distance matrices computed using 1000 ns MD trajectories in MDAnalysis show Sushi-3 (left) forming extensive close-range contacts (yellow/pink spots) with TGFβ RII, compared to Sushi-4 (center) and Sushi-5 (right). **(H)** Western blot showing the co-expression of wild-type SEZ6 (SEZ6 WT) and Sushi-3 deleted SEZ6 (SEZ6 ΔS3), along with TGFβ RII in HEK-293T cells, in the input blot (left panel). The right panel shows the loss of interaction of SEZ6 ΔS3 with TGFβ RII upon immunoprecipitation with DDK antibody, while SEZ6 WT maintains the interaction with TGFβRII. **(I)** Western blot showing the co-expression of wild-type SEZ6 (SEZ6 WT) and only Sushi-3 expressing SEZ6 (SEZ6 ΔDR /S1/S2/C1/C2/S4-5), along with TGFβ RII in HEK-293T cells, in the input blot (left panel). The right panel shows that the only Sushi-3-expressing SEZ6 (SEZ6 ΔDR /S1/S2/C1/C2/S4-5) is sufficient for interaction with TGFβ RII upon immunoprecipitation with a DDK antibody, comparable to the interaction of wild-type SEZ6 with TGFβ RII.

To confirm the above results, we conducted co-immunoprecipitation experiments using protein extracts derived from HEK-293T cells transfected with full-length SEZ6 (SEZ-WT) or a deletion of the Sushi 3 domain (SEZ6-ΔS3) or containing only the Sushi 3 domain (SEZ6- ΔDR/S1/S2/C1/C2/S4-5). SEZ6 immunoprecipitation with α-DDK or a reverse immunoprecipitation with α-TGFβ RII demonstrated an efficient interaction between TGFβ RII and a full-length SEZ6 (SEZ6-WT) or SEZ6 containing the Sushi 3 domain alone (SEZ6- ΔDR/S1/S2/C1/C2/S4-5), but not SEZ6 with a deletion of the Sushi 3 domain (SEZ6-ΔS) (**Figure 5H and I**). These results confirm that the Sushi 3 domain is required for the interaction between SEZ6 and TGFβ RII.

### SEZ6-targeting therapeutic approaches to inhibit glioma

#### Generative AI develops a peptide binder to inhibit SEZ6/TGFβ RII interaction, inhibits angiogenesis, and glioma tumor growth

Designing and developing peptide binders to modulate protein-protein interactions that drive complex biological processes has great therapeutic and biotechnological potential. While several methods are available (**Cao et al., 2022; Gainza et al., 2023; Bennett et al., 2023),** we used BindCraft, an open-source, automated pipeline for de novo protein binder design with a high success rate (**Pacesa et al., 2025**). To this end, we identified the Sushi-3 residues that are both most important and sufficient for interacting with TGFβ RII-ECD. We examined each of the amino acid interacting pairs between Sushi-3 and TGFβ RII-ECD and selected pairs that were at less than 6Å of each other throughout the one-microsecond simulation, as well as were involved in the formation of a chemical bond with each other. We found five such interacting pairs that met both criteria. We found Sushi-3 V26 making a hydrophobic bond with TGFβ RII F11, Sushi-3 S40k, and S41 making a hydrogen bond with TGFβ RII R42 and E51, and Sushi-3 E53 making a salt-bridge with TGFβ RII R42 and K81 (**Figure 6A; Supplementary Figure 10**). Interestingly, these five residues are unique to Sushi-3, further substantiating that Sushi-3 showed the most potent interaction with TGFβ RII ECD (**Figure 5B**).

**Figure 6:**
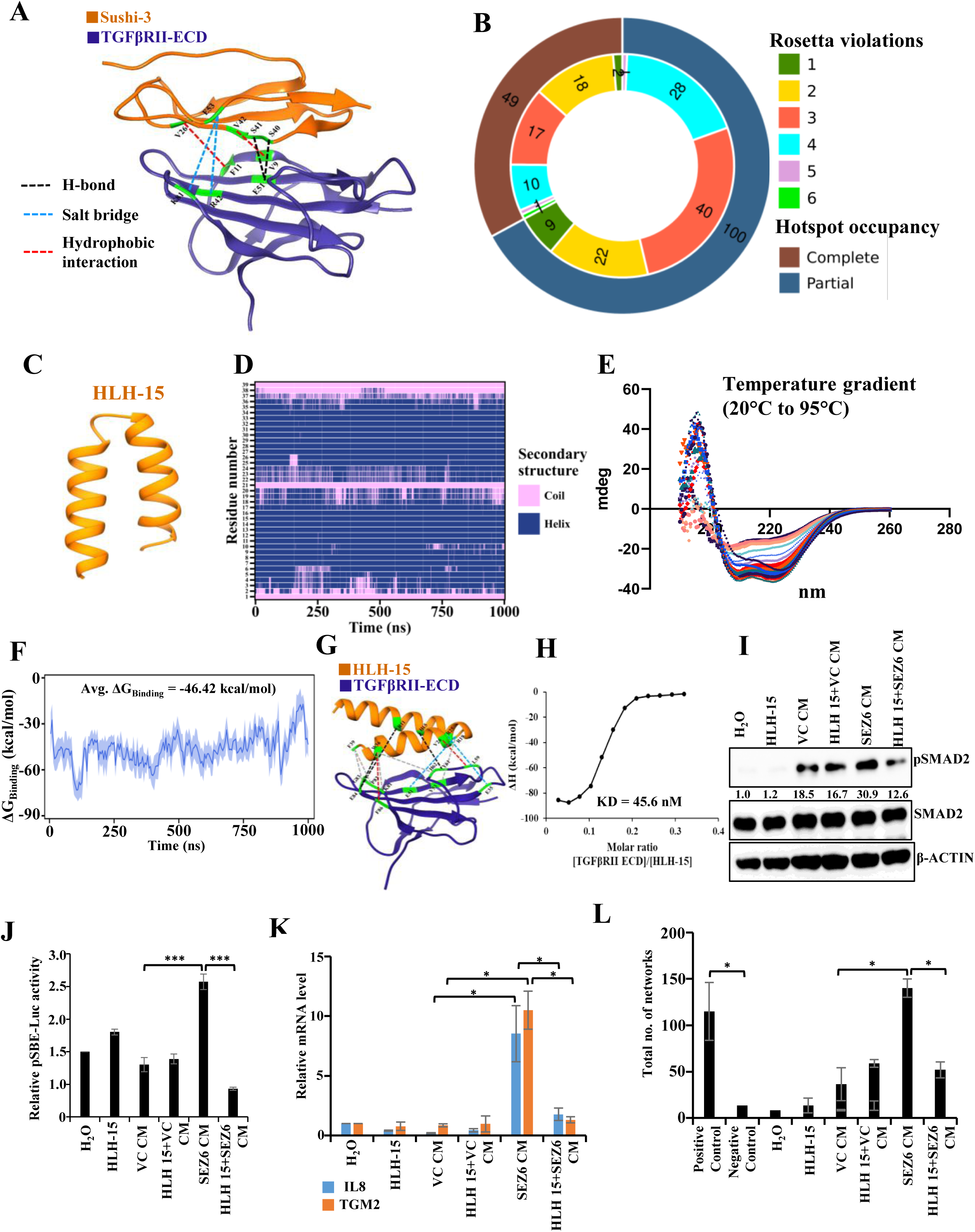
Elucidation of a de novo designed peptide inhibitor to perturb the SEZ6-mediated TGFβ pathway activation in endothelial cells. **(A)** Non-covalent interactions, including hydrogen bonds, salt bridges, hydrophobic contacts, and other interactions in the Sushi3-TGFβ RII ECD interface, identified from 1000 ns MD trajectories using PyContact. **(B)** Nested donut chart displaying the distribution of 149 on-site peptides satisfying all six AlphaFold2-derived parameters, classified by hotspot residues occupancy status (outer space ring) and Rosetta-based metric violations (inner space ring) **(C)** Secondary structure representation of HLH-15 peptide showing intact alpha-helical structure upon 1000 ns unbound MD simulation. **(D)** DSSP profiles of HLH-15 from a 1000 ns unbound MD simulation, confirming stable α-helical secondary structure independent of TGFβ RII ECD interaction. **(E)** Circular Dichroism (CD) profile of HLH-15 characteristic of α-helices, validating stable secondary structural integrity across the given temperature gradient (20°C to 95°C). **(F)** Binding affinity assessment of HLH-15 to TGFβ RII-ECD by MM-PBSA-derived binding free energies (kcal/mol) computed across 1000 ns MD simulations. **(G)** Non-covalent interactions, including hydrogen bonds (dotted black line), salt bridges (dotted blue line), hydrophobic contacts (dotted red line), and other interactions (dotted grey line) in the HLH-15-TGFβ RII ECD interface, were identified from 1000 ns MD trajectories using PyContact. **(H)** Determination of experimental dissociation constant (K_D_) value of HLH-15 with bacterially purified TGFβ RII-ECD using isothermal titration calorimetry (ITC). **(I)** Western blot showing the impact of pre-treatment of ST1 cells with HLH-15, followed by subsequent treatment of VC CM and SEZ6 CM, on the pSMAD2 protein level. The numbers indicate densitometric quantification of pSMAD2 bands using Image Lab (BIORAD) and normalized to total SMAD2. **(J)** Bar graph showing the impact of pre-treatment of ST1 cells with HLH-15, followed by subsequent treatment of VC CM and SEZ6 CM on the relative luciferase activity in SBE-Luc transfected ST1 cells. p-value was calculated by Student T-test, and <0.05 was considered significant with *, **, and *** representing p-values <0.05, 0.01, and 0.001, respectively. ns, non-significant. **(K)** Bar graph showing the impact of pre-treatment of ST1 cells with HLH-15, followed by subsequent treatment of VC CM and SEZ6 CM on the relative mRNA level of IL8 and TGM2. p-value was calculated by Student T-test, and <0.05 was considered significant with *, **, and *** representing p-values <0.05, 0.01, and 0.001, respectively. ns, non-significant. **(L)** Bar graph showing the impact of pre-treatment of ST1 cells with HLH-15, followed by subsequent treatment of VC CM and SEZ6 CM on the in-vitro network-forming ability of ST1 cells. p-value was calculated by Student T-test, and <0.05 was considered significant with *, **, and *** representing p-values <0.05, 0.01, and 0.001, respectively. ns, non-significant.

To design de novo peptide binders that interfere with the Sushi-3/TGFβ RII interaction, we used a design rationale based on the BindCraft pipeline (**Pacesa et al., 2025**), which we specifically adapted and optimized for short peptide design. We implemented BindCraft’s three-stage pipeline, involving AlphaFold2-based hallucination via backpropagation to simultaneously optimize the binder sequence and structure against the target, followed by MPNNsol-based sequence refinement of non-interface residues to improve solubility, and a final re-evaluation using AlphaFold2 monomer and Rosetta-based metrics to select high-confidence designs. TGFβ RII ECD (PDB ID: 1KTZ) was used as the structural template, with the surface containing the five residues V9, F11, R42, E51, and K81 of TGFβ RII, which consistently interacted with Sushi-3 in MD simulations, selected as the peptide-binding region. Cysteines were excluded in the peptide binders to avoid undesired disulfide formation, and design parameters were further constrained to favor predominantly α-helical peptides (>90% helical content) to promote structural stability. α-helices are among the most frequent secondary structural elements at protein–protein interfaces, making helical scaffolds particularly well suited to recapitulating native binding interactions (**Efimov, 1991; Braisted and Wells, 1996; Bullock et al., 2011**). Other parameters of peptide design are described in the **Supplementary section**. Of the 500 designed peptides, 259 showed on-site binding and 241 off-site binding. Among 259 on-site binders, 149 satisfied all six Alphafold2-derived quality parameters, of which 49 fully occupied the target hotspot residues across the entire binding interface and were selected for further analysis. (**Figure 6B; Supplementary Figure 11A**). These peptides were analyzed for their interaction with TGFβ RII ECD by all-atom MD simulations (1 microsecond), which identified 16 stably bound peptides (**Supplementary Figure 11B).** Subsequent simulations of these 16 peptides in isolation revealed that 15 retained their native secondary structures throughout, demonstrating strong intrinsic folding propensities independent of receptor context. These 15 peptides exhibited binding free energies ranging from -30 to -51 kcal/mol, indicating strong binding to TGFβ RII (**Supplementary Figure 11C)**. All 15 peptides adopted helix-loop-helix architectures, a common motif in protein-protein interfaces **(Murre et al., 2019)**. Six of these peptides exhibited coiled-coil heptad repeats, indicating stable hydrophobic cores that enhance structural rigidity **(Supplementary Figure 11C)**. The extended α-helical interface of the coiled-coil (helix-hairpin) scaffold provides a large solvent-accessible surface that is well suited for targeting protein–protein interactions (PPIs), which typically involve broad, relatively flat binding interfaces lacking the well-defined clefts or grooves readily accessible to conventional small-molecule inhibitors (**Khatri et al., 2022**). Among all peptides exhibiting stable secondary structure and coiled-coil architecture, the three most favorable by binding free energy (HLH-4, HLH-13, HLH-15) were chosen for experimental validation.

All three peptides adopted a stable α-helical secondary structure in solution, as indicated by MD simulations (Figure 6C and D; Supplementary Figure 12A, B, D, and E), which was further confirmed by Circular dichroism spectroscopy (**Figure 6E; Supplementary Figure 12C and F**). Isothermal titration calorimetry confirmed HLH-15 as the strongest binder with a K_D_ of 45.6 nM, consistent with its lowest MD simulation-based average binding free energy of -46.42 kcal/mol, among the three candidates **(Figure 6F, G, and H; Supplementary Figure 12G, H, I, J, K, and L)**. Inspection of the key non-covalent interactions between HLH-15 and TGFβ RII identified consistent contributions from hydrogen bonding (R11-E84, H15-Y61, and W36-E84), salt-bridge interactions (R19-E51 and R25-E35), hydrophobic packing (V26-L59 and W36-F86), and other interactions (R19-V53, W36-H62, W36-K81, F39-P82, and F39-G83) across the interface **(Figure 6G)**. HLH-15 peptide was chosen to test its ability to interfere with the ability of SEZ6 to activate the TGFβ pathway and promote angiogenesis. The CM containing SEZ6 failed to increase pSMAD2, luciferase activity from pSBE-Luc, transcript levels of IL8 and TGM2, or promote angiogenic networks in HLH-15 peptide pretreated ST1 endothelial cells (**Figure 6I, J, K, and L; Supplementary Figure 13A**). Collectively, these results demonstrate that HLH-15 was successfully designed and developed as a peptide binder for TGFβ RII, which effectively inhibits the SEZ6–TGFβ RII interaction, thereby abrogating TGFβ pathway activation and angiogenesis.

#### Novel BACE1 inhibitor abrogates SEZ6 shedding, activation of TGFB signaling, and angiogenesis, and glioma tumor growth

SEZ6 is a bona fide substrate of BAC1 (*b*eta-site *A*PP *c*leaving *e*nzyme *1*), which is well known for its crucial role in the pathogenesis of Alzheimer’s disease, as it cleaves the Amyloid precursor protein (APP) to generate amyloid-beta (Aβ) peptides (**Pigoni et al., 2016**; **Vassar et al., 1999**). Hence, based on the premise that inhibition of BACE1 would block SEZ6 shedding and thereby abrogate activation of the TGFβ pathway, angiogenesis, and glioma tumor growth, we embarked on developing novel BACE1 inhibitors. BACE1 is a type I transmembrane aspartyl protease homing Asp93 and Asp289 at its catalytic active site (**Zhao et al., 2020**). While the compound 89, developed by Hilpert et al. (2013) as a BACE1 inhibitor, was potent, it had several disadvantages, including large size and susceptibility to proteolytic cleavage (**Ghosh et al., 2012**). Retaining the pharmacophoric features of compound 89 and employing bioisosteric replacement of a 6-membered oxazine with a 5-membered pyrazole aryl, we developed S2C as a potent inhibitor of BACE1 (**Supplementary information for details**).

Molecular docking and molecular dynamics simulation study revealed that S2C interacts with BACE1 with a Glide docking score of -7.317 kcal/mol, positioning itself within the catalytic pocket and forming hydrogen bonds with Asp93, Gly95, and Thr293, along with a stabilizing π–π stacking interaction with Tyr132 **(Figure 7A and B)**. Subsequent MM-GBSA calculations on the equilibrated trajectory yielded a binding free energy of -37.0 kcal/mol, together supporting a favorable and stable binding mode of S2C within the BACE1 catalytic pocket.

**Figure 7:**
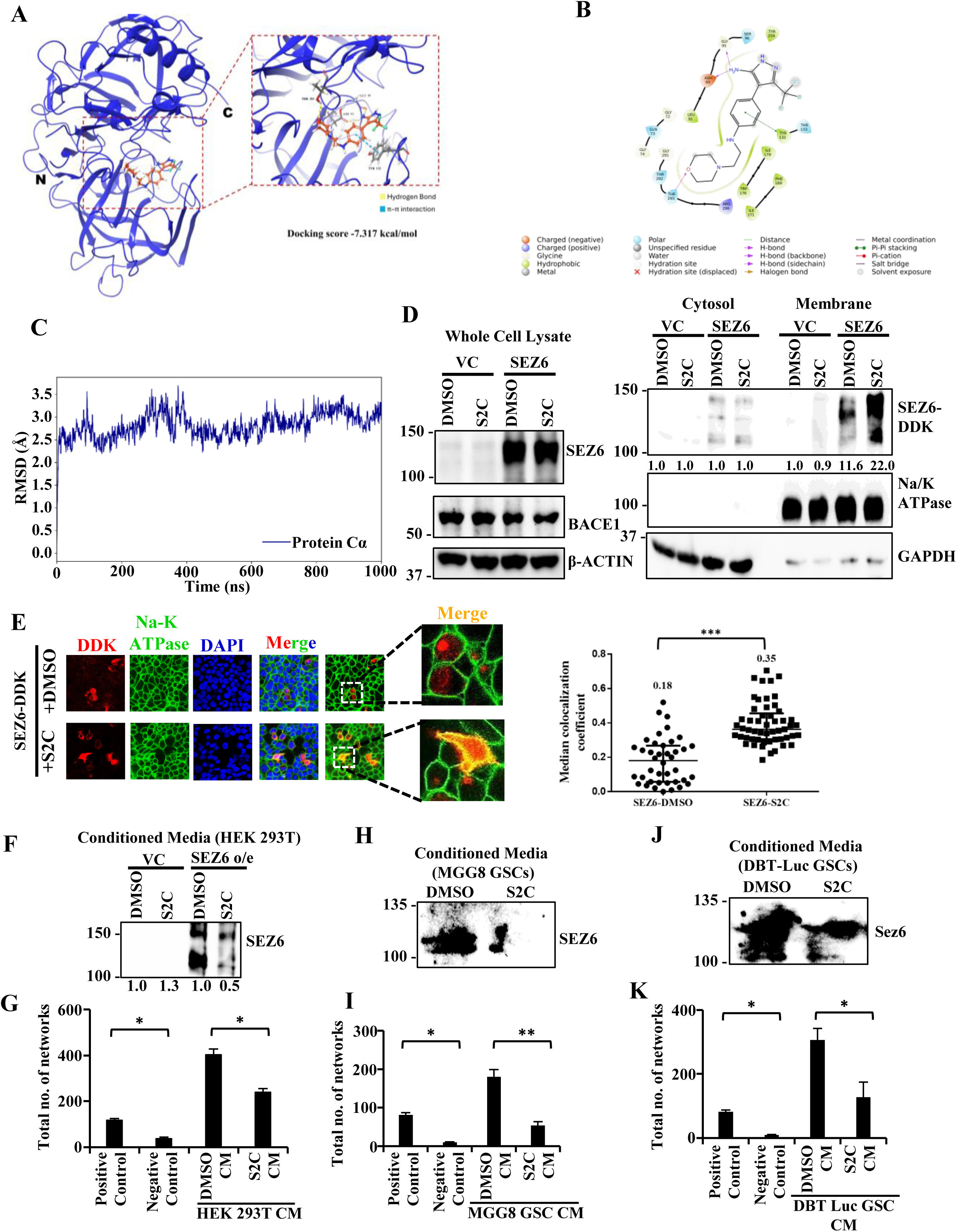
S2C, a novel small-molecule inhibitor of BACE1, inhibits SEZ6 shedding, thereby disrupting SEZ6-mediated TGFβ pathway activation in endothelial cells. **(A)** Molecular docking analysis of compound S2C in the active site of BACE1. Cartoon representation of the BACE1-S2C complex showing the overall binding orientation of S2C (orange sticks) within the catalytic pocket. The N- and C-terminal regions of BACE1 are indicated. Magnified view of the binding pocket depicting the key interacting residues and hydrogen-bond (yellow dashed lines) and π–π interactions (blue dashed lines) stabilizing S2C **(B)** Two-dimensional interaction map illustrating the hydrogen bonds, π–π stacking, and hydrophobic interactions between S2C and the active-site residues of BACE1. **(C)** Root-mean-square deviation (RMSD) of the BACE1–S2C complex during the 1000 ns molecular dynamics simulation. **(D)** Western blot showing the whole cell lysate along with the corresponding cytosolic and membrane fractions upon treatment of SEZ6-overexpressing HEK-293T cells with S2C compared to DMSO. The numbers indicate densitometric quantification of SEZ6 bands using Image Lab (BIORAD) and normalized to the respective loading controls. **(E)** Representative confocal images showing the membrane colocalization of SEZ6 (red) and Na-K ATPase (green, endogenous membrane marker) upon treatment of SEZ6-overexpressing HEK-293T cells with S2C compared to DMSO. The corresponding graph shows the quantification of the colocalization coefficient between SEZ6 and Na-K ATPase under DMSO and S2C treatment conditions. Magnification 63X. **(F)** Western blot showing the shed-SEZ6 in the CM upon treatment of SEZ6-overexpressing HEK-293T cells with S2C compared to DMSO. **(G)** Bar graph showing the impact of S2C-treated HEK-293T-derived CM on the in vitro network formation by ST1 cells, compared to DMSO-treated HEK-293T-derived CM. **(H)** Western blot showing the shed-SEZ6 in the CM upon treatment of MGG8 GSCs with DMSO versus S2C. **(I)** Bar graph showing the impact of S2C-treated MGG8-derived CM on the in vitro network formation by ST1 cells, compared to DMSO-treated MGG8-derived CM. **(J)** Western blot showing the shed-Sez6 in the CM upon treatment of DBT Luc GSCs with DMSO versus S2C. **(K)** Bar graph showing the impact of S2C-treated DBT Luc-derived CM on the in vitro network formation by ST1 cells, compared to DMSO-treated DBT Luc-derived CM.

The structural stability of the BACE1–S2C complex was further assessed by RMSD analysis of the protein Cα atoms across a 1000 ns molecular dynamics simulation **(Figure 7C)**. Following an initial equilibration phase (2.4-2.6 Å within the first few nanoseconds), the complex remained stable up to ∼600 ns, fluctuating between 2.2 and 3.6 Å (average ∼2.8 Å), with a transient increase (3.4-3.7 Å) between 250–380 ns that resolved by ∼450 ns, indicating a reversible conformational adjustment rather than instability. A modest upward shift (2.8–3.3 Å) after ∼700 ns reflected a stable conformational rearrangement rather than unfolding or ligand dissociation, confirming that the BACE1–S2C complex remained structurally stable throughout the full simulation. Per-residue RMSF analysis showed that most BACE1 residues fluctuated below 1.0 Å, with flexibility concentrated at the N- and C-termini (3.3-4.0 Å) and select loop regions (residues 50, 77, 115, 165, 250-270, and 310), of which residue 165 showed the greatest mobility (∼3.9 Å) **(Supplementary Figure 14)**. Most importantly, active-site residues and the protein core remained rigid (RMSF < 1.0 Å) throughout, indicating that S2C engagement preserved the structural integrity of the ligand-binding pocket.

To evaluate the drug-likeness of S2C, we predicted its ADME properties using QikProp **(Jorgensen and Duffy, 2002)**. S2C (molecular weight 355.4 g/mol) satisfied the Lipinski rule of five, with a favorable predicted lipophilicity profile (QPlogPo/w = 2.05), and both hydrogen-bond donor (4) and acceptor (6.2) count within recommended limits **(Lipinski et al., 2001)**. Additional descriptors, including octanol/gas (QPlogPoct = 21.08) and water/gas (QPlogPw = 13.90) partition coefficients, also fell within acceptable ranges. Remarkably, the predicted brain/blood partition coefficient (QPlogBB = -0.36) fell within the CNS-favorable range of - 3.0 to 1.2, indicating that S2C is predicted to cross the blood–brain barrier, a key pharmacokinetic requirement for a centrally acting BACE1 inhibitor targeting Alzheimer’s disease. Low predicted plasma protein binding (QPlogKhsa = -0.03) further supported favorable systemic free-drug availability. Overall, these results demonstrate that S2C exhibits a stable, favorable binding mode within the BACE1 active site and possesses drug-like physicochemical and CNS-penetrant properties consistent with its intended therapeutic application.

Next, we tested S2C’s ability to inhibit SEZ6 shedding and abrogate SEZ6-induced activation of the TGFβ pathway, angiogenesis, and glioma tumor growth. In HEK-293T cells transfected with SEZ6, pretreatment with S2C substantially increased the membrane-localized SEZ6 with no significant difference in cytosolic and overall content of SEZ6 (**Figure 7D and E**). This was further confirmed by confocal microscopy, in which SEZ6 showed a significantly higher co-localization with Na/K-ATPase, a membrane protein, in S2C pretreated HEK-293T cells (**Figure 7F**). The CM collected from S2C pretreated HEK-293T cells, MGG8 GSC cells, and DBT-Luc (murine GSC line) showed significantly reduced angiogenic network formation by ST1 endothelial cells (**Figure 7G, H, and I; Supplementary Figure 15A, B, and C**). Collectively, these results identify S2C as a robust and effective BACE1 inhibitor that abrogates SEZ6 shedding, thereby blocking SEZ6-driven activation of the TGFβ pathway and the downstream pro-angiogenic program. Experiments are ongoing to demonstrate that S2C pretreatment inhibits glioma tumor growth in mouse glioma models.

## Discussion

This study identifies SEZ6 as a GSC-specific molecule that promotes glioma tumor growth by inducing tumor angiogenesis. SEZ6 shed into the stroma upon BACE1 cleavage activates IL8 via TGFβ signaling, promoting angiogenesis. Sushi 3 domain of SEZ6 mediates the interaction with TGFβ RII. Either treatment with a small-molecule inhibitor of BACE1, which increases the retention of SEZ6 on the membrane, or a novel de novo-designed peptide binder that efficiently interrupts the interaction between SEZ6 and TGFβ RII, suppressed glioma tumor growth by abrogating SEZ6-induced TGFβ pathway and tumor angiogenesis.

In addition to malignant cells, the tumor microenvironment (TME) is a dynamic ecosystem comprising immune cells, cancer-associated fibroblasts, endothelial cells, and the extracellular matrix (**Lorenc et al., 2024**). Among these, tumor endothelial cells play a central role by supplying oxygen and nutrients essential for tumor growth. While oncogenic drivers initiate tumorigenesis, sustained tumor progression depends on continuous angiogenesis; without it, tumors experience hypoxia, leading to necrosis or apoptosis. As angiogenesis is largely driven by tumor-derived signals, significant efforts have focused on identifying pro- angiogenic factors and the mechanisms activating endothelial cells, with the goal of developing targeted anti-angiogenic therapies. This work identified SEZ6 as a potent inducer of angiogenesis.

Cancer stem cells (CSCs), a small subpopulation within tumors, possess the unique ability to self-renew, initiate tumor formation, and differentiate into heterogeneous tumor cell populations (**Loh and Ma, 2024**). In glioblastoma (GBM; grade IV glioma), the most aggressive primary brain tumor in adults (**Stupp et al., 2005**), glioma stem-like cells (GSCs) represent the CSC compartment and are major drivers of tumor aggressiveness, therapeutic resistance, and recurrence (**Visvanathan et al., 2017**). Emerging evidence indicates that GSCs reside in specialized perivascular niches, where they engage in bidirectional interactions with endothelial cells to promote tumor angiogenesis and maintain stemness (**Calabrese et al., 2007; Prager et al., 2020).** Through the secretion of pro-angiogenic factors, GSCs actively contribute to the formation of aberrant vasculature, which in turn supports tumor growth and survival. Although vascular endothelial growth factor A (VEGF-A) has been considered a central mediator of tumor angiogenesis, clinical strategies targeting VEGF signaling have yielded limited success in GBM. Notably, Bevacizumab (Avastin), a monoclonal antibody against VEGF-A, failed to improve overall survival in Phase III clinical trials (**Gilbert et al., 2014; Chinot et al., 2014**), highlighting the redundancy and complexity of angiogenic pathways in GBM. These observations underscore the need to identify alternative angiogenic mediators, particularly those specifically expressed or regulated by GSCs. Targeting such GSC-derived factors may provide more effective strategies to disrupt tumor vascularization and overcome resistance to current anti-angiogenic therapies. This study, through an integrated analysis, identified that SEZ6 is specifically expressed by GSCs. The SEZ6 expression is controlled by Wnt/βcatenin pathway. GSC maintenance has been shown to require an activated Wnt signaling in GSCs (**Rheinbay et al., 2013; Suva et al., 2014**). Our results indicate that SEZ6 expression and induced angiogenesis are controlled by the Wnt/β-catenin pathway, as pharmacological inhibition of β-catenin reduced SEZ6 expression and the ability of GSC CM to promote angiogenesis.

SEZ6, a membrane protein, is specifically expressed in the brain, and its expression was found to be increased upon treatment with a convulsant drug (**Shimizu-Nishikawa et al., 1995**). SEZ6 protein contains five copies of short consensus repeat (SCR; also called the Sushi domain) and two copies of CUB domain (Complement C1r/s-like repeat). Reports show that CUB domain-containing proteins activate EGFR, Sonic Hedgehog, and TGFβ signaling (**Law et al., 2016; Wierbowski et al., 2020; Yang et al., 2018; Wu et al., 2011**). Our study shows that SEZ6 binds to TGFβ RII and activates TGFβ signaling to promote angiogenesis. Interestingly, the Sushi 3 domain is found to mediate the interaction with TGFβ RII. Several reports document an important role for TGFβ signaling in vascular biology and angiogenesis (**Goumans et al., 2009; Walshe et al., 2009; Gong et al., 2025).** Our results showed that SEZ6 activates TGFβ signaling to induce angiogenesis. We also found that SEZ6-dependent TGFβ signaling induces IL8, which in turn promotes angiogenesis. Indeed, IL-8 has been shown to be a key mediator of angiogenesis (**Hou et al., 2014; Tan et al., 2014**). Further, we also demonstrate that IL8 is a bona fide target of the TGFβ pathway.

Our study has identified several promising avenues for developing SEZ6-targeted therapeutic strategies for glioblastoma (GBM). Notably, SEZ6 has been characterized as a physiological substrate of BACE1, a beta-site amyloid precursor protein (APP)-cleaving enzyme (**Pigoni et al., 2016**). BACE1-mediated proteolytic processing of SEZ6 is critical for its shedding into the extracellular tumor stroma, where it subsequently facilitates activation of the TGFβ signaling pathway, a key driver of tumor angiogenesis. Given this mechanistic link, BACE1 is an attractive, rational target for indirectly modulating SEZ6 function in the GBM microenvironment. To this end, we have developed S2C, a novel small-molecule inhibitor of BACE1, designed to effectively suppress its enzymatic activity. Inhibition of BACE1 by S2C is evidenced by increased membrane retention of SEZ6, indicating a blockade of its cleavage and release. Functionally, conditioned media collected from GSCs treated with S2C failed to activate the TGFβ pathway in endothelial cells and to promote angiogenic responses in vitro. These findings underscore the ability of S2C to disrupt SEZ6-dependent paracrine signaling mechanisms that are critical for vascular development in tumors. Furthermore, the therapeutic efficacy of S2C in suppressing glioma progression is currently being evaluated in an in vivo mouse model. Collectively, these results highlight the potential of targeting the BACE1–SEZ6 axis as an innovative strategy to attenuate tumor angiogenesis and improve treatment outcomes in GBM.

Given our findings that the interaction between SEZ6 and TGFβ RII is essential for SEZ6-mediated angiogenesis, disrupting this interaction represents a compelling therapeutic strategy for glioma. To this end, we systematically mapped the key amino acid residues mediating the SEZ6–TGFβ RII interface and leveraged this structural insight to guide the rational design of inhibitory molecules. Using the BindCraft computational pipeline (**Pacesa et al., 2025**), we generated a series of de novo peptide binders specifically engineered to interfere with this interaction. Among these candidates, the top-ranked peptide, HLH-15, demonstrated strong binding affinity toward TGFβ RII and formed a stable complex that effectively occluded SEZ6 engagement. Functionally, HLH-15 abrogated SEZ6-driven activation of the TGFβ signaling pathway and significantly impaired downstream angiogenic responses in vitro. These results underscore the therapeutic potential of targeting protein– protein interactions within the tumor microenvironment using rationally designed peptide inhibitors. Ongoing studies are currently evaluating the efficacy of HLH-15 in vivo, with particular emphasis on its ability to suppress glioma tumor growth in a mouse model. Collectively, this work highlights a novel, mechanistically informed approach to targeting the SEZ6 axis, offering a promising avenue for developing next-generation anti-angiogenic therapies for glioma.

Since our results show that the SEZ6/TGFβ RII interaction is essential for SEZ6-induced angiogenesis, a peptide that could prevent this interaction could be an attractive therapeutic for glioma. We mapped the interacting amino acid residues between SEZ6 and TGFβ RII and used this information to design de novo peptide binders with the BindCraft pipeline (**Pacesa et al., 2025**) to inhibit SEZ6/TGFβ RII interaction. The top-ranking peptide, HLH-15, bound to TGFβ RII, forming a stable complex that inhibited SEZ6’s ability to activate the TGFβ pathway and promote angiogenesis. Experiments are currently being conducted to determine HLH-15’s ability to inhibit glioma tumor growth in a mouse glioma model.

Overall, this study identifies SEZ6 as a GSC-specific molecule that plays a pivotal role in promoting tumor angiogenesis. By delineating its functional significance within the tumor microenvironment, we highlight SEZ6 as a key driver of vascular development in glioma progression. Importantly, multiple SEZ6-targeted therapeutic strategies have been developed to exploit this vulnerability, including a novel BACE1 inhibitor and a rationally designed de novo peptide binder that specifically disrupts the SEZ6-TGFβ receptor II (TGFβ RII) interaction. These targeted interventions have demonstrated significant efficacy in inhibiting SEZ6-mediated signaling pathways in endothelial cells, thereby attenuating angiogenic processes in vitro. The ability of these approaches to interfere with both molecular signaling and downstream functional outcomes underscores their therapeutic potential. Collectively, these findings provide compelling evidence that SEZ6, selectively expressed in GSCs, is a promising, previously underexplored therapeutic target. Targeting SEZ6 may offer a novel and effective strategy to limit tumor vascularization and ultimately improve treatment outcomes in glioma.

## Abbreviations

GBM: Glioblastoma
GSC: Glioma Stem-like cells
DGC: Differentiated Glioma cells
SEZ6: Seizure-related gene 6
BACE1: Beta-site APP cleaving enzyme 1

## Acknowledgments

We acknowledge the shRNA consortium (Dr. Subba Rao) at IISc, India, for the shRNA constructs. We thank Prof. Shashank Deep, IIT Delhi, for providing the plasmid for the pET32a-TGFβ RII ectodomain plasmid. The results published here are, in whole or in part, based upon data generated by The Cancer Genome Atlas (TCGA) pilot project established by the National Cancer Institute (NCI) and the National Human Genome Research Institute (NHGRI). Information about TCGA and the investigators and institutions that constitute the TCGA research network can be found at http://cancergenome.nih.gov. KS acknowledges CEFIPRA, DBT, DST-SERB, and DST-ANRF (Govt. of India) for research grants. Infrastructure supported by DST-FIST, the DBT-IISc partnership program, and the UGC is acknowledged.

---

**Supplementary Figure 1.**
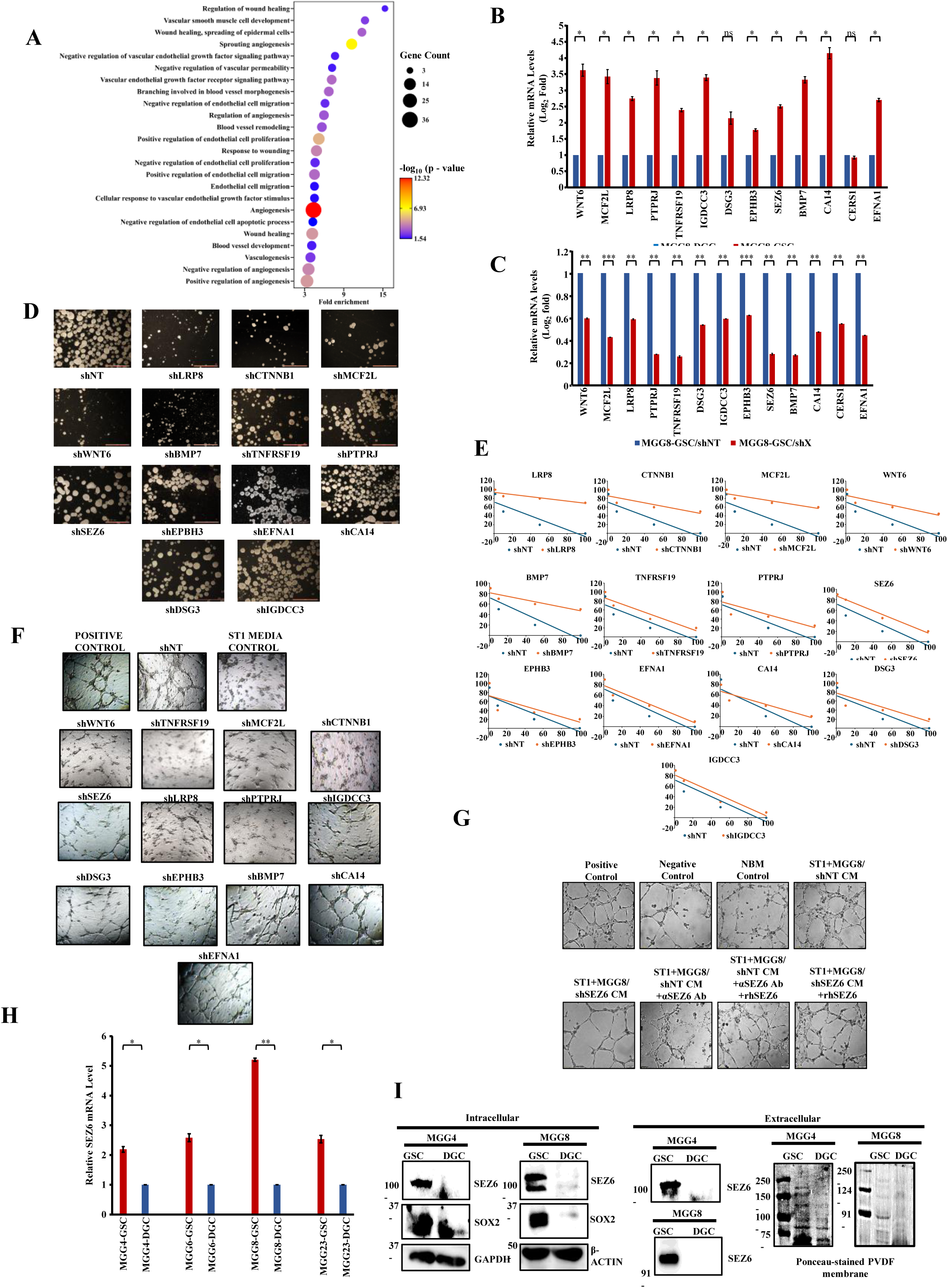

**Supplementary Figure 2.**
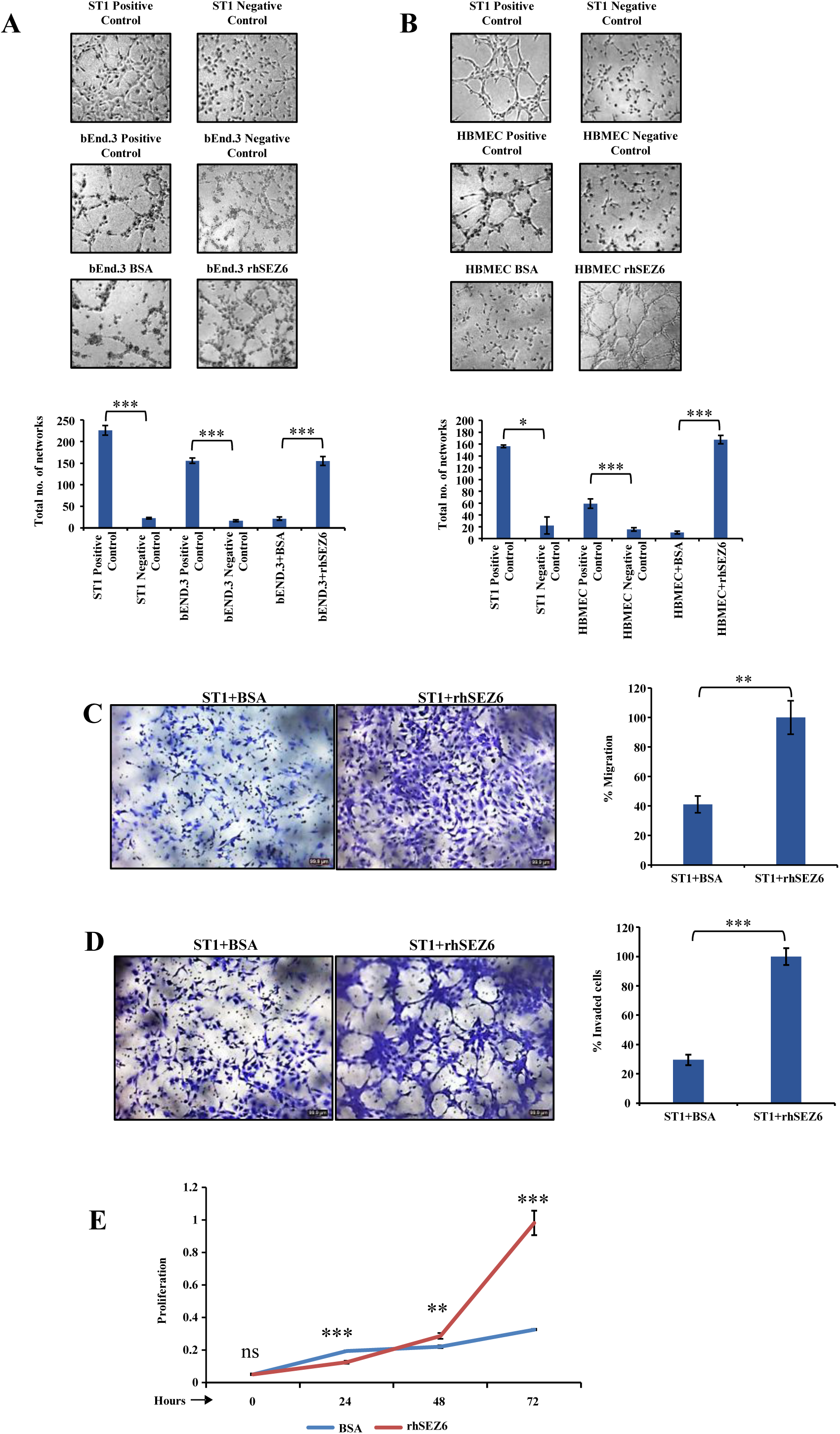

**Supplementary Figure 3.**
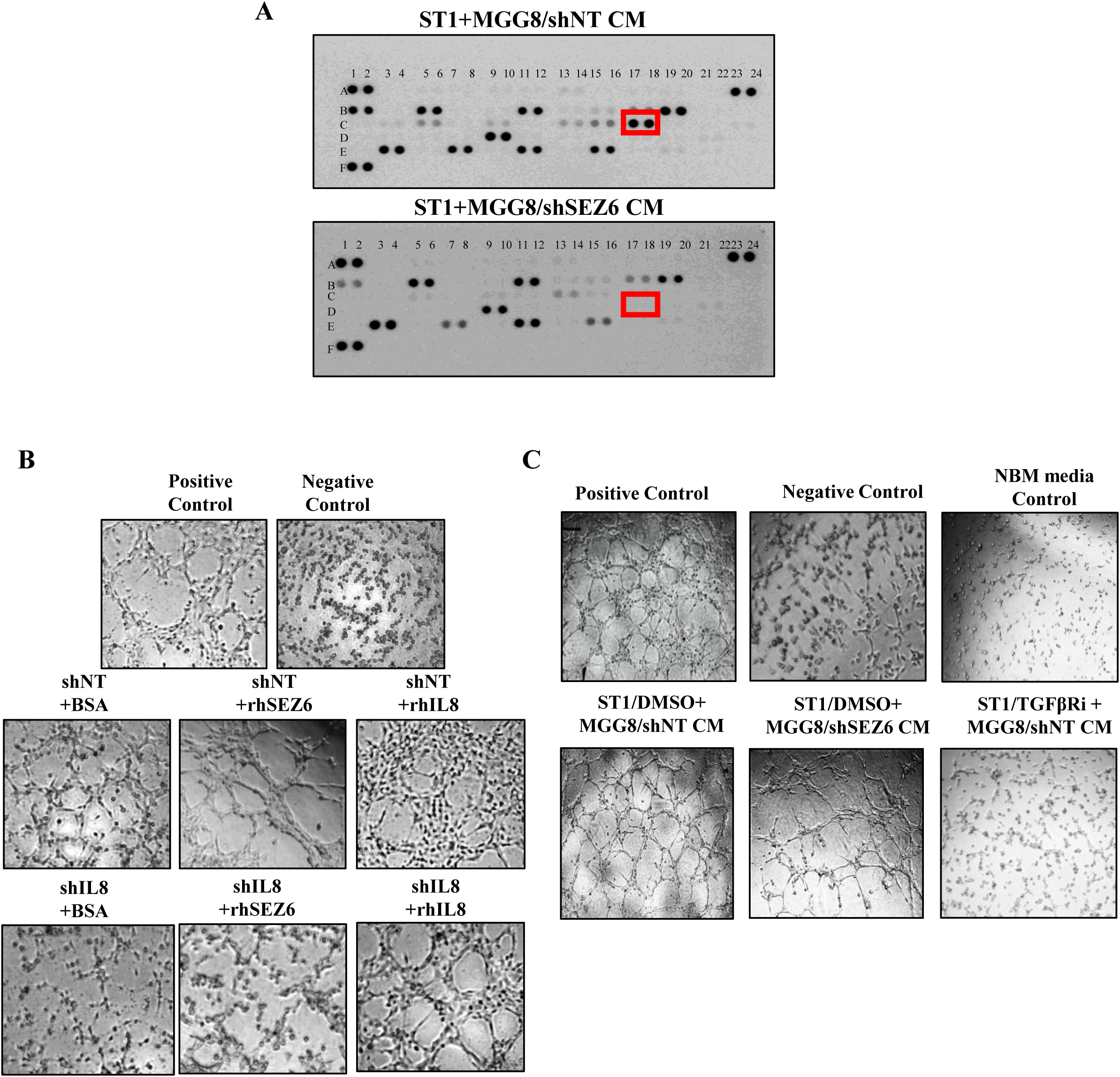

**Supplementary Figure 4.**
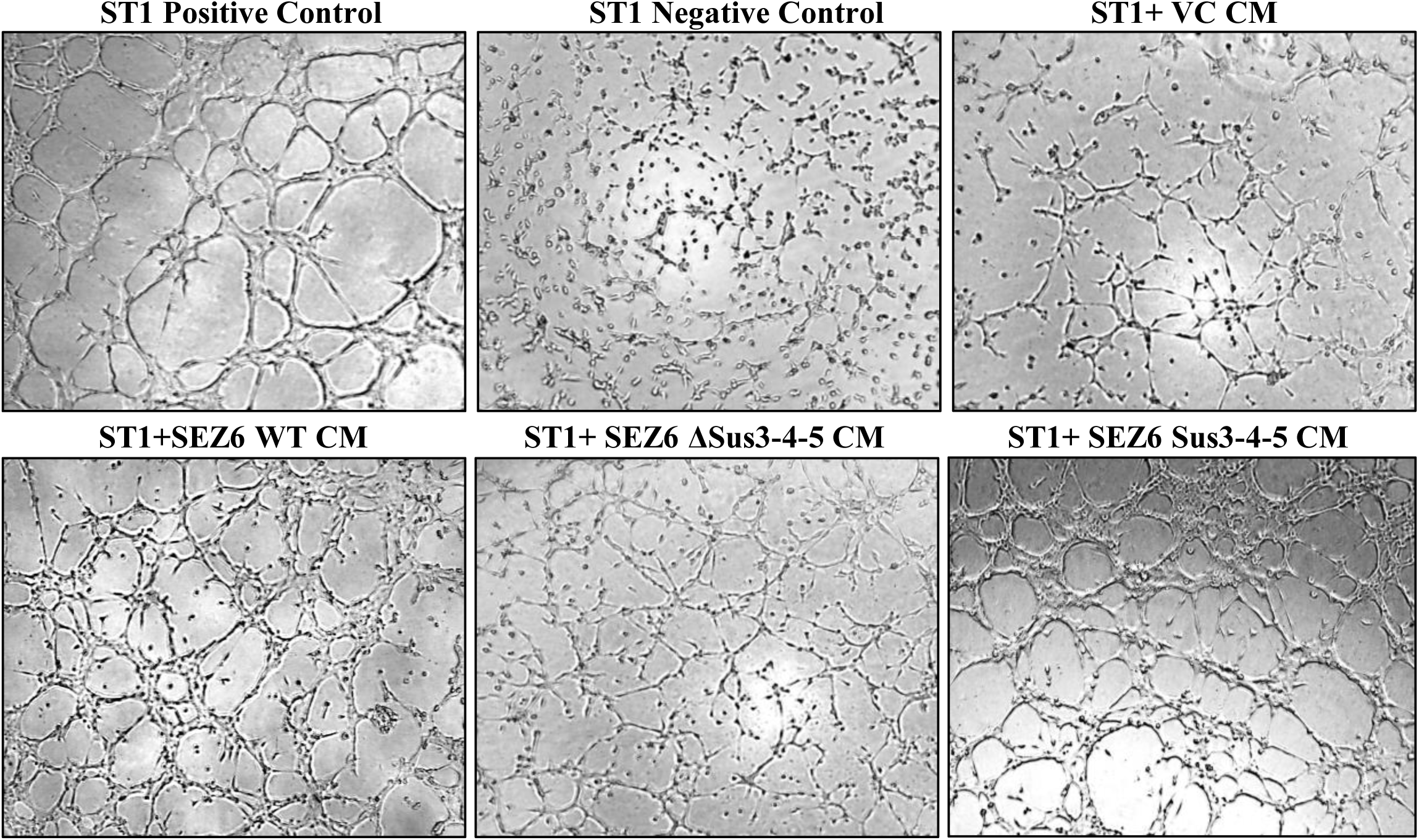

**Supplementary Figure 5.**
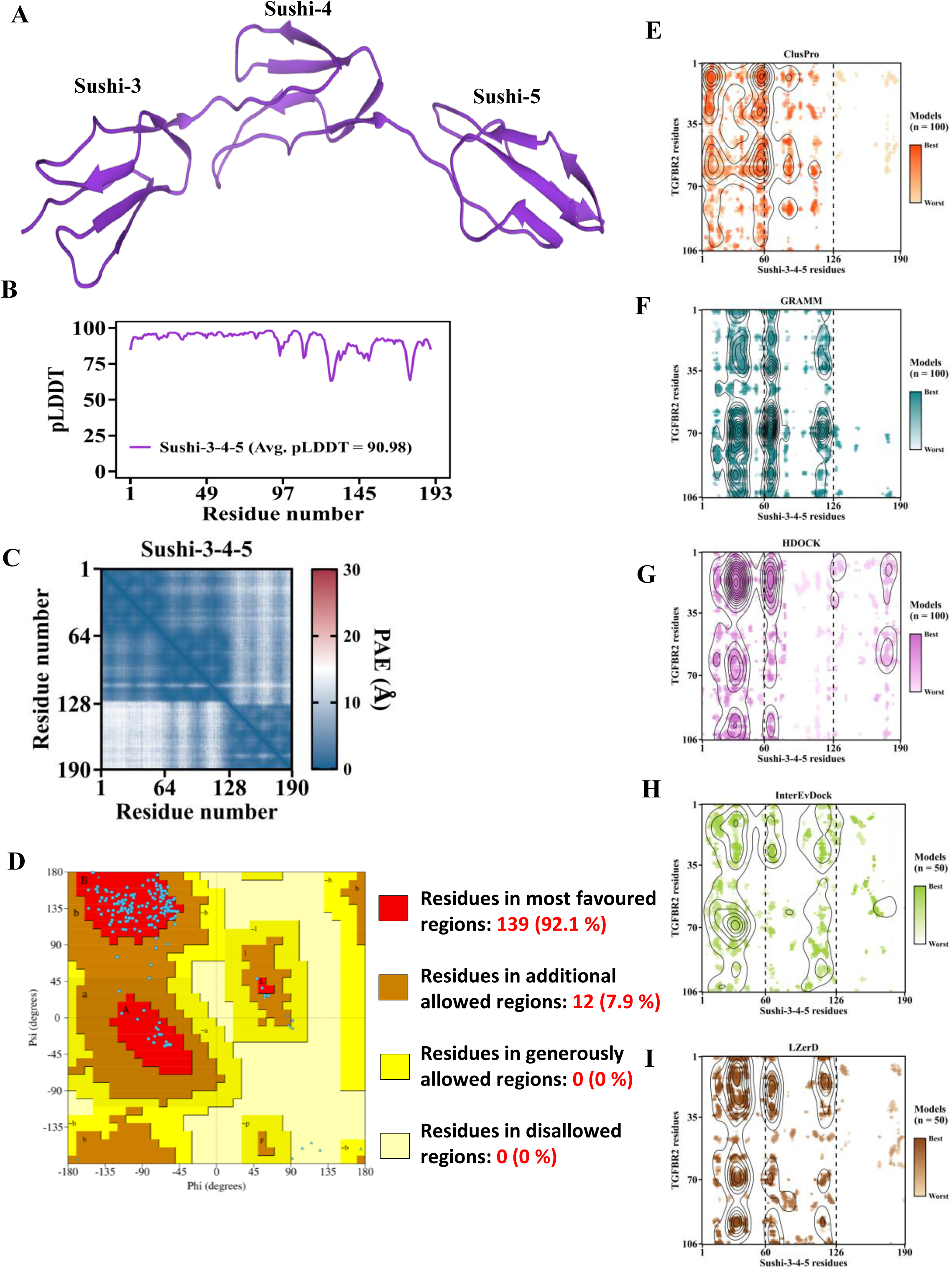

**Supplementary Figure 6.**
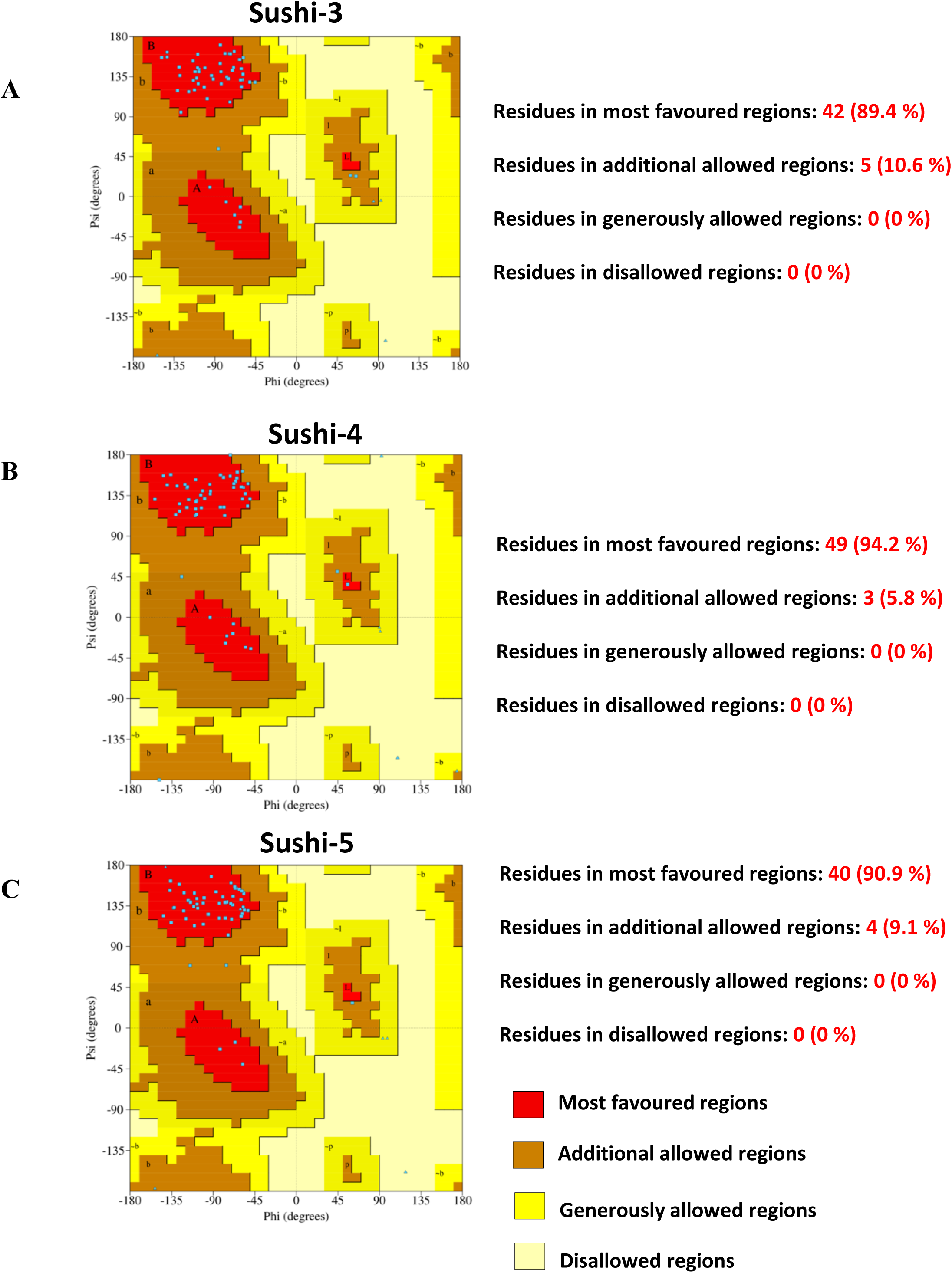

**Supplementary Figure 7.**
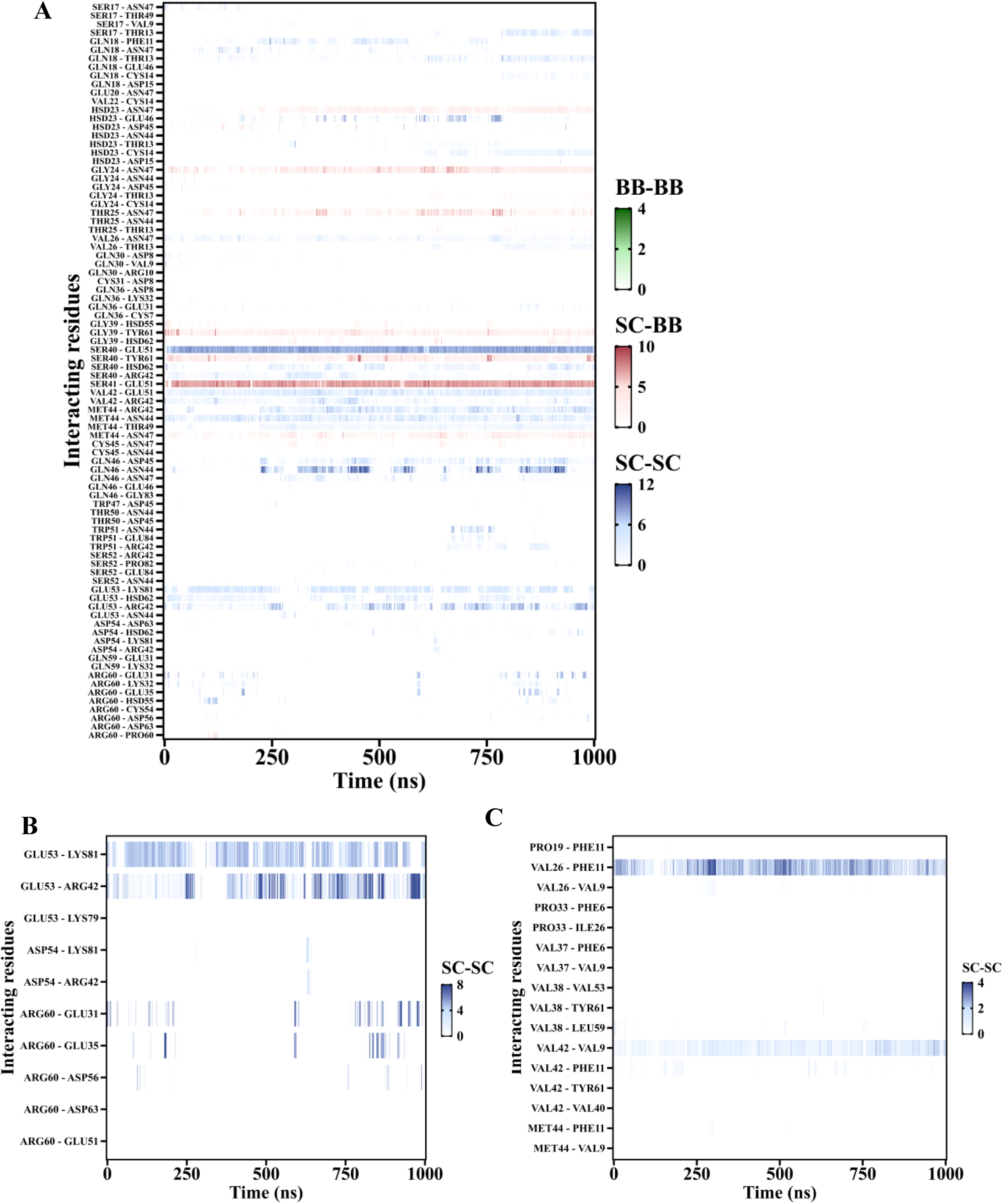

**Supplementary Figure 8.**
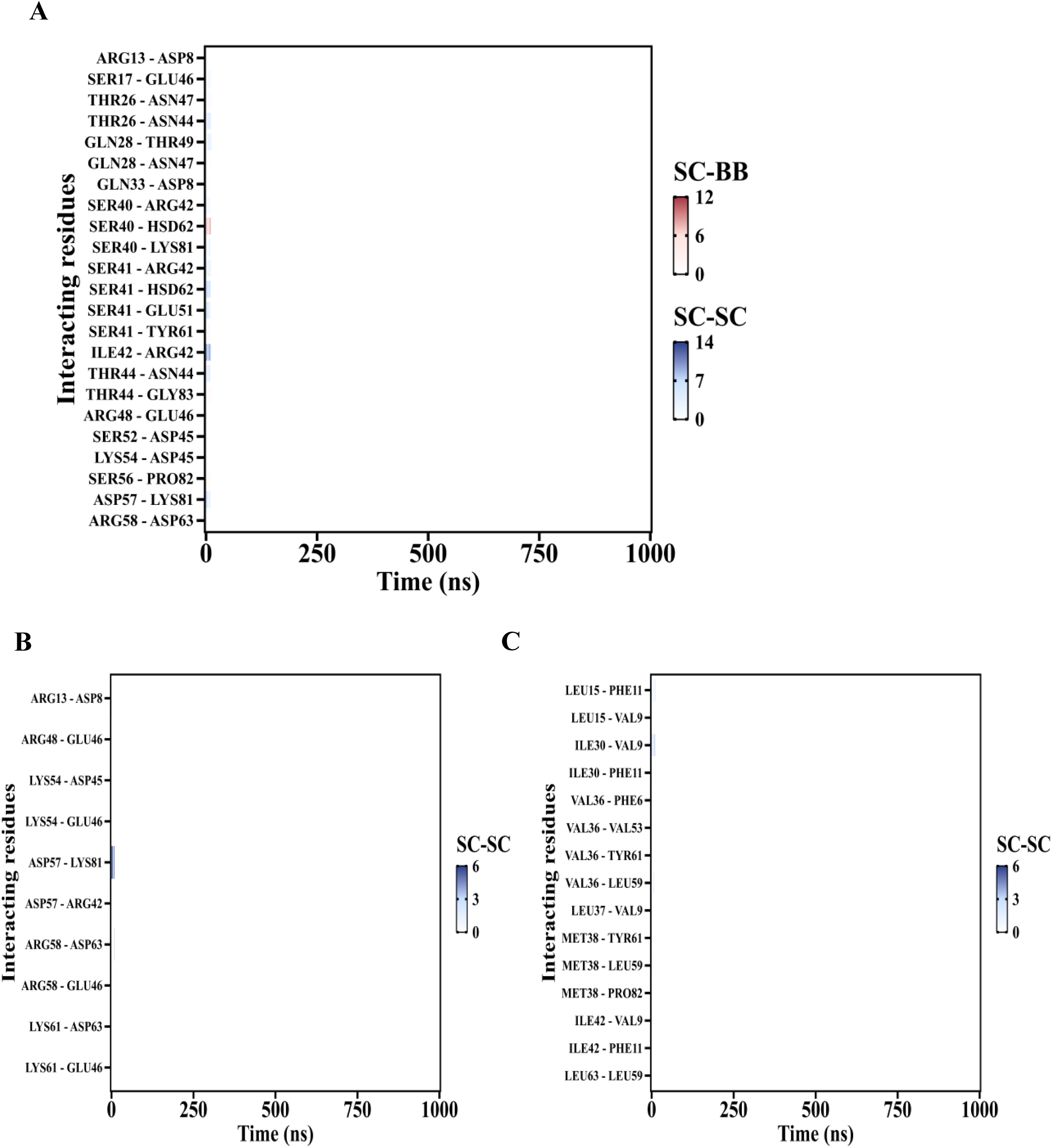

**Supplementary Figure 9.**
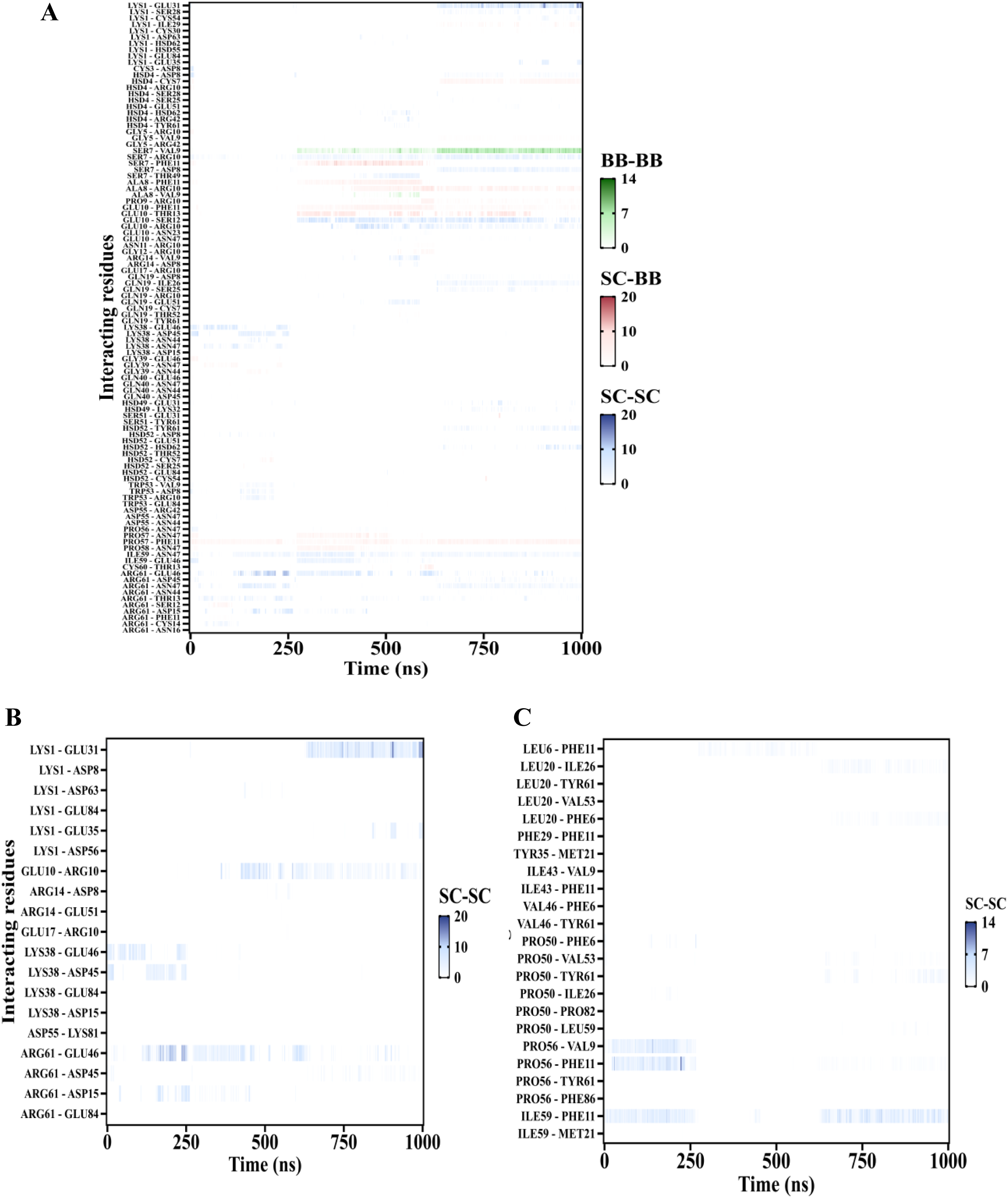

**Supplementary Figure 10.**
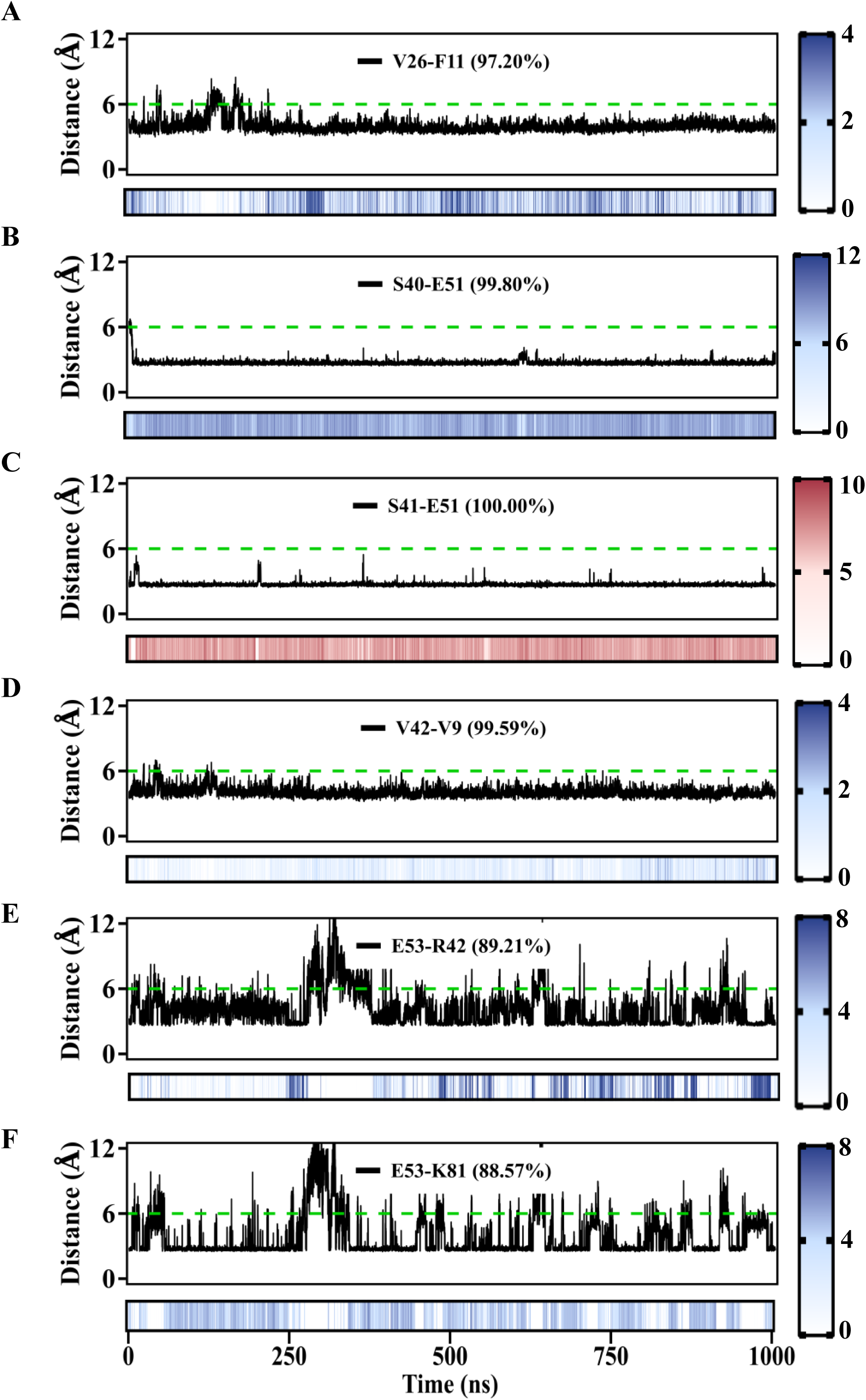

**Supplementary Figure 11.**
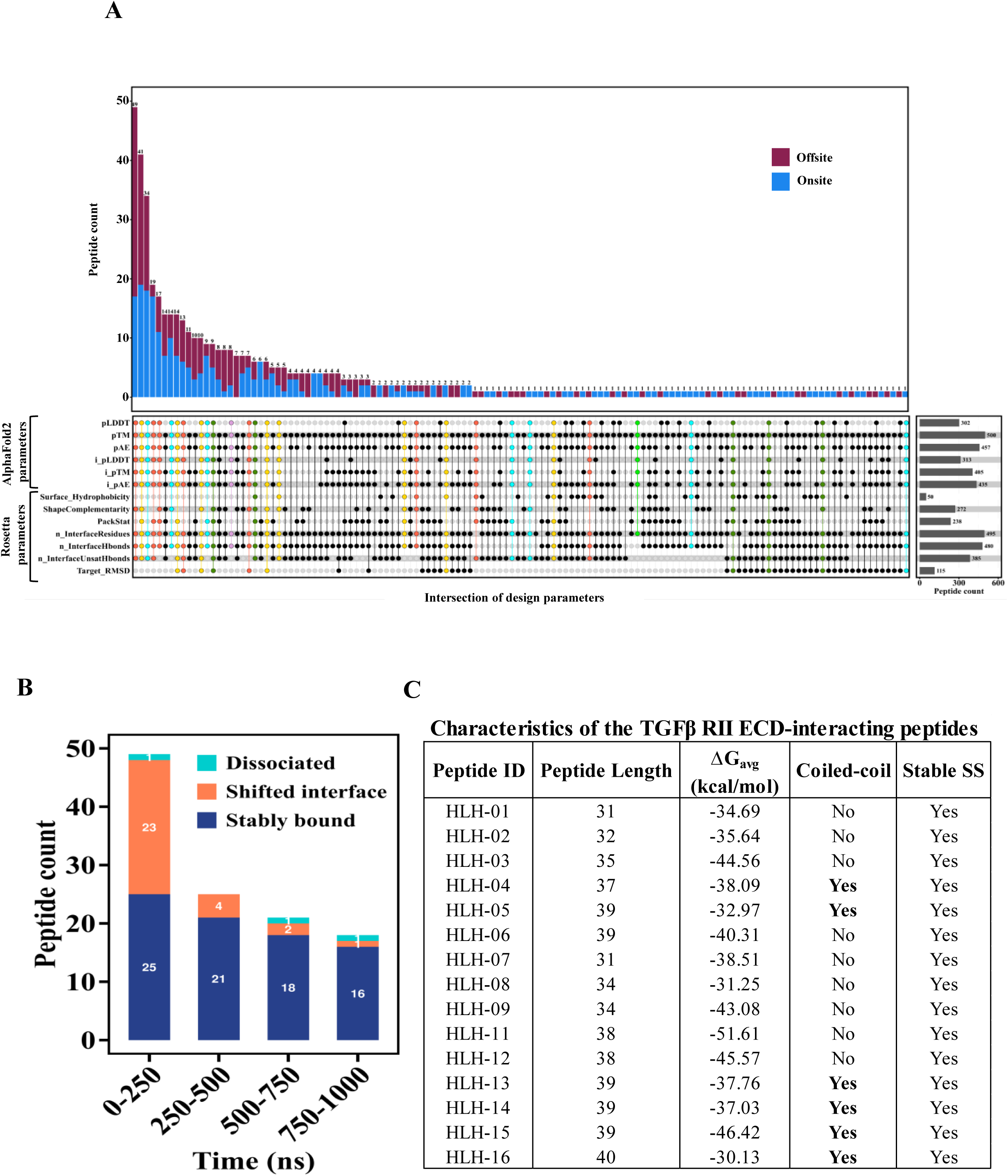

**Supplementary Figure 12.**
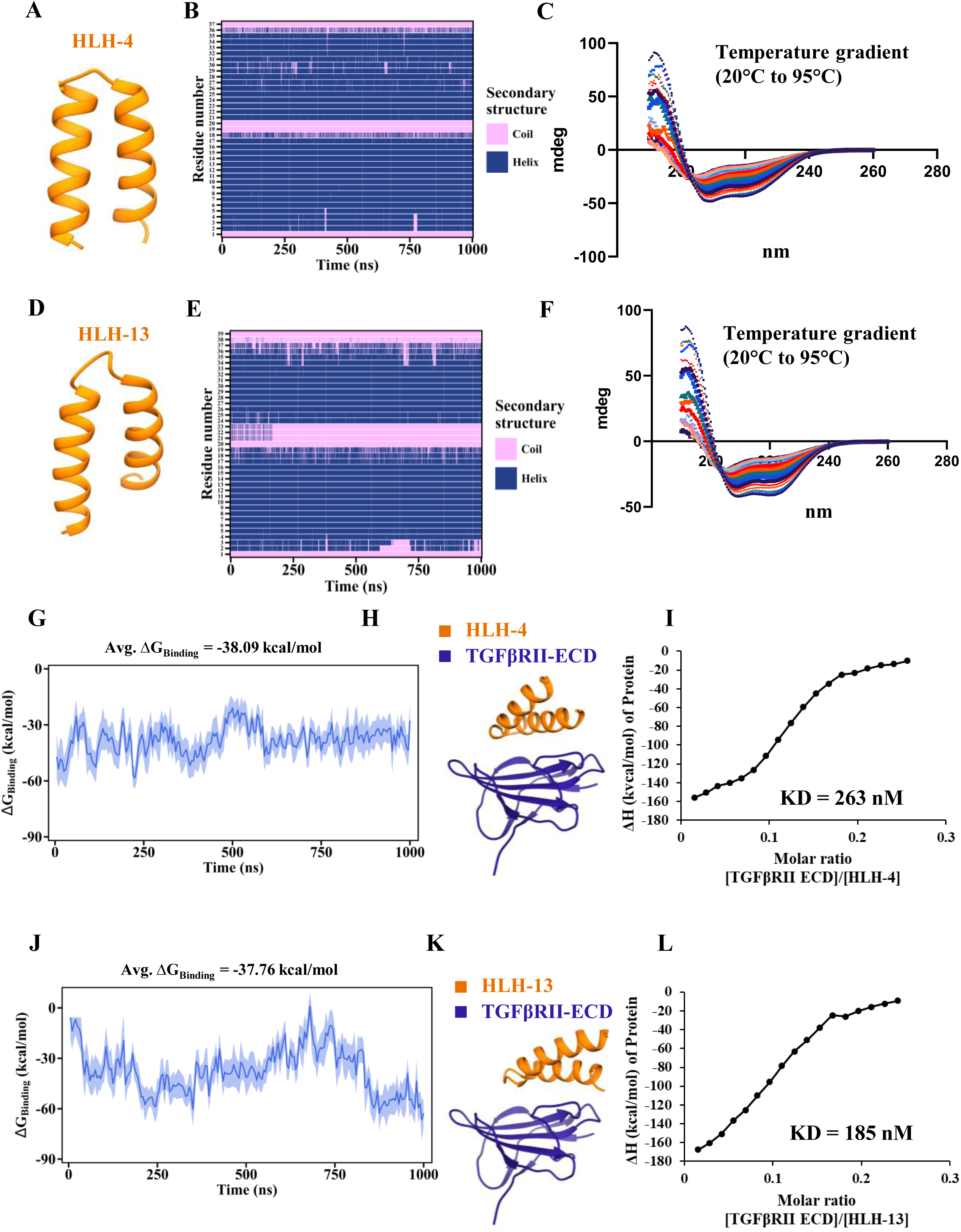

**Supplementary Figure 13.**
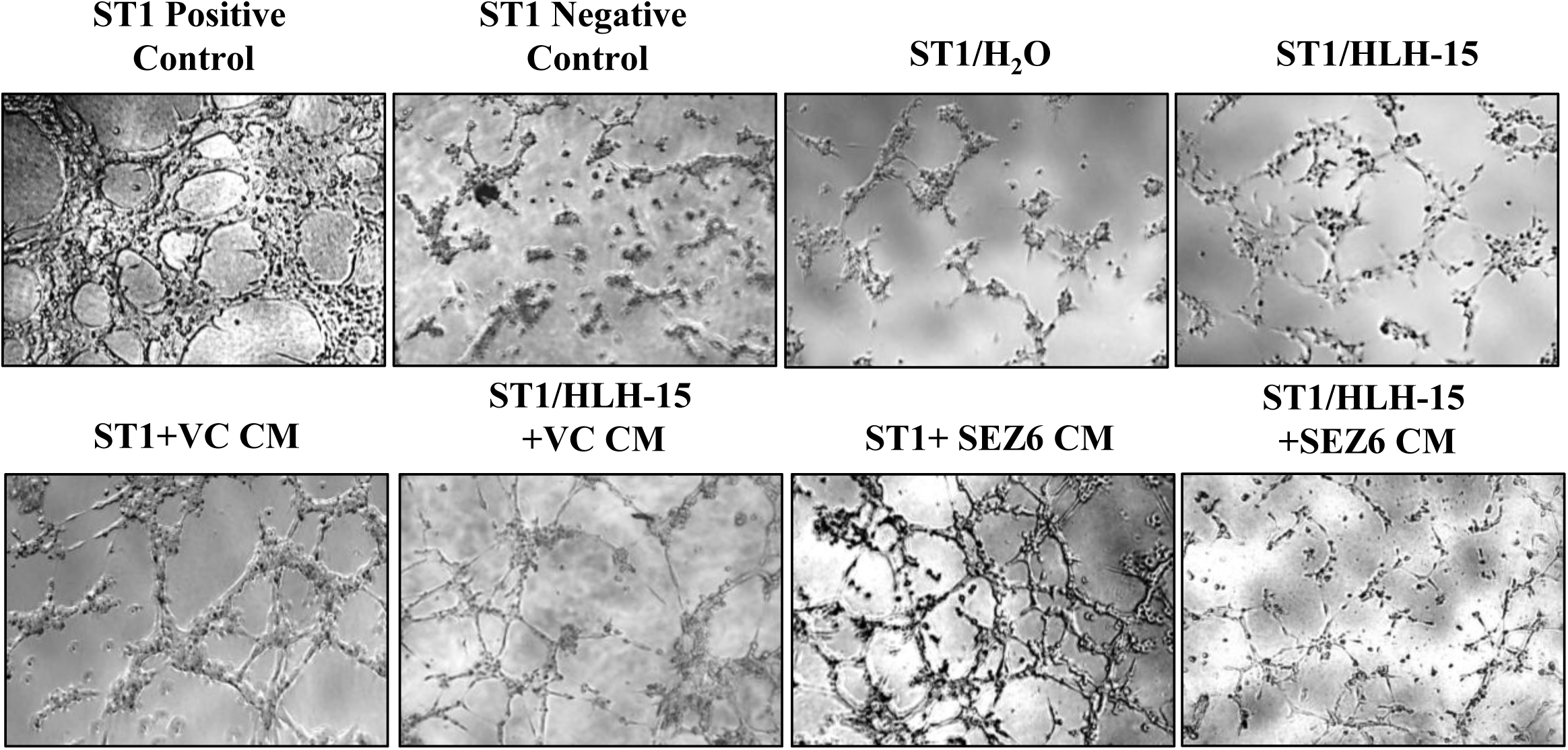

**Supplementary Figure 14.**
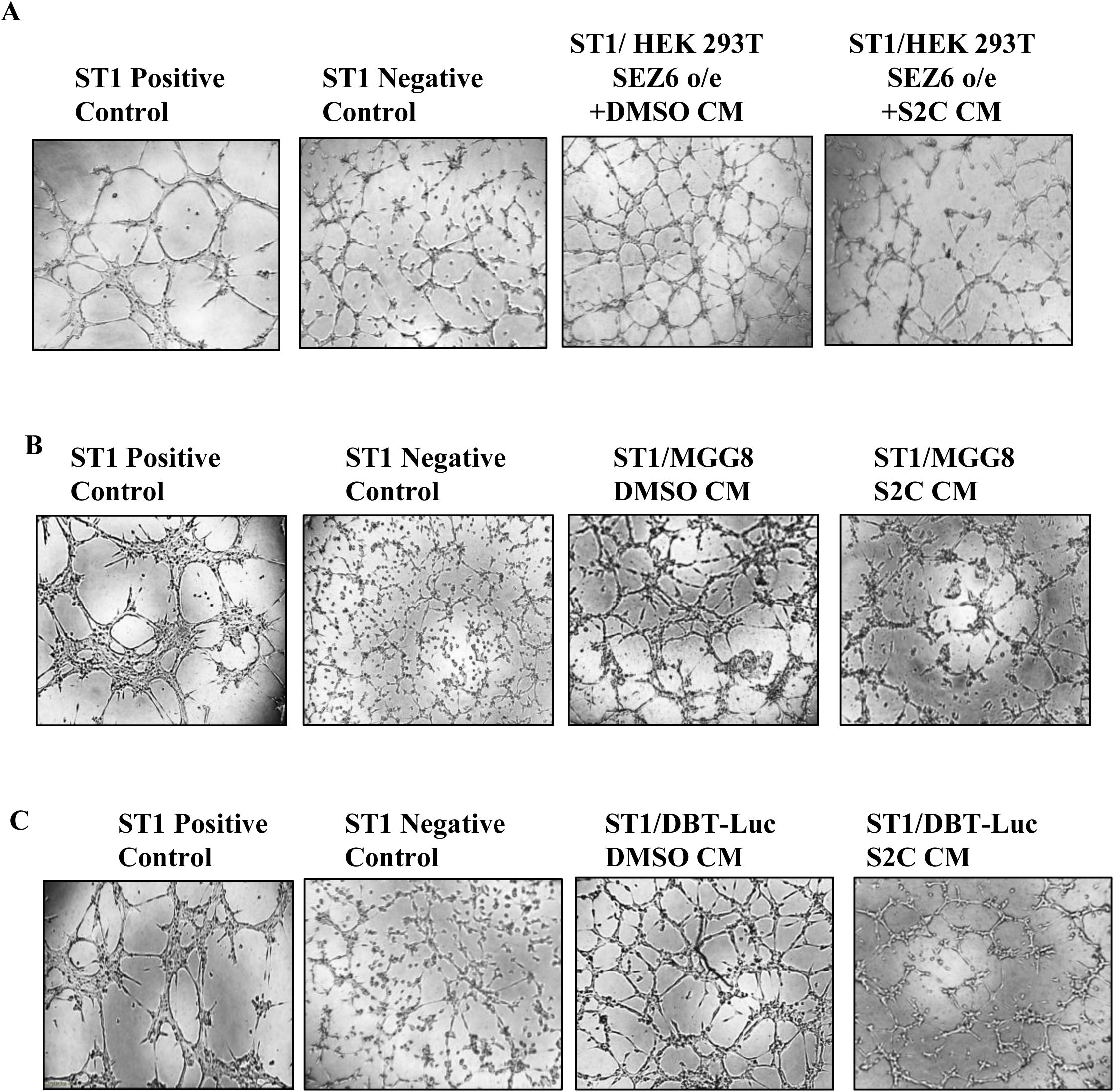

