## Supplementary Information for "Seizure-related gene 6 (SEZ6) encodes a Cancer stem cell-specific proangiogenic molecule: A novel glioma therapeutic target"

### Identifying Sushi-3 domain as the interacting module and mapping key residues involved in TGFβ RII ECD binding

Sushi domains, also known as complement control protein (CCP) modules or short consensus repeats (SCRs), are approximately 60-amino acid modular units arranged in tandem repeats, and are found commonly in complement regulatory and cell-adhesion proteins **(Lehtinen et al., 2004)**. Multiple sequence alignment of the Sushi-3, Sushi-4, and Sushi-5 domains confirmed the presence of all sequence hallmarks of the Sushi domain family **(Kirkitadze et al., 2001)**, including four conserved cysteines which form two disulfide bonds (C1-C3 and C2-C4) that stabilize the compact β-sandwich fold of Sushi domains, an invariant tryptophan between C3 and C4, and conserved glycines and prolines **(Fig. 5A)**. Although pairwise sequence identity was limited to 27-34%, similarity values of 44-47% and conservation of key residues support the preservation of structural features typically maintained across evolutionarily divergent Sushi domains. To delineate the interaction of Sushi-3-4-5 with the TGFβ RII ECD, the three-dimensional structure of the Sushi-3-4-5 region was predicted using AlphaFold2 **(Jumper et al., 2021)**. The top-ranked model exhibited a high average pLDDT (Predicted Local Distance Difference Test) score of 90.94 **(Fig. S5A)** and low PAE (Predicted Aligned Error) values within and between domains **(Fig. S5B)**, indicating high confidence in both the individual domain folds and their relative spatial organization. Ramachandran statistics demonstrated that 92.1% and 7.9% of residues occupied the most favored and additionally allowed regions, respectively, supporting high stereochemical quality of the predicted Sushi-3-4-5 structure **(Fig. S5C)**. Blind protein-protein docking of the top-ranked Sushi-3-4-5 model with the TGFβ RII ECD (PDB ID: 1KTZ) using five independent algorithms (ClusPro2.0 **(Kozakov et al., 2017)**, GRAMM **(Singh et al., 2024)**, HDOCK **(Yan et al., 2020)**, InterEvDock3 **(Quinot et al, 2021)**, and LZerD **(Christoffer et al., 2021)** consistently positioned Sushi-3 at the receptor-binding interface **(Fig. S5D-H)**. Superposition of the top-ranked docked complexes revealed substantial agreement in binding interfaces across methods, identifying Sushi-3 as the most likely TGFβ RII-interacting domain **(Fig. 5B)**.

To investigate the structural basis of the apparent preference of Sushi-3 for TGFβ RII binding, Sushi-3, Sushi-4, and Sushi-5 were modelled individually using AlphaFold2 and subsequently analyzed in complex with TGFβ RII. The predicted structures were of high quality, with average pLDDT scores of 88.79-95.98 **(Fig. 5C)**, low PAE values **(Fig. 5D)**, and Ramachandran statistics showing 89.4-94.2% and 5.8-10.6% of residues in the most favored and additionally allowed regions, respectively, and no residues in disallowed conformations **(Fig. S6A-C)**. Consistent with the conserved Sushi fold, the domains exhibited high structural similarity, with pairwise $C_\alpha$ RMSD values of 0.66-0.78 Å **(Fig. 5E)**. To investigate their interactions with TGFβ RII, each domain-receptor complex was subjected to 1000 ns all-atom molecular dynamics (MD) simulations. While Sushi-3 and Sushi-5 remained stably associated with TGFβ RII throughout the simulations, Sushi-4 dissociated from the receptor. Binding free-energy calculations using the MM-PBSA (Molecular Mechanics Poisson-Boltzmann Surface Area) approach identified Sushi-3 as the strongest binder (-33.01 kcal/mol), followed by Sushi-5 (-30.43 kcal/mol) and Sushi-4 (-0.36 kcal/mol) **(Fig. 5F)**. Sushi-3 also formed a substantially larger interaction interface than Sushi-4 and Sushi-5 **(Fig. 5G),** and exhibited the highest persistence of non-covalent contacts throughout the MD trajectory **(Fig. S7-S9)**. Together with the docking analyses, these results identify Sushi-3 as the most likely mediator of TGFβ RII binding. Notably, interfacial residues in the Sushi-3-TGFβ RII complex, involved in hydrogen bonds (S40-E51 and S41-E51), salt bridges (E53-R42 and E53-K81), and hydrophobic interactions (V26-F11 and V42-V9), remained within 6 Å for 88-100% of the simulation time, highlighting a stable interaction network **(Fig. S10 and Fig. 5J)**.

**De novo peptide design and analysis**

From the MD simulation of the Sushi-3-TGFβ RII complex, we identified six residues on the TGFβ RII ECD - V9, F11, R42, E51, Y61, and K81, that consistently engaged with Sushi-3 throughout the trajectory. Based on this footprint, we designed *de novo* peptide binders that would compete for the same surface patch on TGFβ RII comprising the six hotspot residues, and competitively block SEZ6 from engaging with the same interface. The design rationale was based on the BindCraft pipeline **(Pacesa et al., 2025)**, which we specifically adapted and optimized for short peptide design. We implemented BindCraft's three-stage pipeline involving AlphaFold2-based hallucination via backpropagation to simultaneously optimise binder sequence and structure against the target, followed by $MPNN_{sol}$ based sequence refinement of non-interface residues to improve solubility, and final re-evaluation using AlphaFold2 monomer and Rosetta-based metrics to select high-confidence designs. Binders were designed in two length ranges (4-25 and 26-40 residues) to evaluate the influence of binder size on target engagement. Cysteines were excluded to avoid undesired disulfide formation, and design parameters were further constrained to favor predominantly α-helical peptides (>90% helical content) to promote structural stability. Using this strategy, we generated 500 designs for the longer-peptide-length bucket (26-40 residues). The resulting peptide designs predominantly adopted helix-loop-helix architectures (n = 337), while the remaining designs consisted of single α-helical structures (n = 163). For each design, we quantitatively assessed 13 design parameters encompassing six AlphaFold-derived confidence metrics and seven Rosetta-based physicochemical descriptors. Designs were graded by the number of parameter violations, with all peptides failing any AlphaFold2 confidence metric excluded, as these are critical indicators of structural integrity and interface quality. All peptides were visually inspected and classified as on-site or off-site binders based on TGFβ RII interface localization. Of 500 designs, 259 were on-site and 241 off-site binders **(Fig. S11A)**. Among 259 on-site peptides, 149 satisfied all six AlphaFold2-derived quality parameters, of which 49 occupied the target hotspot residues wholly over the binding interface. Within this subset, parameter violations ranged from 1-6, with 2 peptides violating 1 parameter, 18 violating 2, 17 violating 3, 10 violating 4, and 1 each violating 5-6 parameters **(Fig. 6A)**. All-atom MD simulations of 49 peptides complexed with TGFβ RII identified 16 stably bound peptides **(Fig. S11B)**. Subsequent simulations of these 16 peptides in isolation revealed that 15 retained their native secondary structures throughout, demonstrating strong intrinsic folding propensities independent of receptor context. These 15 peptides exhibited binding free energies of -30 to -51 kcal/mol, indicating strong TGFβ RII binding. All 15 peptides adopted helix-loop-helix architectures, a common motif in protein-protein interfaces **(Murre et al., 2019)**. Six of these exhibited coiled-coil heptad repeats, indicating stable hydrophobic cores that enhance structural rigidity **(Fig. S11C)**. Among the short peptides (4-25 residues), only 1 helix-loop-helix design remained stably bound in MD, but it lost secondary structure in isolation. Due to instability of short helix-loop-helix designs, further analysis focused on longer peptides. Among all peptides exhibiting stable secondary structure and coiled-coil architecture, the three most favorable by binding free energy (HLH-4, HLH-13, HLH-15) were chosen for experimental validation. Circular dichroism spectroscopy confirmed that all three peptides adopted α-helical secondary structures in solution, correlating with *in silico* predictions **(Fig. S12A-C)**. Isothermal titration calorimetry confirmed HLH-15 as the strongest binder with a $K_d$ of 45.6 nM, consistent with its lowest MM-PBSA based average binding free energy of -46.42 kcal/mol, among the three candidates **(Fig. S13A-C)**. Inspection of the key non-covalent interactions between HLH-15 and TGFβ RII identified consistent contributions from hydrogen bonding (R11-E84, H15-Y61, and W36-E84), salt-bridge interactions (R19-E51 and R25-E35), hydrophobic packing (V26-L59 and W36-F86), and other interactions (R19-V53, W36-H62, W36-K81, F39-P82, and F39-G83) across the interface **(Fig. 6C)**.

### Compound S2C characterization via high-resolution mass spectrometry and NMR spectroscopy

#### Step – 1

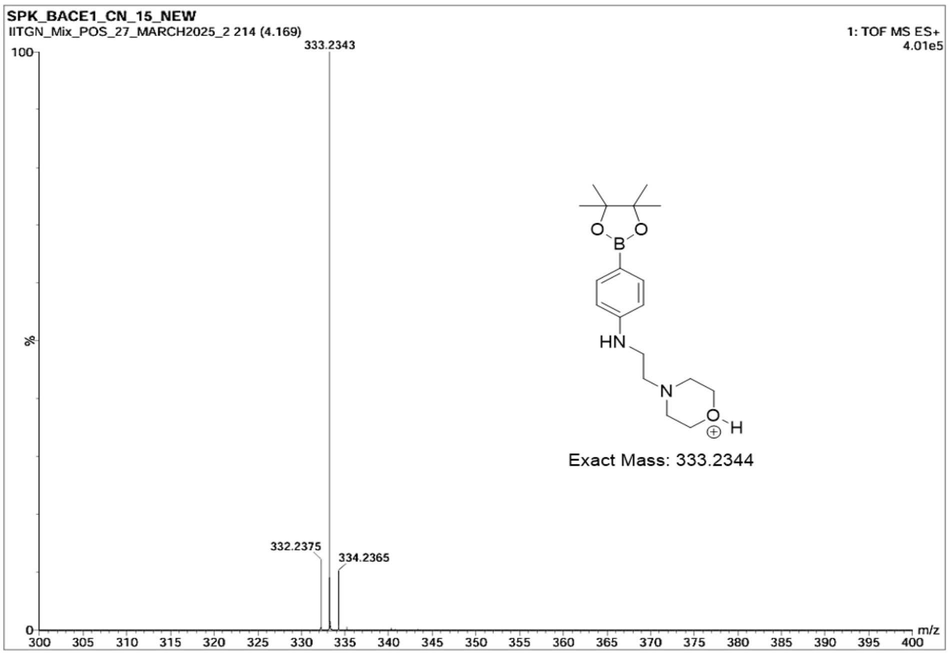

**Step – 2**

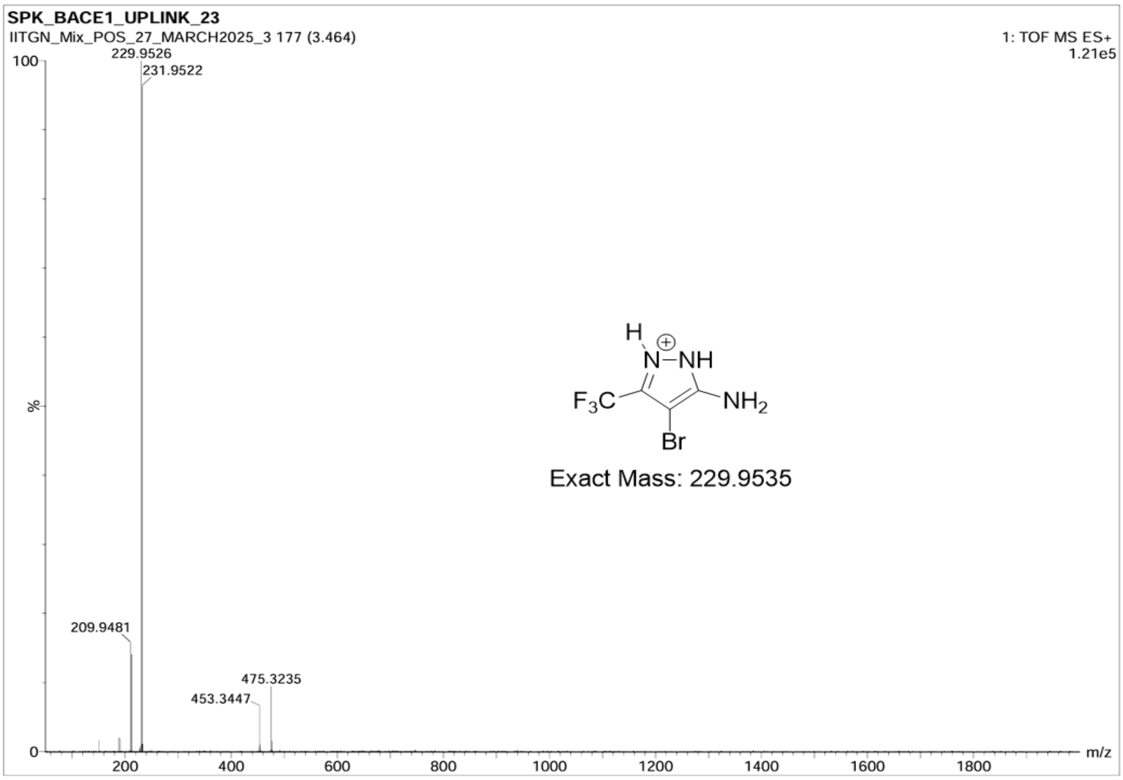
SPK_BACE1_UPLINK_23
IITGN_Mix_POS_27_MARCH2025_3 177 (3.464)
1: TOF MS ES+
1.21e5
100
229.9526
231.9522
%
209.9481
475.3235
453.3447
0
200
400
600
800
1000
1200
1400
1600
1800
m/z
Exact Mass: 229.9535

**Step – 3**

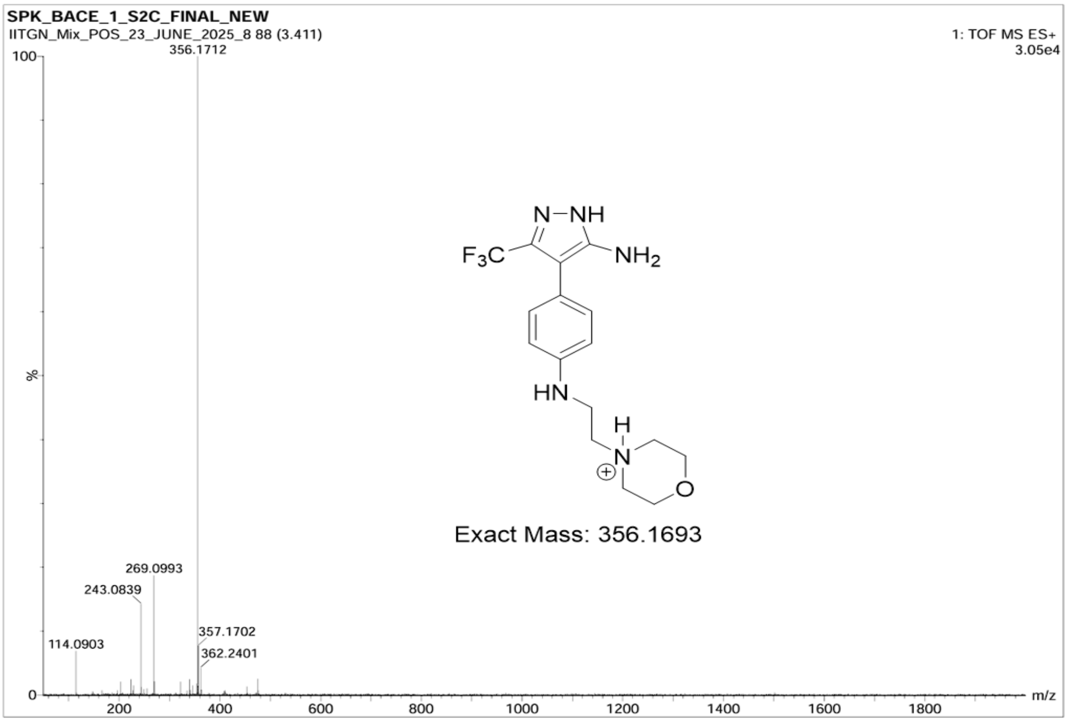
SPK_BACE_1_S2C_FINAL_NEW
IITGN_Mix_POS_23_JUNE_2025_8 88 (3.411)
1: TOF MS ES+
3.05e4
100
356.1712
%
269.0993
243.0839
357.1702
114.0903
362.2401
0
200
400
600
800
1000
1200
1400
1600
1800
m/z
Exact Mass: 356.1693

### NMR characterization of Step-1, Step-2 and Step-3 compounds

**(i) N-(2-morpholinoethyl)-4-(4,4,5,5-tetramethyl-1,3,2-dioxaborolan-2-yl)aniline:** $^{1}$H NMR (500 MHz, DMSO-$d_6$)

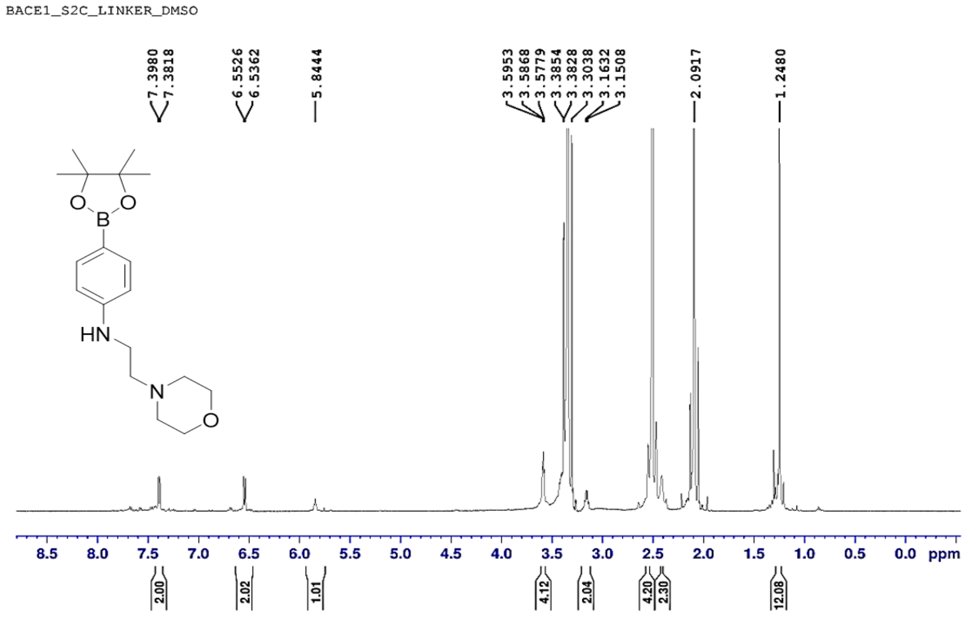


**(ii) 4-bromo-3-(trifluoromethyl)-1H-pyrazol-5-amine:** $^{1}$H NMR (500 MHz, DMSO-$d_6$)

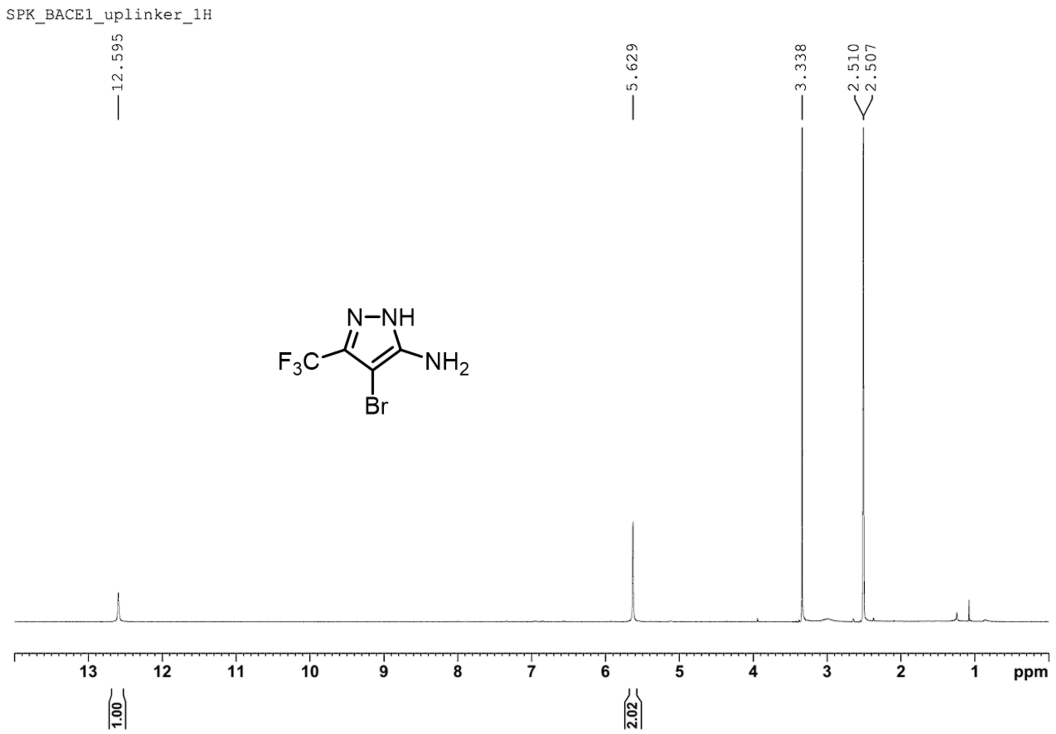

**(iii)** **4-(4-((2-morpholinoethyl)amino)phenyl)-3-(trifluoromethyl)-1H-pyrazol-5-amine:** $^{1}$H NMR (500 MHz, DMSO-$d_6$)

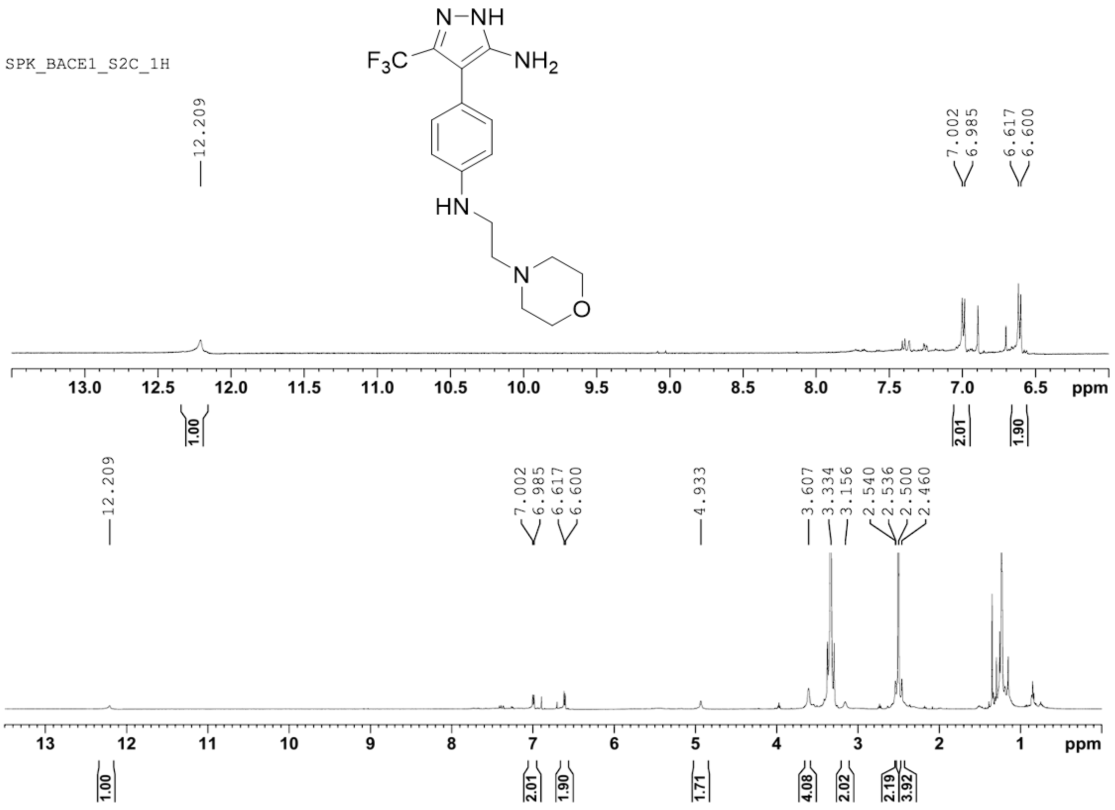


**4-(4-**

**(iv)** **((2-morpholinoethyl)amino)phenyl)-3-(trifluoromethyl)-1H-pyrazol-5-amine:** $^{1}$H NMR (500 MHz, $CDCl_3$)

### Supplementary Figure legends

#### Supplementary Figure 1

**(A)** Bubble plot showing significant enrichment of several angiogenesis-related terms when the differentially expressed genes in the publicly available transcriptome (GSE54791, Suva et al, 2013) of Glioma Stem-like Cells (GSCs) over Differentiated Glioma Cells (DGCs) were subjected to SignalP analysis followed by Gene Ontology (GO) analysis. Gene Count indicates the number of genes from the input list involved in the corresponding GO term. p-Value <0.05 is considered significant.

**(B)** RT-qPCR showing the upregulation of most of the thirteen genes of the TCF/LEF gene set in MGG8-GSCs over MGG8-DGCs. p-Value <0.05 was considered significant with *, **, and *** representing p-values <0.05, 0.01, and 0.001, respectively. ns, non-significant.

**(C)** RT-qPCR showing the downregulation of the thirteen genes of the TCF/LEF gene set upon shRNA-mediated knockdown in MGG8-GSCs as compared to MGG8-GSCs transduced with shNT. p-Value <0.05 was considered significant with *, **, and *** representing p-values <0.05, 0.01, and 0.001, respectively. ns, non-significant.

**(D)** Representative images of light microscopy showing the impact on neurosphere formation upon knockdown of each of the thirteen genes of the TCF/LEF gene set in MGG8-GSCs compared to MGG8-GSCs transduced with shNT. Magnification 4X, Scale 200 µm.

**(E)** Limiting dilution assay showing the impact on neurosphere formation upon knockdown of each of the thirteen genes of the TCF/LEF gene in MGG8-GSCs compared to MGG8-GSCs transduced with shNT.

**(F)** Representative images of the in vitro network formation by ST1 cells upon treatment with CM collected from MGG8-GSCs silenced for each of the thirteen genes of the TCF/LEF gene set, as compared to MGG8-GSCs transduced with shNT. The positive control is for ST1 cells plated with serum-supplemented complete endothelial cell media, the negative control is for

ST1 cells plated in incomplete medium, and the NBM control is for ST1 cells plated in incomplete neurobasal media (NBM). Magnification 4X, Scale 200 µm.

**(G)** Representative images of the in vitro network formation by ST1 cells upon treatment with CM collected from MGG8-shNT and MGG8-shSEZ6, followed by rescue with rhSEZ6 in shSEZ6 CM and inhibition by SEZ6-specific antibody in shNT CM. The positive control is for ST1 cells plated with serum-supplemented complete endothelial cell media, the negative control is for ST1 cells plated in incomplete medium, and the NBM control is for ST1 cells plated in incomplete neurobasal media (NBM). Magnification 4X, Scale 200 µm.

**(H)** RT-qPCR showing upregulation of the SEZ6 transcript in four GSC lines (MGG4, MGG6, MGG8, MGG23) compared with their corresponding DGC counterparts. p-Value <0.05 was considered significant with *, **, and *** representing p-values <0.05, 0.01, and 0.001, respectively. ns, non-significant.

**(I)** Western blot showing increased expression of intracellular (left panel) and extracellular (right panel) SEZ6 protein level in MGG4-GSC and MGG8-GSC over MGG4-DGC and MGG8-DGC, respectively. For the intracellular western blot, GAPDH is used as the loading control, and a decrease in SOX2 levels in DGCs relative to GSCs is used as a marker of completed differentiation. For the extracellular western blot, the Ponceau-stained membrane shows equal loading of the conditioned medium (right panel).

**Supplementary Figure 2**

**(A)** Representative images of the in vitro network by b.End.3 cells, murine endothelial cells derived from the brain tissue of a mouse with an endothelioma, upon treatment with recombinant SEZ6 (rhSEZ6). The ST1 positive control is ST1 cells plated in serum-supplemented complete endothelial cell media, and the negative control is ST1 cells plated in incomplete medium. The b.End.3 positive control is b.End.3 cells plated in serum-

complete endothelial cell media, and the negative control is b.End.3 cells plated in incomplete medium. Quantification of the total number of networks formed (lower panel) shows a significant increase in network formation by b.End.3 cells upon SEZ6 treatment compared with BSA treatment. p-Value <0.05 was considered significant with *, **, and *** representing p-values <0.05, 0.01, and 0.001, respectively. ns, non-significant.

**(B)** Representative images of the in vitro network by HBMEC cells, immortalized human brain microvascular endothelial cells, upon treatment with recombinant SEZ6 (rhSEZ6). The ST1 positive control is ST1 cells plated in serum-supplemented complete endothelial cell media, and the negative control is ST1 cells plated in incomplete medium. The HBMEC positive control is HBMEC cells plated in serum-supplemented complete endothelial cell media, and the negative control is HBMEC cells plated in incomplete medium. Quantification of the total number of networks formed (lower panel) shows a significant increase in network formation by HBMECs upon SEZ6 treatment compared with BSA treatment. p-Value <0.05 was considered significant with *, **, and *** representing p-values <0.05, 0.01, and 0.001, respectively. ns, non-significant.

**(C)** Representative images of the trans-well migration assay, upon treatment of ST1 cells with rhSEZ6 and BSA. The bar graph (right panel) shows a significant increase in migrated cells upon treatment with rhSEZ6 compared with BSA. p-Value <0.05 was considered significant with *, **, and *** representing p-values <0.05, 0.01, and 0.001, respectively. ns, non-significant.

**(D)** Representative images of the Matrigel-coated trans-well invasion assay, upon treatment of ST1 cells with rhSEZ6 and BSA. The bar graph (right panel) shows a significant increase in cell invasion upon rhSEZ6 treatment compared with BSA. p-Value <0.05 was considered significant with *, **, and *** representing p-values <0.05, 0.01, and 0.001, respectively. ns, non-significant.

**(E)** MTT assay showing a significant increase in the proliferation of ST1 cells upon rhSEZ6 treatment for 72 hours as compared to BSA. p-Value <0.05 was considered significant with *, **, and *** representing p-values <0.05, 0.01, and 0.001, respectively. ns, non-significant.

**Supplementary Figure 3**

**(A)** Membranes from the Human Proteome Profiler Angiogenesis Array show the differentially expressed proteins in the ST1 cells upon treatment with CM derived from MGG8/shNT compared to MGG8/shSEZ6 cells.

**(B)** Representative images of in vitro network formation assay in ST1/shNT cells and ST1/shIL8, followed by treatment with rhSEZ6 and rhIL8 in both these conditions. The positive control is ST1 cells plated in serum-supplemented complete endothelial cell medium. The negative control is for ST1 cells plated in incomplete medium. Magnification 4X, Scale 200 μm.

**(C)** Representative images of in vitro network formation assay ST1 cells upon pretreatment with TGFβ-RI inhibitor, SB341542 (10 μM), followed by addition of CM derived from MGG8/shNT. The positive control is ST1 cells plated in serum-supplemented complete endothelial cell medium. The negative control is for ST1 cells plated in incomplete medium. The NBM control is for ST1 cells plated in incomplete Neurobasal Media (NBM). Magnification 4X, Scale 200 μm.

**Supplementary Figure 4**

Representative images of the in-vitro angiogenesis assay showing the network formation by ST1 cells upon treatment with CM containing the shed forms WT-SEZ6 and the shed forms of the indicated deletion mutants of SEZ6.

**Supplementary Figure 5**

**(A)** Structure of the stretch of Sushi 3-4-5 as predicted and modeled by AlphaFold2.

**(B)** Residue-wise pLDDT scores for the AlphaFold2-predicted Sushi-3-4-5 structure reveal consistently high confidence in local structural predictions across all residue positions.

**(C)** PAE analysis of the Sushi-3-4-5 structure reveals low positional error (blue spots) across residue pairs, confirming high accuracy in relative residue positioning.

**(D)** Ramachandran plot of the AlphaFold2-predicted Sushi-3-4-5 structure generated using PROCHECK illustrates phi-psi dihedral angles for all residues, with the majority of residues falling within most favored regions (red) and a few residues occupying additional allowed regions (brown), validating the stereochemical quality of the predicted model.

**E-I)** Consensus docking results from five independent methods (ClusPro2.0, GRAMM, HDOCK, InterEvDock3, and LZerD) identify interfacial residue contacts (≤ 8 Å) between Sushi-3-4-5 and TGFβ RII, with color-coding indicating the rank of the corresponding docked complex. The predominant localization of interfacial residues to Sushi-3 across all methods establishes it as the primary binding domain.

**Supplementary Figure 6**

Ramachandran plots of the AlphaFold2-predicted structures of **A)** Sushi-3, **B)** Sushi-4, and **C)** Sushi-5 generated using PROCHECK, illustrating phi-psi dihedral angles for all residues, reveal predominant localization of residues in the most favored regions (red), with minimal occupancy in additional allowed regions (brown), validating the high stereochemical quality of all three predicted models.

**Supplementary Figure 7**

PyContact-derived interaction maps of the Sushi-3-TGFβ RII complex from 1000 ns MD simulation, stratified into **(A)** hydrogen bond contact maps showing frequency and persistence, **(B)** electrostatic interaction networks revealing salt bridge contributions to binding, and **(C)** hydrophobic interaction landscape at the interface, with color-coding distinguishing residue pair involvement (backbone-backbone in green, sidechain-backbone in red, sidechain-sidechain in blue) and color intensity denoting interaction strength.

**Supplementary Figure 8**

PyContact-derived interaction maps of the Sushi-4-TGFβ RII complex from 1000 ns MD simulation, stratified into **(A)** hydrogen bond contact maps showing frequency and persistence, **(B)** electrostatic interaction networks revealing salt bridge contributions to binding, and **(C)** hydrophobic interaction landscape at the interface, with color-coding distinguishing residue pair involvement (sidechain-backbone in red, and sidechain-sidechain in blue) and color intensity denoting interaction strength.

**Supplementary Figure 9**

PyContact-derived interaction maps of the Sushi-5-TGFβ RII complex from 1000 ns MD simulation, stratified into **(A)** hydrogen bond contact maps showing frequency and persistence, **(B)** electrostatic interaction networks revealing salt bridge contributions to binding, and **(C)** hydrophobic interaction landscape at the interface, with color-coding distinguishing residue pair involvement (backbone-backbone in green, sidechain-backbone in red, sidechain-sidechain in blue) and color intensity denoting interaction strength.

**Supplementary Figure 10**

**A-F)** Time-resolved interaction dynamics of six key Sushi-3-TGFβ RII residue pairs from 1000 ns MD simulations analyzed using MDAnalysis. Residue pairs were selected based on mean interaction strength exceeding 50% of the maximum observed strength for their respective interaction class (hydrogen bond, salt bridge, or hydrophobic interaction), highlighting the most persistent interfacial contacts across the simulation. For each pair, the upper panel displays the distance between nearest heavy atoms throughout the trajectory, with percentage occupancy within a 6 Å threshold annotated. The lower panel shows the corresponding interaction strength profile derived from PyContact analysis, with color-coding indicating the involvement of interaction residue pairs (sidechain-backbone in red and sidechain-sidechain in blue) and color intensity denoting interaction strength.

**Supplementary Figure 11**

**(A)** UpSet plot summarizing design parameter compliance across 500 designed peptides. The matrix (lower panel) displays intersections of 13 design parameters (6 AlphaFold2-derived and 7 Rosetta-based metrics), with filled circles indicating satisfied parameters. Matrix columns are color-coded by number of Rosetta-based metric violations (1 violation in dark green, 2 in yellow, 3 in red, 4 in cyan, 5 in violet, and 6 in light green). The bar chart (upper panel) quantifies peptide counts for each parameter combination, with bars color-coded to distinguish on-site binders (blue) from off-site binders (maroon). The threshold values for the parameters are as follows: pLDDT≥0.8; pTM≥0.55; pAE≤0.35; ipLDDT≥0.8; ipTM≥0.50; iPAE≤0.35; Surface hydrophobicity≤0.35; Shape complementarity≥0.6; PackStat≥0.65; Interface residues≥7; Interface H-bonds≥3; Interface unsaturated H-bonds≤4; RMSD≤3.5

**(B)** Binding dynamics of 49 selected peptides over 1000 ns MD simulations stratified into four consecutive time intervals. Stacked bars show the count of peptides remaining bound, dissociated, or exhibiting binding interface changes within each time interval. Notably, only 16 peptides remained stably bound to the TGFβ RII interface throughout the entire 1000 ns trajectory.

**(C)** Summary of the 16 stably bound peptides showing peptide length, MM-PBSA binding free energy (kcal/mol) averaged over 1000 ns MD simulations, presence of coiled-coil architecture identified using Socket2, and secondary structure stability assessed via DSSP analysis across 1000 ns MD trajectories.

**Supplementary Figure 12**

**(A)** Secondary structure representation of HLH-4 peptide showing intact alpha-helical structure upon 1000 ns unbound MD simulation.

**(B)** DSSP profiles of HLH-4 from 1000 ns unbound MD simulation, confirming stable α-helical secondary structure independent of TGFβ RII interaction.

**(C)** Circular Dichroism (CD) profile of HLH-4 characteristic of α-helices, validating stable secondary structural integrity across the given temperature gradient (20°C to 95°C).

**(D)** Secondary structure representation of HLH-13 peptide showing intact alpha-helical structure upon 1000 ns unbound MD simulation.

**(E)** DSSP profiles of HLH-13 from 1000 ns unbound MD simulation, confirming stable α-helical secondary structure independent of TGFβ RII interaction.

**(F)** Circular Dichroism (CD) profile of HLH-13 characteristic of α-helices, validating stable secondary structural integrity across the given temperature gradient (20°C to 95°C).

**(G)** Binding affinity assessment of HLH-4 to TGFβ RII-ECD by MM-PBSA-derived binding free energies (kcal/mol) computed across 1000 ns MD simulations.

**(H)** Average structure of the HLH-4 bound to TGFβ RII-ECD during the 1000 ns MD simulation.
**(I)** Determination of experimental dissociation constant ($K_d$) value of HLH-4 with bacterially purified TGFβ RII-ECD using isothermal titration calorimetry (ITC).
**(G)** Binding affinity assessment of HLH-13 to TGFβ RII-ECD by MM-PBSA-derived binding free energies (kcal/mol) computed across 1000 ns MD simulations.
**(H)** Average structure of the HLH-13 bound to TGFβ RII-ECD during the 1000 ns MD simulation.
**(I)** Determination of experimental dissociation constant ($K_d$) value of HLH-13 with bacterially purified TGFβ RII-ECD using isothermal titration calorimetry (ITC).

**Supplementary Figure 13**

Representative images of the angiogenesis assay showing the impact of pre-treatment of ST1 cells with HLH-15, followed by subsequent treatment of VC CM and SEZ6 CM on the in-vitro network-forming ability of ST1 cells.

**Supplementary Figure 14**

RMSF profile of the Cα atoms of BACE1 in complex with S2C over the 1000 ns molecular dynamics simulation. The plot shows residue-wise fluctuations, with lower RMSF values indicating stable regions and higher RMSF values indicating flexible loop and terminal regions.

**Supplementary Figure 15**

**(A)** Representative images of the angiogenesis assay showing the impact of S2C-treated HEK 293T-derived CM on the in vitro network formation by ST1 cells, compared to DMSO-treated HEK 293T-derived CM.

(**B**) Representative images of the angiogenesis assay showing the impact of S2C-treated MGG8-derived CM on the in vitro network formation by ST1 cells, compared to DMSO-treated MGG8-derived CM.

(**C**) Representative images of the angiogenesis assay showing the impact of S2C-treated DBT Luc-derived CM on the in vitro network formation by ST1 cells, compared to DMSO-treated DBT Luc-derived CM.
